# A combinatorially complete epistatic fitness landscape in an enzyme active site

**DOI:** 10.1101/2024.06.23.600144

**Authors:** Kadina E. Johnston, Patrick J. Almhjell, Ella J. Watkins-Dulaney, Grace Liu, Nicholas J. Porter, Jason Yang, Frances H. Arnold

**Author notes:** Corresponding Author: Frances H. Arnold **Email:**. Merck & Co., Inc., South San Francisco, CA 94080. Department of Biochemistry, Stanford University, Stanford, CA 94305. Harvard Medical School, Boston, MA 02115. Codexis Inc., Redwood City, CA 94063. These authors contributed equally to this work and are listed alphabetically (Patrick J. Almhjell, Ella J. Watkins-Dulaney). **Author Contributions:** K.E.J., E.J.W.-D., and F.H.A. designed research. K.E.J. and N.J.P. collected data. K.E.J., P.J.A., G.L., and J.Y. wrote software and analyzed data. K.E.J. wrote the manuscript. K.E.J., E.J.W.-D., P.J.A., G.L., N.J.P., J.Y., and F.H.A. revised and edited the manuscript. **Competing Interest Statement:** The authors declare no competing financial interest. **Classification:** Biological Sciences, Applied Biological Sciences.

## Abstract

Protein engineering often targets binding pockets or active sites which are enriched in epistasis— non-additive interactions between amino acid substitutions—and where the combined effects of multiple single substitutions are difficult to predict. Few existing sequence-fitness datasets capture epistasis at large scale, especially for enzyme catalysis, limiting the development and assessment of model-guided enzyme engineering approaches. We present here a combinatorially complete, 160,000-variant fitness landscape across four residues in the active site of an enzyme. Assaying the native reaction of a thermostable β-subunit of tryptophan synthase (TrpB) in a non-native environment yielded a landscape characterized by significant epistasis and many local optima. These effects prevent simulated directed evolution approaches from efficiently reaching the global optimum. There is nonetheless wide variability in the effectiveness of different directed evolution approaches, which together provide experimental benchmarks for computational and machine learning workflows. The most-fit TrpB variants contain a substitution that is nearly absent in natural TrpB sequences—a result that conservation-based predictions would not capture. Thus, although fitness prediction using evolutionary data can enrich in more-active variants, these approaches struggle to identify and differentiate among the most-active variants, even for this near-native function. Overall, this work presents a new, large-scale testing ground for model-guided enzyme engineering and suggests that efficient navigation of epistatic fitness landscapes can be improved by advances in both machine learning and physical modeling.

**Significance statement:** Predictive models for protein engineering seek to capture the relationship between protein sequence and function. While many methods and datasets exist for predicting the effects of single substitutions across a range of protein functions, fewer capture interactions among substitutions, which are much more difficult to predict. Even fewer do this comprehensively for a catalytic function. To provide a testbed for evaluating predictive models for enzyme engineering, we constructed and analyzed a 160,000-member enzyme sequence-fitness dataset at four interacting residues near the active site of tryptophan synthase, capturing significant non-additive effects of substitutions on catalytic function. It is necessary to predict and understand such interactions in order to efficiently traverse evolutionary landscapes and build machine learning models that accelerate protein engineering.

## Introduction

To engineer a protein for desired properties one must navigate the complex, and largely unknown, relationship between sequence and fitness, where fitness is defined by the engineer rather than evolution. The effects of amino acid substitutions on the multiple properties that determine protein fitness for a given task or application cannot be predicted reliably, especially in enzymes, which must guide substrates through intricate, often multi-step reaction pathways with high efficiency and selectivity. Currently, enzymes are optimized by directed evolution, in which sequential rounds of semi-rational or random mutagenesis and screening are used to accumulate beneficial mutations (1), a process that is time- and resource-intensive. Predictive models for protein fitness, including physical and machine-learning models, will help address urgent needs for new and better enzymes (2). However, the development of such models requires high-quality datasets for training, testing, and comparing new approaches.

Most datasets used for developing and testing predictive models measure the effects of every amino acid substitution across many or even all residues in a protein sequence (3), providing information about single substitutions in the context of a single background sequence. Datasets like these provide no information about the effects of substitutions in different sequence contexts or how different substitutions interact with one another. In many instances, such effects are approximately independent (and therefore “additive”) (4, 5). When substitution effects are mostly independent, combining beneficial substitutions works well to generate improved variants (1), and simple models can often predict the fitnesses of double and even triple mutants from single-site data alone (6). Additivity breaks down, however, when substitutions interact, which can happen when residues are in close proximity or simultaneously interact with cofactors or substrates as in an enzyme active site (7). Because these sites are particularly important for improving fitness, efficient navigation of epistasis is important for many problems in enzyme engineering.

In contrast to our rapidly improving ability to predict protein structure from sequence using machine learning methods (8–11), progress toward fitness prediction, most notably in epistatic regions of proteins, has been slow (12, 13). Combinatorial landscapes in which multiple sites are mutated simultaneously have been an important testing ground for prediction and navigation of epistatic effects (14–16), but few landscapes exist that deeply examine these interactions. There are a variety of landscapes that randomly sample single- and multi-mutants around a given starting sequence (17–20), but only a few that sample multi-mutants comprehensively via extensive mutagenesis at two or three sites simultaneously (21–23). Such landscapes are combinatorially complete, and every variant is characterized within a constrained search space, which enables exact calculation of all epistasis in the landscape (24). These landscapes can be used to test predictions of combined mutational effects and allow for direct comparison of laboratory evolution and machine learning-assisted methods via simulation (14, 15). The few existing 4-site-saturation landscapes measure binding (25, 26), a simpler function than enzyme catalysis which has proven to be more difficult for design (27, 28). We sought to measure a combinatorial fitness landscape of an enzyme active site, furnishing a dataset that would enable examination of epistasis in enzyme catalysis. By comparing this landscape to the well-studied GB1 binding protein landscape (26), we investigate how epistasis constrains evolution on protein fitness landscapes and use these analyses to inform protein engineering approaches and guide the development of new data-driven or physics-based methods for enzyme engineering.

## Results

### Construction of multi-site-saturation combinatorial landscapes using a growth-based assay

For this investigation, we chose the β-subunit of tryptophan synthase (TrpB), which synthesizes L-tryptophan (Trp) from indole and L-serine (Ser). TrpB has been extensively characterized and has a rich sequence diversity due to its presence in all kingdoms of life besides animals. Because Trp is essential for proteome replication and cell growth, a growth-based selection can be developed to measure fitness based on the activity of TrpB (Fig 1A). Such a growth-based selection has been demonstrated in a continuous evolution system, where TrpB variants were selected for their ability to complement Trp auxotrophy in yeast supplied with exogenous indole (29). We decided to monitor fitness in a similar fashion through a pooled-culture enrichment scheme that connects TrpB activity to growth rate. More-active TrpB variants in an *Escherichia coli* Trp auxotroph increase in frequency within the population over the course of selection. From measurements of the relative frequency of mutants through deep sequencing at various timepoints we can calculate a fitness value for each unique variant (Fig 1B) (30).

**Figure 1.**
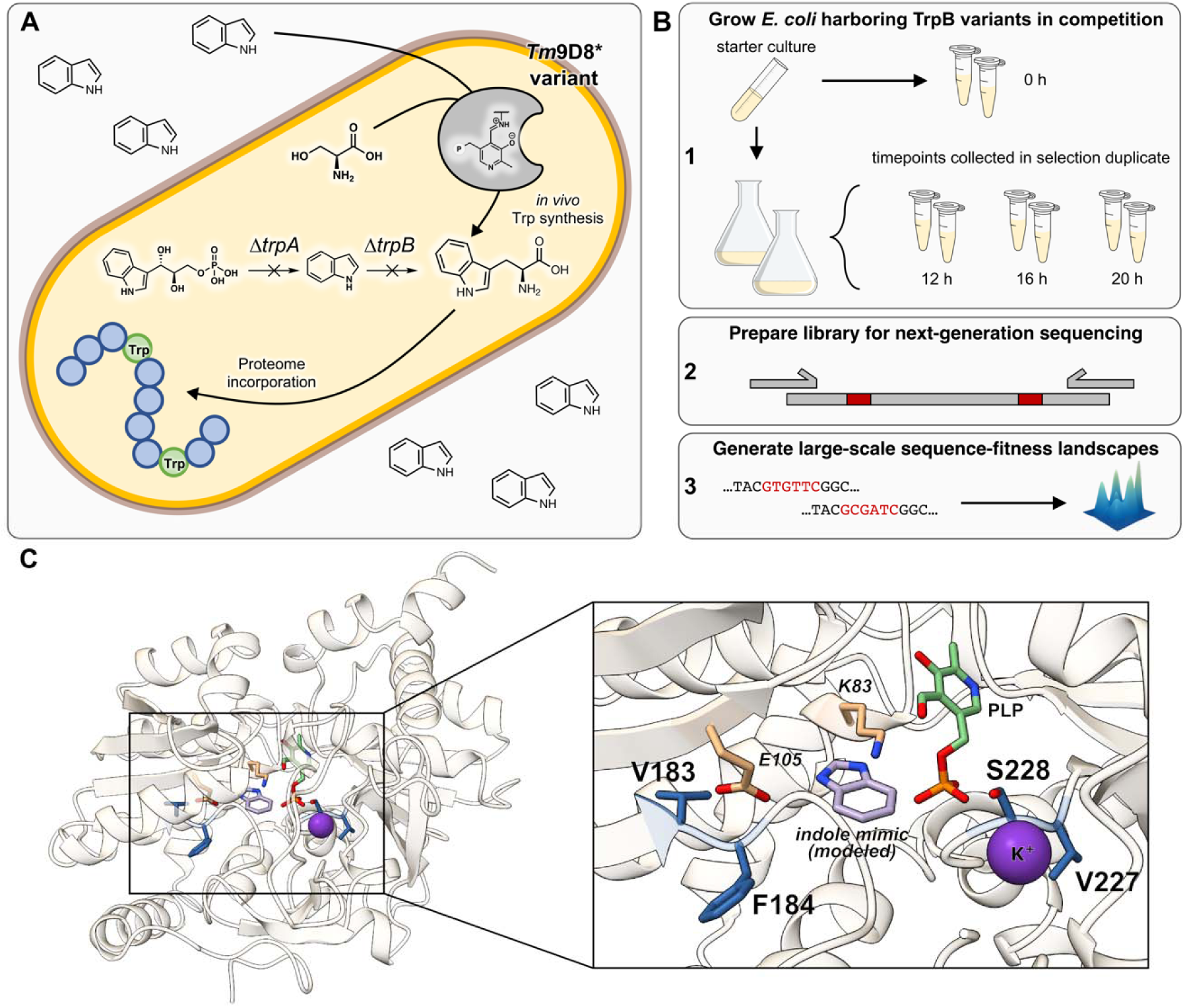
Overview of TrpB-based combinatorial landscapes. **A** An *E. coli* strain with deletions of the *trpA* and *trpB* genes is transformed with plasmid harboring TrpB. When provided with exogenous indole, TrpB expressed from the plasmid produces Trp, enabling proteome and cellular replication at levels that reflect TrpB activity. **B** For each landscape, the *E. coli* Trp auxotroph transformed with a plasmid library is used as a starter culture to inoculate two replicate flasks and to obtain an initial timepoint, T_0_. Samples at different timepoints are collected in duplicate for 36–44 h and prepared for sequencing. **C** The 4-site-saturation library targeted two pairs of positions: 183/184 and 227/228 (blue). The pyridoxal 5’-phosphate (PLP) cofactor of TrpB is colored green, an indole mimic modeled based on PDB ID: 4HPX in lavender (56), and two important catalytic residues are orange, demonstrating the proximity of the selected sites to the catalytic core of TrpB.

We first constructed a strain of Trp-auxotrophic *E. coli* by deleting the *trpA* and *trpB* genes. Deletion of *trpA* avoids potential confounding allosteric interactions between the native *E. coli* α-subunit of tryptophan synthase, TrpA, and the heterologous *Tm*TrpB (31). TrpB variant *Tm*9D8*, derived from the hyperthermophile *Thermotoga maritima,* was selected as the parent enzyme. This variant was evolved to function in the absence of its native allosteric binding partner TrpA (32). *Tm*9D8* is also highly thermostable and exhibits high activity at *E. coli* growth temperatures (33), a useful feature for decoupling a substitution’s contribution to loss of catalytic activity from a loss of stability (4). *Tm*9D8* differs from wildtype *Tm*TrpB by ten amino acid substitutions (P19G, E30G, I69V, K96L, P140L, N167D, I184F, L213P, G228S, and T292S). We verified that *Tm*9D8* could complement Trp auxotrophy with exogenous indole in Trp-dropout media (SI Appendix, Fig S1) and optimized indole concentration and expression conditions (Methods, Preliminary plate-based independent growth assays, SI Appendix, Fig S2).

We next created small libraries to validate the selection protocol against *in vitro* assays. Plate-based independent growth rates and pooled-culture enrichment assays (Fig 1B) were performed for single- (20 possible variants) and double-site- (400 possible variants) saturation libraries using positions near the active site or ones that modulated TrpB activity in previous experiments. Plate-based growth rate assays measured fitness by monitoring cell density of independent cultures over time (Methods, Preliminary plate-based independent growth assays, SI Appendix, Fig S3), whereas the pooled-culture enrichment assay fitness values were obtained by sequencing (Methods, Pooled-culture enrichment assay, SI Appendix, Fig S4). We compared both to the rates of Trp formation collected with *in vitro* lysate-based assays (34), observing a reasonable correlation across each of these activity measurements (SI Appendix, Fig S4–5). These results gave us confidence that the growth assays report on the rate of Trp synthesis (assay details in Methods, Preliminary *in vitro* rate of tryptophan formation assays).

We then designed 3-site-saturation libraries (8,000 possible variants per library), targeting mainly residues in the active site known to impact activity as well as their neighbors. In total, 20 different residue positions were targeted in nine 3-site landscapes, with some overlapping positions (SI Appendix, Fig S6–7). These preliminary tests were designed to evaluate how well the pooled-culture enrichment selection scaled to the larger library size, to identify residues that may participate in epistatic interactions, and to eliminate sites that were completely intolerant to mutation. As expected, we observed a range in the number of substitutions tolerated. Tolerance to substitution also depended on the sequence context in which sites were mutated, indicating epistatic effects (SI Appendix, Fig S8–10).

From among these positions, we chose four residues that displayed epistasis and provided a breadth of activities for a 4-site-saturation (160,000 variant) landscape: 183, 184, 227, and 228 (Fig 1C). From the selection performed on the library at these four sites, 159,129 variants (99.45% of the total library) had sufficient sequencing coverage for quantification in both replicate experiments. Fitness values were calculated for all measured variants as described in Kowalsky *et al.* (35) for multiple timepoints and aggregated into a final fitness score per variant (Methods, Fitness score calculations, SI Appendix, Fig S11–12). Calculated fitness values of overlapping subsets of the 3- and 4-site libraries were highly correlated (SI Appendix, Fig S13). Analysis of the nearly one million unique codon combinations sampled showed that synonymous mutations had minimal impact on fitness, so these sequences were aggregated to report fitness for unique amino acid sequences (SI Appendix, Fig S14–15). The missing 871 fitness values were imputed for downstream analyses (Methods, Fitness score calculations, SI Appendix, Fig S16).

### Investigating the epistasis in combinatorial protein fitness landscapes

The highest-fitness variant of the 4-site landscape—the global optimum—contained substitutions at all four sites (V183A, F184I, V227K, and S228G) with respect to the parent and is referred to hereafter as AIKG. Two substitutions, F184I and S228G, are reversions to wild-type residues in *Tm*TrpB, consistent with the application of an assay designed to capture a near-native function. A third substitution, V183A, incorporates the fourth most-common residue at this position based on a multiple sequence alignment (MSA) for the parent *Tm*9D8* (referred to hereafter as VFVS), found in 10.41% of sequences (SI Appendix, Fig S17). The fourth V227K substitution was surprising: V227K occurs at near-noise levels (0.01%) in natural sequences (SI Appendix, Fig S17) but is clearly beneficial under the assay conditions. Overall, the sequences which were beneficial under these assay conditions appeared more diverse than what might be suspected based on an MSA (SI Appendix, Fig S18). Beyond AIKG, 227K occurred in all ten top variants and in nearly half of the top fifty (SI Appendix, Fig S19). Despite this strong preference, however, 227K is not uniformly beneficial. For example, in the context of the parent sequence of VFVS, it essentially ablates activity and requires the S228G substitution before yielding an improved variant.

For further analysis, we defined an activity threshold as 1.96 standard deviations above the mean fitness of all stop-codon-containing sequences (all of which are expected to be inactive) over both replicates. This threshold is the 97.5^th^ percentile of a normal distribution fit to the fitnesses of the stop-codon-containing sequences. Sequences with fitness values below this threshold were classified as “inactive”, which left 9,783 “active” variants (6.11% of the library) whose activities could be reliably quantified (SI Appendix, Fig S20).

Among the active TrpB and GB1 variants, we quantified pairwise epistasis (Methods, Pairwise epistasis calculations and analyses), including magnitude, reciprocal sign, and sign epistasis for all paths (Fig 2A). Magnitude epistasis, which occurs when the combined effects of two substitutions are in the same direction as expected but are not perfectly additive, is navigable by step-wise or recombination-based DE approaches. Since both additive and magnitude epistasis are navigable by traditional DE approaches, we binned additive mutations along with magnitude effects, which additionally avoids setting a tolerance for experimental deviation to determine truly additive vs. non-additive effects. Sign epistasis occurs when the effect of one of the substitutions changes direction in the context of the other—it is therefore only navigable by step-wise DE approaches if substitutions are made in the correct order, which is not known *a priori*. Finally, reciprocal sign epistasis occurs when the effects of both substitutions change direction when they are made together and is therefore not navigable by DE approaches that use only beneficial substitutions.

**Figure 2.**
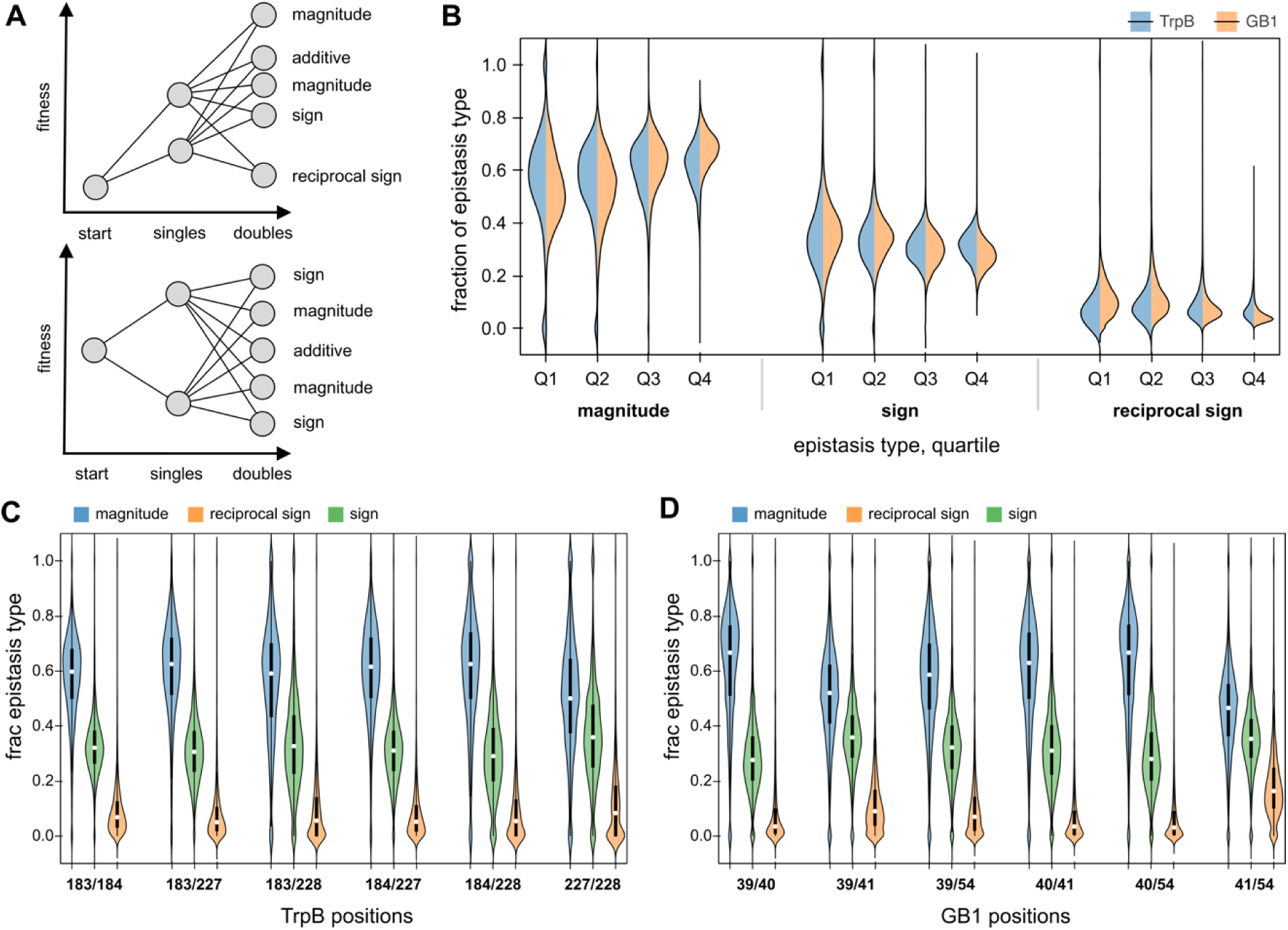
Examining the epistasis in the TrpB and GB1 landscapes. **A** The types of pairwise epistasis where both single substitutions are beneficial (top) or one single substitution is beneficial while the other is deleterious (bottom). **B** Distributions of the three types of pairwise epistasis within the TrpB and GB1 landscapes separated by quartile of the fitness (Q1, Q2, Q3, Q4) of the starting variant, differentiating epistasis prevalence among low-fitness variants (Q1) to high-fitness variants (Q4). **C** Distributions of the three types of pairwise epistasis across all unique pairs of sites for TrpB. **D** Distributions of the three types of pairwise epistasis across all unique pairs of sites for GB1.

We examined the fractions of the different types of pairwise epistasis in the TrpB and GB1 landscapes across all fitness quartiles of the starting variant (Fig 2B). Generally, magnitude epistasis dominates across all fitness quartiles, followed by sign epistasis, and finally by reciprocal sign epistasis. Previous work showed that the mean fraction of each type of epistasis could be impacted by the fitness of the initial variant for the GB1 landscape (26), but we were surprised to see that the mean fraction stayed fairly constant for the TrpB landscape (mean sign epistasis: Q1=35% vs. Q4=31%; mean reciprocal sign epistasis: Q1=9% vs. Q4=8%), with the variance of the distributions decreasing with increasing fitness (SI Appendix Fig S21, Table S12). The mean fraction was more variable across fitness quartiles in the GB1 landscape, supporting previous results, but changes were overall modest (mean sign epistasis: Q1=35% vs. Q4=28%; mean reciprocal sign epistasis: Q1=11% vs. Q4=5%).

When grouping across pairs of positions for the pairwise analysis, we expected to see differences between the distributions of epistasis types for each pair, especially since the TrpB landscape is composed of two pairs of positions on either side of a cofactor. We were surprised to see how similar the fractions of each type of epistasis were across position pairs (Fig 2C,D), with averages ranging from 30–37% sign and 8–12% reciprocal sign for TrpB and 27–35% sign and 4–25% reciprocal sign for GB1 (SI Appendix, Table S13). The 41/54 pair in GB1 was a notable standout as it had significantly more reciprocal sign epistasis than the other pairs (25% vs. 13% for the next highest pair). The distributions were quite broad, however, and there was a wide range of fractions of each epistasis type for a given starting sequence (SI Appendix, Fig S22–23). The fact that the average fractions of each type of epistasis are so similar across six pairs of positions that are close in proximity in both a binding protein and an enzyme is striking. Further investigation of other multi-site saturation landscapes is required to probe the generality of this observation.

Next, we quantified the magnitude of the epistasis types for both landscapes and compared all of these effects to a null model built from an additive landscape injected with noise based on that of the TrpB landscape (Methods, Construction of a null model). The null model had 74% magnitude, 22% sign, and 4% reciprocal sign epistasis—significantly less sign and reciprocal sign epistasis than the TrpB and GB1 landscapes (SI Appendix, Fig S21–24, Table S13–14). Furthermore, the extent of these effects is much smaller than in the TrpB and GB1 landscapes (average absolute value of epsilon for each epistasis type (magnitude/sign/reciprocal sign) for null model: 0.11/0.23/0.39; TrpB: 0.34/0.59/1.04; GB1: 0.41/0.78/1.56; SI Appendix, Fig S25–27). Finally, we examined how well a global epistasis model proposed by Otwinowski *et al.* (36), which seeks to capture how a nonlinear function may transform underlying additive interactions on a trait into epistatic ones, accounts for the epistasis found in the TrpB and GB1 landscapes (SI Appendix, Figure S28). We observed correlations of r^2^=0.59 for GB1 and r^2^=0.76 for TrpB between the true fitness and the fitness predicted by the global epistasis model, suggesting a mixture of both global and complex epistasis contributes to the epistasis in both landscapes.

### Epistasis constrains navigation of sequence-fitness landscapes

To investigate how navigation of the landscape is constrained by epistasis, we first built directional graphs linking any active variant to the best variant in the landscape, AIKG, via single substitutions (Fig 3A, Methods, Path analyses). For this analysis, only direct paths were considered, with a maximum number of steps equal to the Hamming distance (HD) between the initial and final variants (i.e., we did not allow “side-steps” through other variants via extradimensional bypass (26)). Using these graphs, we determined the fraction of starting points which have at least one possible path to the top and found that, if no deleterious steps are allowed, ∼20% of the starting points cannot reach the global optimum, AIKG, via any single-step path, and therefore must navigate sign and/or reciprocal sign epistasis to do so (Fig 3B). Accounting for assay noise or a less restrictive evolutionary pressure, we looked at how changing the allowed magnitude of deleterious steps enabled access to more paths to the best variant. We allowed steps accompanied by a 0% to a 100% decrease in fitness (at which point all steps are allowed, and thus all paths accessible) and observed that even allowing any step up to 50% worse still resulted in 6.8% of starting variants having no possible pathway to the top. Such strongly deleterious substitutions are unlikely to accumulate during natural or laboratory evolution under an explicit selective pressure. During directed evolution, screening throughput is usually limiting, and resources must be allocated with care. There are far more deleterious substitutions than beneficial ones and no way to know which downward step eventually leads to a more-fit variant. This makes accepting deleterious substitutions a highly risky decision, as one must effectively take a blind chance. In reality, such variants would be discarded and prevented from reaching the global optimum in an evolutionary search.

**Figure 3.**
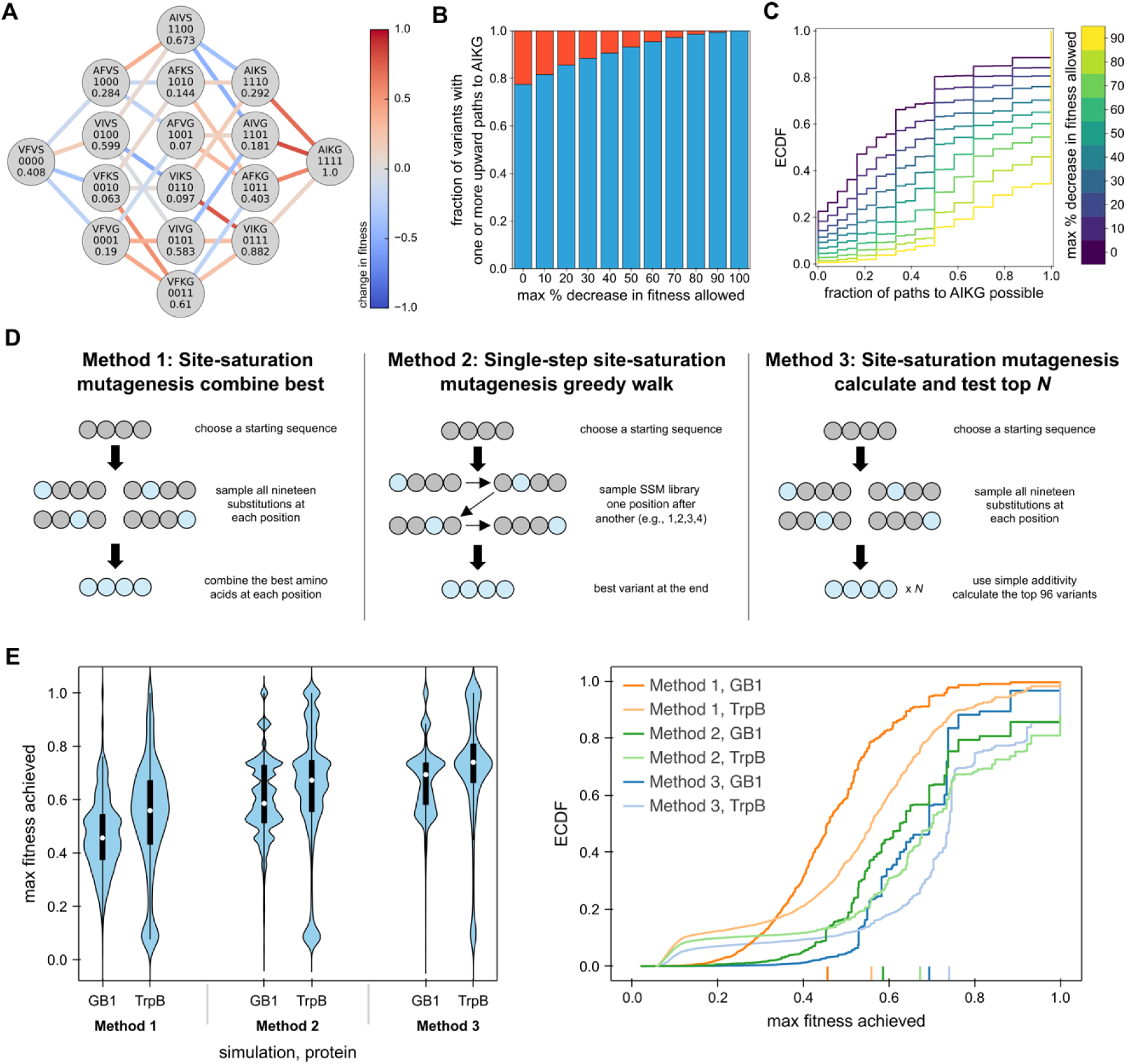
Investigation of evolutionary constraints in enzyme fitness landscapes. **A** Path map from the parent variant (*Tm*9D8*, VFVS) to the top variant in the landscape (AIKG). Nodes are labeled with the amino acids at positions 183, 184, 227, and 228 along with the fitness of that variant. Paths are colored by the change in fitness between the two variants. **B** A path map like the one pictured in A can be built connecting every detectably active variant to the top variant for a total number of path maps equal to the total number of active variants. Considering each of these maps, the fraction of maps with at least one possible path to the top variant is colored in blue while the fraction with no possible paths is colored in red. When no downward steps are allowed, max % decrease in fitness allowed = 0% and only strictly neutral or beneficial substitutions are allowed. This stringency is relaxed by accepting increasingly deleterious substitutions up to 100% (where all paths are accessible). **C** An empirical cumulative distribution function built from all possible starting points and displaying the fraction of paths reaching the top, given a specified cutoff. The x-axis denotes the fraction of possible paths to the top variant, and the y-axis denotes the fraction of starting variants which have up to that fraction of paths possible to the top variant. **D** Three different baselines of directed evolution methodologies. **E** The max fitness achieved from each starting point is plotted as a violin and empirical cumulative distribution function (ECDF) for each of the three directed evolution simulation methodologies. We show the results for the 4-site-saturation landscape on TrpB and on GB1. For the ECDF, color indicates the simulation method, with TrpB results in the lighter shade and GB1 results in the darker. A left-shifted curve indicates fewer starting variants can achieve a high max fitness.

These evolutionary constraints can be investigated in more detail using empirical cumulative distribution functions (ECDFs) that represent the fraction of variants that have at least a given fraction of paths accessible to the top variant (Fig 3C). The greater the number of variants that display a low fraction of accessible paths to the top (e.g., 1/24 paths), the more left-shifted the ECDF. When no deleterious steps are allowed, only ∼25% of the paths to the top variant are accessible from the median starting variant (the ECDF at *y* = 0.5). To enable the median starting variant to have all paths to the top variant accessible, one must be willing to accept steps with up to a 90% reduction in fitness.

We next examined the local optima, defined as variants where no single substitution of an active variant yields an improvement. For TrpB, there are 520 total optima (5.3% of the active variants), one of which is the global optimum, AIKG (SI Appendix, Fig S29). The second and third highest optima, CLKG and VLCS, have fitness values of 0.93 and 0.75, respectively. Of the 519 local optima, 510 (98.3%) could be escaped via two simultaneous substitutions, while the remaining nine required three simultaneous substitutions. The GB1 landscape has only 29 total optima (26). We also examined the local optima within the null model landscape, finding that experimental noise may make low-fitness regions appear more rugged. This landscape had 87 local optima, which were all below 10% of the max fitness (SI Appendix, Fig S29). Accounting for this, the TrpB landscape still had 175 local optima with fitnesses above that of the highest fitness local optimum in the null model landscape—over five times as many as GB1. These data could perhaps suggest that enzyme fitness landscapes are inherently more rugged (e.g., perhaps due to higher sensitivity to substitutions for ground states and transition states), but we caution against generalizing before many more such comparisons are made, because ruggedness is highly dependent on site- and selection-based factors as well.

For further characterization of the local optima, we focused on the top 20 (fitness values ranged from 0.38–1). Allowing no downward steps, we observed a tendency for the number of starting variants with at least one path to the local optimum to decrease as the fitness of the local optimum decreased (SI Appendix, Fig S29–30). This could be due to multiple factors, one of which is that more of the paths must pass through variants with higher fitnesses than the optima, which would make them inaccessible. A strong correlation between optima fitness and basin of attraction size (number of variants that can access an optimum) has also been reported (22). Most of the top 20 local optima remained reasonably accessible, however, meaning they could trap single-step experimental approaches. Altogether, these results suggest that the TrpB landscape is enriched in evolution-constraining epistasis compared to GB1, and experimental paths may be trapped more easily at local optima.

We also examined how the effects of epistasis influence the results of directed evolution. We considered three different directed evolution approaches that can also serve as competitive benchmarks for predictive approaches: Method 1) site-saturation mutagenesis (SSM) at each of the four sites in parallel followed by recombination of the best variants at each site; Method 2) single-step sequential SSM, using the best variant at one site as the parent for the next until all four sites have been examined, starting from any of the sites; and Method 3) SSM at each site in parallel followed by direct synthesis of the top *N* additivity-predicted variants — 96 variants in this case (Fig 3D). Because we enforced the sampling of every single substitution during the site-saturation mutagenesis steps, we would expect to obtain the max fitness every time with each of these approaches if the landscapes exhibited no epistasis. Only Method 2 has the potential to navigate sign and reciprocal sign epistasis, as it samples a new sequence context in each round of SSM, and therefore can discover previously deleterious substitutions that have become beneficial (37). However, the substitutions must be made in the correct order, which is unknown *a priori*, for it to be efficient.

For both the TrpB and the GB1 data, we saw the same pattern of performance: Method 3 performed the best, then Method 2, and then Method 1 (Fig 3E, F), starting from one of the top 9,783 variants in either landscape (the number of active variants in the TrpB landscape). The same simulations were run for the null model, where the same pattern of performance was observed, but with a much higher probability of reaching the global maximum for all methods (SI Appendix, Fig S31, Table S15). This suggests that the epistasis due to noise alone is insufficient to exert a strong influence over the outcome of directed evolution on this landscape. Increasing the number of variants tested in the second round of Method 3, *N*, improved the results slightly (SI Appendix, Fig S32). Note, however, that Method 3 requires direct synthesis of these *N* sequences, which can result in significant extra cost over Methods 1 and 2. Changing the number of starting points used for GB1 changed the results significantly, with simulations starting from any active variant performing much worse (SI Appendix, Fig S33). If simulations are allowed to start from any variant (active or inactive), the performance was also much worse for both landscapes (SI Appendix, Fig S34). The performance drop was similar for the two landscapes for Methods 1 and 3, but Method 2 on GB1 saw a much less drastic drop in performance, working better than Method 3 on average. This suggests that Method 2 may be more robust than Method 3 and able to increase fitness more reliably in noisy, low-fitness regimes. These results in combination with time and cost considerations provide guidelines for selecting an experimental approach to explore epistatic regions of sequence space.

### Experimental design enables capture of enzymatic parameters of interest

The conditions of the pooled-culture enrichment assays and the starting sequence for this experiment were chosen to improve the chances of capturing enzyme-specific attributes and avoid the dominance of stability effects. To verify that this was indeed the case, we compared observed fitness values with catalytic parameters of select variants and measured stability changes. As noted previously, sequences with K227 dominated the growth assay despite lysine at this site being nearly non-existent across known TrpB-like sequences. This suggested to us that K227 may exert some deleterious effect that is not observed under the assay conditions but is subject to natural selection. For example, it may increase the *K*_M_ for indole such that it is not competitive under physiological conditions but works well when indole is added exogenously at 200 µM, as in this assay. Alternatively (or additionally), K227 may be highly destabilizing, but not enough to unfold the thermostable *Tm*TrpB variant at *E. coli* growth temperatures. (Indeed, this possibility motivated the use of a thermostable parent sequence at the outset of this study.) Therefore, we chose a set of variants to characterize in more depth, including all variants in the single possible path from *Tm*9D8* to the top variant (Fig 3A), as well as the four other variants in the top five (CLKG, ALKG, CIKG, and VLKG), which also all contained K227.

To assess stability, we determined the temperature at which a 1-hour incubation causes an irreversible 50% reduction in activity as compared to a room temperature incubation (*T*_50_) (Table 1, SI Appendix, Table S16, Fig S35–36, Methods, *T*_50_ measurements). The only possible upward path between *Tm*9D8* (VFVS) and AIKG first requires F184I, which exerts no effect on *T*_50_, followed by S228G, which imparts a >1 °C increase. From here, two large decreases in stability come from the remaining two substitutions: a change of more than -7.2 °C from V227K and -1.7 °C from V183A to 91.0 °C — a *T*_50_ about 8.0 °C lower than the starting variant under these conditions. All five top variants (all of which contain K227) exhibited *T*_50_ values similar to AIKG (90.1–93.2 °C, or 5.8–8.9 °C below the starting variant). Drops in stability this large would likely lead to loss of function under native conditions, where proteins are typically only marginally more stable than the optimal growth temperature of their host (38), suggesting one reason why K227 is absent from the evolutionary record despite emerging as highly fit at *E. coli* growth temperatures.

**Table 1.** Enzymatic parameters measured for selected variants.

| **enzymatic parameter** | **variant** |  |  |
| --- | --- | --- | --- |
|  | *Tm*9D8* (VFVS) | VIVG | AIKG |
| $k_{\text{cat}}$ (min<sup>-1</sup>) | 22.6 ± 0.3 | 51.1 ± 1.0 | 67.1 ± 1.1 |
| $K_{\text{M,serine}}$ (mM) | 0.30 ± 0.04 | 0.18 ± 0.01 | 0.17 ± 0.02 |
| $K_{\text{M,indole}}$ (μM) | 22.9 ± 1.4 | 4.2 ± 0.6 | 20.2 ± 1.5 |
| $k_{\text{cat}}/K_{\text{M,indole}}$ (M<sup>-1</sup>s<sup>-1</sup>) | 1.6 ± 0.1×10<sup>4</sup> | 2.0 ± 0.3×10<sup>5</sup> | 5.5 ± 0.4×10<sup>4</sup> |
| $T_{50}$ (°C) | 99.0 ± 0.6 | >100 | 91.0 ± 0.3 |

Additionally, we measured the kinetic parameters of the starting variant, *Tm*9D8* (VFVS), the best variant (AIKG), and VIVG, the variant with two wild-type reversions and high stability in the middle of the path from *Tm*9D8* to AIKG. All three enzymes were expressed and purified for characterization via Michaelis-Menten kinetics (Table 1, Methods, Enzyme purification, crystallography, and measurement of kinetic parameters, SI Appendix, Fig S37–39). We observed that the *k*_cat_ values for the three enzymes roughly mirrored the fitness values we obtained for them in the high-throughput growth assay. We also observed that the two wild-type reversions caused a significant decrease in *K*_M_ for indole in VIVG, while the *K*_M_ of AIKG for indole was similar to *Tm*9D8*.

Importantly, while AIKG displays a higher rate of Trp formation at the 200 µM indole concentration used during the growth assay, it reacts more slowly than VIVG at indole concentrations below ∼50 µM (SI Appendix, Fig S38), which may better represent its native conditions. Both AIKG and VIVG had *K*_M_ values for Ser roughly half that of *Tm*9D8* (Table 1). These results, coupled with the decrease in stability, help explain how K227 can be nearly non-existent in native TrpB enzymes but optimal in this near-native assay. The observation of such effects was enabled by a highly thermostable parent enzyme that decouples mutation effects on stability from effects on activity.

### Classifying active and inactive variants using off-the-shelf fitness predictors

Given the complexity of enzyme catalysis, we suspected that structure-based predictors of the fitness effects of substitutions that work well for a small binding protein (15, 39) may not be as useful for an enzyme. Instead, we recognized that the abundance of TrpB sequences can be used to generate deep multiple sequence alignments (MSAs) that are more amenable to evolutionary-scale predictors (40, 41), especially since we assayed a near-native function of TrpB. We analyzed the enzyme data with both Triad protein design software using a Rosetta energy function (Protabit, Pasadena, CA, USA: https://triad.protabit.com), which provides a score that aims to predict stability, and EVmutation (40), which gives a score that aims to predict the fitness effect of a given set of substitutions based on conservation and evolutionary couplings. As a starting structure for the Triad calculations, we obtained a 2.15-Å resolution structure of *Tm*9D8* (PDB ID: 8VHH, SI Appendix, Table S17). We found that EVmutation is the better classifier of active and inactive variants for the TrpB landscape, despite performing poorly for GB1, which has a much less diverse MSA (Fig 4A). However, neither EVmutation nor Triad is well correlated to fitness for TrpB (Fig 4B) or GB1 (Fig 4C). Despite this, sampling variants above a threshold from EVmutation can increase the fraction of fit variants sampled as well as significantly improve the mean fitness of a sample. A Triad score threshold is less successful for TrpB, struggling to enrich in both fraction active and mean fitness despite working well for GB1.

**Figure 4.**
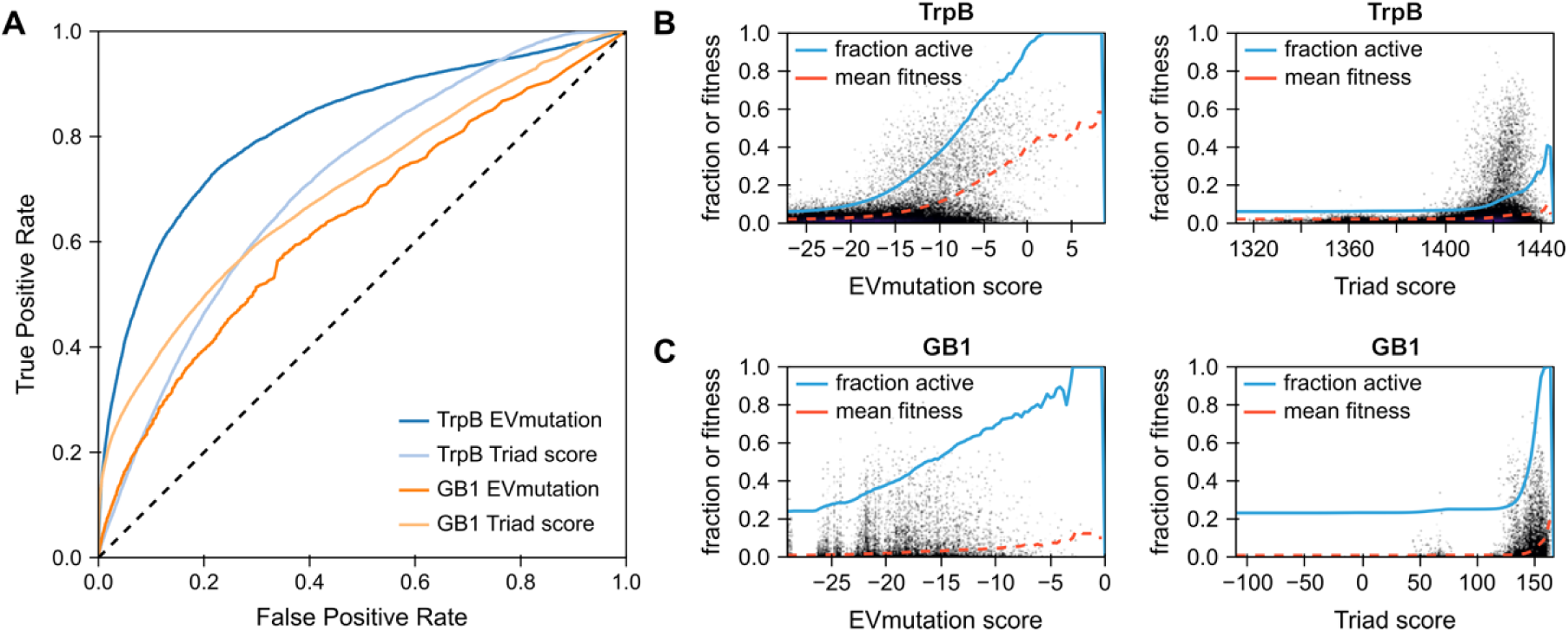
Assessment of simple fitness predictors on the TrpB landscape. **A** Receiver operating characteristic (ROC) curve for performance of EVmutation and Triad as classifiers of active and inactive variants for the TrpB and GB1 landscapes. The dashed black line represents the performance of a non-predictive model. Left- and up-shifted curves indicate better performance. **B** True TrpB fitness plotted versus EVmutation or Triad scores. **C** True GB1 fitness plotted versus EVmutation or Triad scores. For plots in both B and C, the blue line indicates the fraction of variants above the predictor threshold which are active while the dashed red line indicates the average fitness of the variants above the predictor threshold.

## Discussion

TrpB is conserved across all domains of life, acting in primary metabolism to perform the final step of Trp biosynthesis. Here we provide a combinatorially complete, 160,000-variant fitness landscape of substitutions at four residues in the active site of this ubiquitous enzyme, the first landscape of its kind that reports on enzyme catalysis. The topography of this landscape reflects significant epistasis, which results in many indirect adaptive paths and local optima that can stymie traditional directed evolution methodologies. We expect this landscape to provide a useful testing ground for laboratory and predictive protein engineering approaches as we learn to navigate epistatic enzyme fitness landscapes.

To generate a landscape useful for examining epistasis, we first generated nine 3-site libraries (8,000 variants each) across 20 different sites, aiming to identify sites that were permissive enough to retain a significant number of active variants to analyze. From this we identified four sites in TrpB enriched in epistatic mutations to construct the full 160,000-variant library. The resulting landscapes and the epistasis they contain reflect the fitness of the TrpB variants under the selected experimental conditions, which can influence the extent of the epistasis in a landscape (42). Calculation of pairwise epistasis of both the TrpB and previous GB1 landscapes showed that both have significant sign and reciprocal sign epistasis that occurs between all sampled positions and for initial variants of all fitness levels. This epistasis was shown to constrain the ability of directed evolution to reach the fitness peak, and our evolution simulations showed that some methods are more effective than others. However, the methods we tested require the measurement of different numbers of variants, a different number of evolution rounds, and different costs. For example, although site-saturation mutagenesis at all positions followed by testing of the top 96 additively predicted variants performed the best, direct synthesis of variants can be expensive. If synthesis cost is a major factor, other methods might be better choices, at the cost of time.

The assay conditions in this study were designed to capture catalytic parameters, but ultimately the high-throughput fitness measurements are scalar quantities resulting from the aggregate influence of factors such as stability, substrate binding, catalytic rate, and environment on growth of the *E. coli* expressing TrpB. The emergence of the destabilizing but activating K227 substitution suggests that catalytic rate is a prominent factor contributing to the calculated fitness values. However, here we characterized only a few of the most active variants; loss of stability could have caused catastrophic loss of fitness for others that were not observed (43). Emerging high-throughput stability (44) and kinetics measurements (45) could further disentangle the contributions of stability and activity within this fitness landscape.

Navigation of epistatic landscapes is made more efficient by recognizing that the sources of epistasis are diverse, and these non-linear effects can arise due to changes in any of the myriad factors contributing to fitness. Our results also show that the epistasis of both the GB1 and TrpB landscapes arises from both global and complex (local) epistasis, increasing the complexity that must be considered. Within complex functions such as catalysis, there can be many potential sources of epistasis. For example, beneficial substitutions that reduce stability may remain beneficial if they still meet minimal stability requirements, but when combined they could push the protein over the stability threshold and ablate activity altogether (46–48). Alternatively, epistasis can arise via a change in the rate-limiting catalytic step (49). Even if perfect predictors existed for each factor contributing to fitness, any single predictor may break down in regimes where another factor dominates. Alone, methods that predict specific attributes of protein fitness cannot be expected to accurately model a sequence-fitness landscape, especially of a complex enzymatic task. However, we can envision a multi-modal approach where models predicting specific facets of enzyme fitness are aggregated into a final fitness prediction for a specific set of conditions.

These methods might be composed of supervised and semi-supervised data-driven models, physics-based approaches, or a combination, leveraging the information of datasets across different attributes of protein fitness. Developing such approaches will require time, testing, and data, and this epistatic enzyme landscape provides a compelling and useful challenge for them through close examination of an instance where traditional laboratory evolution methods struggle.

## Materials and Methods

### Construction of Trp-auxotrophic *Escherichia coli* strain and *Tm*9D8* plasmids

The Trp auxotroph used for the 3- and 4-site pooled-culture enrichment assays was constructed from BW25113, a K-12 derivative and the parent strain for the Keio collection of single-gene knockouts (50), using λ red-mediated gene replacement (51). Both *trpA* and *trpB* were deleted and replaced with a kanamycin resistance cassette to use a selection for the strain. Initial studies were done in an alternate NEB^®^ 5-alpha strain (New England Biolabs, Catalog # C2987H) where *trpA* and *trpB* were replaced with a chloramphenicol resistance cassette. Further details in SI Appendix, *Escherichia coli* Trp knockout strain construction.

Two different *Tm*9D8* constructs were made for this study. For the pooled-culture enrichment assays, *Tm*9D8* was cloned into the arabinose-inducible pBAD24 backbone while for *in vitro* assays and protein expression for purification, the pET22b(+) backbone was used. pBAD24-sfGFPx1 was a gift from Sankar Adhya & Francisco Malagon (Addgene plasmid # 51558; http://n2t.net/addgene:51558; RRID:Addgene_51558) (52). Further details in SI Appendix, *Tm*9D8* plasmid construction.

### Preliminary *in vitro* rate of tryptophan formation assays

Protein was expressed for *in vitro* assays in T7 Express cells (New England Biolabs, Catalog # C2566H) in 96-well deep-well plates using the pET22b(+) vector. Colonies were picked into each well and grown to stationary phase overnight in Luria Broth containing 100µg/mL carbenicillin (LB_carb_). These cultures were used to inoculate expression plates of Terrific Broth containing 100 µg/mL carbenicillin (TB_carb_). Upon completion of expression, *E. coli* cells were pelleted by centrifugation, the supernatant was decanted, and pellets were frozen for later use. Further details in SI Appendix, Deep-well plate protein expression.

Single-site saturation preliminary data were obtained from Wittmann *et al.* (34) and double-site saturation preliminary data were obtained using the same procedure starting from deep-well expression plates described above. Sequences were obtained using evSeq (34).

### DNA library construction and cloning methods

In this study, libraries were constructed using NNK degenerate codons or the 22-codon-trick (53). Constant regions of target constructs were amplified using Phusion^®^ High-Fidelity DNA Polymerase according to manufacturer recommendations (New England Biolabs, Catalog # M0530L). For small libraries (1- or 2-site libraries) degenerate primers were used directly for amplification off the plasmid template. However, to reduce bias in the larger 3- and 4-site libraries, inner primers were first used to amplify regions adjacent to the variable positions via PCR. These fragments were then purified and used as template for a second PCR where degenerate primers introduced the variation of the library. Fragments were assembled into circular plasmid using NEBuilder^®^ HiFi DNA Assembly (New England Biolabs, Catalog # E2621X) following manufacturer instructions.

Smaller libraries (1- or 2-site) were transformed directly into Trp auxotroph cells, but higher efficiency was needed for the 3- and 4-site libraries. Therefore, assembled circular plasmid DNA was first transformed into NEB^®^ 10-beta electrocompetent *E. coli* (New England Biolabs Inc., Catalog # C3020K). These cells were incubated overnight, and plasmid was prepared via miniprep the following day. These plasmid preps were used as the source of library DNA for downstream pooled-culture enrichment assays. More details on library construction (including primers) are available in SI Appendix, DNA library construction.

### Preliminary plate-based independent growth assays

Libraries were first transformed into the Trp auxotroph strain (SI Appendix, Preparation of Trp auxotroph electrocompetent cells and electroporation). These libraries were plated onto LB agar supplemented with 35 µg/mL kanamycin and 100 µg/mL carbenicillin. Liquid LB_carb,kan_ was inoculated with single colonies, and cultures were grown overnight at 37 °C, 220 rpm, and 80% humidity for 16–20 h. Cultures were diluted 20- to 200-fold into Trp-dropout media (see SI Appendix, Trp-dropout media) into UV-transparent microplates (Caplugs/Evergreen Catalog # 290-8120-0AF) to a total volume of 200 µL. Plates were then incubated at 37 °C and 240 rpm with a 2 mm amplitude in a Tecan^®^ SPARK^TM^. Absorbance at 600 nm (OD_600_) was measured every 10 min for 12–48 h to monitor cell culture density. Between readings the plate remained covered.

### Pooled-culture enrichment assay

To perform the pooled-culture enrichment assay, electrocompetent Trp auxotroph cells were transformed with the relevant DNA library. After a 1-h rescue, cells were transferred to LB_kan,carb_ and incubated overnight at 37 °C and 220 rpm for 16–20 h. At this point, cells were either used directly or diluted 1:1 with sterile 50% glycerol, aliquoted, and frozen at -80°C for later use. Frozen aliquots were prepared by thawing on ice and transferring into LB_carb,kan_ and incubated overnight at 37 °C and 220 rpm for 16–20 h. From here, the LB culture was pelleted by centrifugation at 5,000 *g*, the supernatant was decanted, and the pellet was resuspended in Trp-dropout (Trp-DO) media (single- and double-site libraries) or 1X PBS, pH 7.4 (triple- and quadruple-site libraries) (Invitrogen, Catalog # AM9625). The resuspension was once again pelleted by centrifugation at 5,000 *g*, and the supernatant was decanted to remove as much Trp in the solution as possible. The pellets were then resuspended in Trp-DO media to the OD_600_ reported in Tables S7–S9.

Cells were incubated in Trp-DO media at 37 °C and 250 rpm in total volumes of 25 (1-, 2-, or 3-site libraries) or 50 mL (4-site library), with 1.5-mL samples collected at each timepoint seen in Tables S7–S9. These samples were centrifuged at 5,000 *g* for 5 minutes and stored at -20 °C until further use and sequencing preparation.

### Sequencing library preparation and data pre-processing

All libraries were prepared via a two-step PCR approach with attempts to minimize amplification cycles so as to reduce bias introduced by PCR. In the first step, primers that included complementarity to the region of interest, a 10-nucleotide random sequence (10xN), and handles for Illumina barcodes were used in a two-cycle PCR. These PCR products were digested with ExoCIP (New England Biolabs, Catalog # E1050L) and used as template for a second, 10-cycle PCR using IDT^®^ for Illumina^®^ DNA/RNA UD Indexes Set A, Tagmentation (Illumina, Catalog # 20027213). Samples were then digested with DpnI according to manufacturer directions (New England Biolabs, Catalog # R0176L), and the DNA products isolated via magnetic bead cleanup using Agencourt AMPure XP (Beckman Coulter, Catalog # A63880) according to manufacturer recommendations. Sample concentrations were measured using Quant-iT^TM^ PicoGreen^TM^ (ThermoFisher Scientific, Invitrogen, Catalog # P7581) and pooled equimolarly for submission to high-throughput sequencing with an Illumina HiSeq2500. Further details in SI Appendix, Sequencing library preparation.

Processing of the sequencing data was performed to provide filtered and aligned data for determining accurate, sequence-dependent codon/amino acid counts. Forward and reverse reads were filtered independently using `fastq-filter` (https://github.com/LUMC/fastq-filter) with an average read quality (option `–q`) of `25` (or an error rate of 0.00316 across the 51 sequenced bases). Corresponding forward and reverse reads were then matched, retaining only pairs of reads that passed both filters. The first 13 bases of each read were trimmed to remove sequence- and experiment-specific identifiers. Pairs of files were then aligned using `minimap2` (https://github.com/lh3/minimap2) using the following process: ‘`minimap2 –ax sr ref.fasta forward.fastq reverse.fastq -k 5 -w 3`’, where `ref.fasta` is a fasta file containing the *Tm*9D8* (parent) reference sequence, `forward.fastq` is the filtered and trimmed forward fastq file, and `reverse.fastq` is the matching filtered and trimmed reverse fastq file. The option `-w 3` was used due to the trimmed reads being short (min 38 bp) to provide a sufficiently small window for proper alignment. (The `-k 5` option is based on the standard ratio of `k/w` ∼1.5.) Aligned reads were then filtered based on the following criteria: both reads must align with no indels, starting at the first aligning base of the trimmed forward read in the *Tm*9D8* sequence with a total length for each aligned forward and reverse read equal to the expected value. For example, the 4-site library started at base 517 and spanned 193 bases to the beginning of the reverse read. Codon identities for each position were then indexed from these filtered and aligned reads for determining fitness. Python scripts and documentation can be found in the associated code.

### Fitness score calculations

Fitness calculations were determined based on theory proposed by Kowalsky *et al.* to obtain specific growth rates for each variant (35). For each timepoint captured, a specific growth rate, μ_*i*_, was calculated for each unique amino acid sequence as follows:

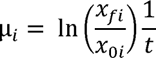

where *x*_0*i*_ and *x_fi_* represent the concentration of *E. coli* harboring the given amino acid sequence *i*, in the initial population and the population at time *t* respectively. These values were calculated based on the OD_600_ of the culture and the frequency of sequence in each population. Sequences with fewer than an average of ten sequencing counts at *t* = 0 or zero counts in either replicate at any other timepoint were omitted from the fitness calculation pipeline. We observed that sequences containing stop codons, which can be presumed non-functional, had slightly non-zero μ_*i*_ values. Therefore, we subtracted the average μ_*stop*_ from each μ_*i*_. Finally, we divided this value by the background-subtracted μ_*max*_ for that timepoint to normalize the maximum fitness of each landscape to 1.

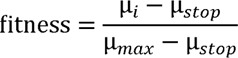

Fitness values were then calculated for each timepoint in a landscape and averaged for each replicate where applicable (all landscapes besides 3-site landscapes A, B, and C). The activity threshold was imposed at this point by enforcing that the fitness for a variant in each replicate was at least 1.96 standard deviations above the mean fitness (97.5^th^ percentile) of all stop-codon-containing sequences. Therefore, for the 4-site landscape, variants were labeled active if 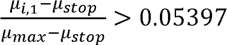 and 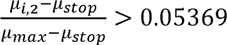 and inactive if not, resulting in just 1.05% of the stop-codon-containing sequences being labeled active. In libraries where only one replicate was sequences (3-site landscapes A, B, and C), the 99.5% percentile was used for the single replicate (2.576 standard deviations above the mean). Finally, the fitness values for the two replicates were averaged together to obtain a final fitness metric for each sequence. Timepoints were omitted from several of the 3-site libraries when there was minimal fitness separation of stop-codon-containing sequences and remaining variants (as compared to later timepoints) or there was poor correlation between variant fitnesses in both replicates likely due to sample preparation error (see Table S8 for details). To enable the analyses in this paper, fitness values for the missing variants in the 4-site landscape were imputed with the sklearn KNN imputer (Fig S16), and the imputed scores reported previously by the authors were used for GB1 (26).

### Pairwise epistasis calculations and analyses

Fraction of pairwise epistasis was calculated using python and functions that classify each type into one of the three categories: magnitude, sign, and reciprocal sign epistasis. Notably, for this analysis additive effects were grouped into magnitude. For each unique starting variant (**00**), all possible double substitutions (**11**) were tested such that all variants within the set of **00**, **01**, **10**, and **11** were active and an epistasis type was assigned. Doing this for all possible double substitutions, we computed the fraction of each type of epistasis for the starting variant. In order to reduce the impact of noisy, low fitness measurements, we only classified epistasis for sets of variants (00, 01, 10, 11) where all variants had fitness greater than or equal to that of the minimum fitness of active variants. These results were sorted into quartiles based on the fitness values of the starting variant to create the distributions.

Epsilon was calculated as described by Khan *et al.* (54) for all variant sets using the initial variant as the reference:

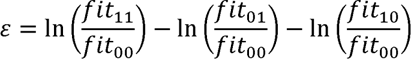

Since variant sets were defined with a directionality between a pair of variants (00 to 11), each pair of variants was examined twice, once considering each variant as the initial. This resulted in perfectly symmetric distributions of epsilon. To reduce the redundancy of the resulting distributions, we examine the absolute value of epsilon in this text.

### Construction of a null model

To construct the null model, we began with the parent, VFVS, and sampled all single substitutions from there. We then calculated the fold-change in activity relative to parent for each of these substitutions and used them to calculate a perfectly additive landscape. To add noise to the landscape, we fit an exponential distribution to the distribution of differences in fitness between the two replicates of the TrpB landscape and sampled from it, randomly adding or subtracting the value from the fitness. The resulting landscape had the same parent fitness as the TrpB landscape and a maximum fitness of 1.23. The activity threshold was defined as the minimum fitness of the of a variant labeled “active” in the TrpB landscape, resulting in 5,404 active variants.

### Path analyses

For each active variant, networkx (55) was used to construct a directed graph from that variant to the best variant in the landscape, AIKG. For this analysis, any fitness below zero was set to zero. No imputed variants were used as starting points, but they were used as intermediate variants when necessary, so that no graphs had missing nodes. The number of direct paths possible to the top variant were counted allowing downward steps ranging from 0% to 90% decrease in fitness. The same analyses were run for each of the top twenty local optima.

### Determination of local optima

Local optima were determined by looking at all non-imputed variants classified as active. For each of these variants, all single substitutions were made *in silico*. If no single-substitution variant had a higher fitness than the original, that original variant was classified as a local optimum. To determine if two simultaneous substitutions could enable escape from the local optimum, all double substitutions were made *in silico*. If at least one double-substitution variant had a higher fitness than the original, that original variant was said to be able to escape the local optimum via double-site saturation mutagenesis. Details can be found in the associated code.

### Simulations of directed evolution

#### Method 1: Site-saturation mutagenesis combine best

For every variant classified as active, all nineteen substitutions are made *in silico* at each of the four positions independently in the background of the initial sequence. The best amino acid at each position is obtained and the sequence consisting of these amino acids is built. The best variant from among the initial sequence, all single-site mutagenesis variants, and the recombined variant is reported as the maximum fitness achieved.

#### Method 2: Single-step site-saturation mutagenesis greedy walk

For every variant classified as active and each possible order of sampling positions (*M*! where *M* = number of positions) site-saturation mutagenesis is performed iteratively *in silico* with *M* rounds. For the first position, all nineteen substitutions are tested for that position in the starting background of the other positions. Once the best residue for that position is determined, it is fixed, and the next position is targeted, and so on until all positions have been targeted once. The fitness of the final variant is reported as the max fitness achieved.

#### Method 3: Site-saturation mutagenesis calculate and test top *N*

For every variant classified as active, all nineteen substitutions are made *in silico* at each of the four positions independently in the background of the initial sequence. Using these *M* x 19 + 1 (starting variant) datapoints, fitness scores for all 20*^M^* possible combinations are calculated as the product of the fold-change for each single substitution over the initial sequence. These sequences are then ranked, and the best *N* are tested *in silico*. The max fitness achieved is reported as the maximum fitness of the initial sequence, any of the single substitutions, and the top *N* predicted sequences.

### Generating off-the-shelf fitness predictions

Using the methods from Wittmann *et al.* (15) we obtained fitness predictions using both Triad estimates of ΔΔG and EVmutation (40). For Triad (https://triad.protabit.com), the crystal structure obtained here was used as the starting point for the calculations and EVmutation (https://evcouplings.org/) used *Tm*9D8* as the starting sequence with a bitscore of 0.3.

### Enzyme purification, crystallography, and measurement of kinetic parameters

T7 Express cells were transformed with pET22b(+) harboring the enzyme sequence of interest and plated onto LB agar supplemented with 100 µg/mL carbenicillin. Single colonies were transferred into 5-mL liquid LB_carb_ and grown overnight at 250 rpm and 37 °C. Expression cultures were started by inoculating 250 mL of TB_carb_ with 1 mL of each overnight culture and subsequently grown at 300 rpm and 37 °C. After 6 h of outgrowth, 250 µL of 1 M IPTG were added to each culture and the cultures were grown an additional 22 h at 300 rpm and 30 °C. (*Note:* Expression levels of *Tm*TrpB were found to be high when cultures were dense at the time of induction; OD_600_ was not measured.) Cultures were pelleted via centrifugation at 5,000 *g* for 20 min and then frozen at -20 °C.

For purification, cell pellets were thawed at room temperature before resuspension with 10 mL of lysis buffer containing 25 mM potassium phosphate, 100 mM NaCl, and 20 mM imidazole, pH 8.0 (Buffer A) supplemented with 200 µM PLP, 0.02 mg/mL DNAse I, 1 mg/mL lysozyme, and 0.1X BugBuster^®^. This cell suspension was incubated at 37 °C and 220 rpm for 1 h and, as all enzymes displayed *T*_50_ measurements much greater than 75 °C, was followed by a 30 min heat treatment at 75 °C to further disrupt remaining *E. coli* proteins. The lysate was clarified by centrifugation at 14,000 *g* for 20 min and the supernatant was collected. Gravity columns containing 2.5 mL Ni-NTA (Qiagen, Catalog # 30210) were pre-equilibrated with >10 column volumes (CV) of Buffer A. Clarified lysate was then added to the columns, and they were washed with another 10 CV of Buffer A. Protein was eluted with 50% Buffer A and 50% of an elution buffer made with 25 mM potassium phosphate, 100 mM NaCl, and 500 mM imidazole, pH 8.0 (Buffer B). An additional 1 mM PLP was added to each sample, and they were buffer exchanged into KPi by dialysis.

Protein concentrations were obtained with the Pierce BCA Protein Assay Kit (ThermoFisher Scientific, Catalog # 23225), and purified protein was frozen in aliquots on dry ice. Protein was used directly from these aliquots for crystallography with no buffer exchange. Full crystallography and structure determination protocols are provided in SI Appendix, *Tm*9D8* crystallization, *Tm*9D8* crystal structure determination.

Enzyme parameters, including *K*_M_ and *k*_cat_, were determined via Michaelis-Menten kinetics by collecting 290 nm absorbance continuously over 500 seconds with a UV spectrophotometer (Shimadzu, Catalog # EW-83400-20) for reactions containing enzyme (62.5–250 nM), indole (1.56–500 µM), Ser (0.05–20 mM), and DMSO (4%) in KPi. Indole *K*_M_ was collected at 20 mM Ser and Ser *K*_M_ was collected at 200 µM indole (the concentration used for the growth rate assay). Initial rates were obtained using linear or exponential fits of the data within a time frame not impacted by burst phase kinetics. These rates were fit with a Michaelis-Menten model to obtain estimates for *K*_M_ and *k*_cat_.

### *T*_50_ measurements

Using the plasmids prepared via site-directed mutagenesis, variants were expressed as described in “Deep-well plate protein expression” with six biological replicate wells for each variant. Frozen pellets were fully thawed at room temperature and then resuspended by light vortexing in lysis buffer composed of 1 mg/mL lysozyme, 0.1X Bug Buster^®^, 0.2 mg/mL DNase I, and 200 µM PLP in 50 mM KPi. Plates were incubated at 37 °C, 220 rpm, and 80% humidity for 1 h and then clarified by centrifugation at 4,500 *g* for 10 min. Clarified lysate for each variant was pooled into individual 15-mL conical centrifuge tubes and stored at 4 °C until needed.

For heat treatments, 40 µL of clarified lysate was aliquoted into full-skirted PCR plates and incubated at the reported temperature for 1 h using a gradient on a Mastercycler^®^ X50s (Eppendorf, Catalog # 6311000010). Room temperature controls were incubated for 1 h on the benchtop in 200-µL PCR tubes. Room temperature controls were then added to the PCR plate, and all samples were centrifuged for 8 min at 4,000 *g* to remove accumulated debris. Rate of Trp formation was measured as described in Wittmann *et al.* (34) using 20 µL lysate, 20 µL KPi, and 160 µL of reaction master mix.

## Acknowledgments

The authors thank Igor Antoshechkin (Caltech Millard and Muriel Jacobs Genetics and Genomics Laboratory) for discussions on library preparation and performing library sequencing. The authors also thank Bruce Wittmann for discussions on building landscapes and Sabine Brinkmann-Chen for reviewing the manuscript. We would also like to thank Stephanie Breunig and Elliot MacKrell for discussion and materials for construction of the *E. coli* Trp auxotroph strain. Finally, we thank our reviewers, whose suggestions greatly strengthened the manuscript. This work was supported by the U.S. Department of Energy, Office of Science, Office of Basic Energy Sciences under award number DE-SC0022218 (to F.H.A.). This report was prepared as an account of work sponsored by an agency of the United States Government. Neither the United States Government nor any agency thereof, nor any of their employees, makes any warranty, express or implied, or assumes any legal liability or responsibility for the accuracy, completeness, or usefulness of any information, apparatus, product, or process disclosed, or represents that its use would not infringe privately owned rights. Reference herein to any specific commercial product, process, or service by trade name, trademark, manufacturer, or otherwise does not necessarily constitute or imply its endorsement, recommendation, or favoring by the United States Government or any agency thereof. The views and opinions of authors expressed herein do not necessarily state or reflect those of the United States Government or any agency thereof. Additional support was provided by the Amgen Chem-Bio-Engineering Award (to F.H.A.) and the Caltech Center for Environmental Sciences (to F.H.A.). Individual support comes from: Caltech AI4Science/Amazon AWS Fellowship (K.E.J.); Merck-Helen Hay Whitney Postdoctoral Fellowship (N.J.P.); National Science Foundation Graduate Research Fellowship (J.Y.).

