## Supporting Information for "A combinatorially complete epistatic fitness landscape in an enzyme active site"

[1] Division of Biology and Bioengineering, California Institute of Technology, Pasadena, CA 91125
[2] Division of Chemistry and Chemical Engineering, California Institute of Technology, Pasadena, CA 91125
[A] Present address: Merck & Co., Inc., South San Francisco, CA 94080
[B] Present address: Department of Biochemistry, Stanford University, Stanford, CA 94305
[C] Present address: Harvard Medical School, Boston, MA 02115
[D] Present address: Codexis Inc., Redwood City, CA 94063
[†]These authors contributed equally to this work and are listed alphabetically.

*Corresponding Author: Frances H. Arnold

**This PDF file includes:**

Supporting text
Figures S1 to S35
Tables S1 to S13
SI References

### I. Table of Contents

#### II. General procedures

***Escherichia coli* Trp knockout strain construction.** An initial Trp auxotroph strain used in preliminary assays was constructed from the NEB® 5-alpha strain (New England Biolabs, Catalog # C2987H) starting strain via λ red-mediated gene replacement (1). Both *trpA* and *trpB* were deleted and replaced with a chloramphenicol resistance cassette using primers based on those reported for deletion of *trpA* and *trpB* in the Keio collection. The chloramphenicol resistance cassette was amplified out of pKD3 with NEB5α_TrpAB::CamR_fwd and NEB5α_TrpAB::CamR_rev and used as the linear DNA for homologous recombination. Gene deletion was confirmed via colony PCR and Sanger sequencing with NEB5α_TrpAB_external_fwd and NEB5α_TrpAB_external_rev. All primers used can be found in Table S1. This strain was used for preliminary single- and double-site saturation for plate-based growth assays and pooled, sequencing-based growth assays.

Due to unusually slow growth we observed with larger libraries in the NEB® 5-alpha Trp auxotroph, we decided to build a new Trp-auxotroph strain with kanamycin resistance to see if this could be remedied. Starting with a K-12 derivative and the parent strain for the Keio collection of single gene knockouts (2), BW25113, we performed λ red-mediated gene replacement (1). Both *trpA* and *trpB* were deleted and replaced with a kanamycin resistance cassette using primers based on those reported for deletion of *trpA* and *trpB* in the Keio collection. The kanamycin resistance cassette was amplified out of pKD13 with BW25113_TrpAB::KanR_fwd and BW25113_TrpAB::KanR_rev and used as the linear DNA for homologous recombination. Gene deletion was confirmed via colony PCR and Sanger sequencing with BW25113_TrpAB_external_fwd, BW25113_TrpAB_external_rev, BW25113_TrpAB_internal_fwd, and BW25113_internal_rev. All primers used can be found in Table S1. This strain was used as the host organism for all reported triple- and quadruple-site saturation landscapes from pooled growth assays.

The protocol "Recombineering/Lambda red-mediated gene replacement" from OpenWetWare was used to construct both knockouts following the system from Datsenko & Wanner (1).

***Tm*9D8* plasmid construction.** *In vitro* assays and protein expression for purification were all performed with a pET22b(+) plasmid harboring *Tm*TrpB genes as previously reported for *Tm*9D8* (3). For use in the growth assays, we constructed an arabinose-inducible TrpB expression vector with the pBAD24 backbone. pBAD24-sfGFPx1 was a gift from Sankar Adhya & Francisco Malagon (Addgene plasmid # 51558; http://n2t.net/addgene:51558; RRID: Addgene 51558). The gene for *Tm*9D8* was exchanged for sfGFPx1 by amplification of *Tm*9D8* with TrpB_pBAD24_insert_fwd and TrpB_pBAD24_insert_rev and backbone amplification of pBAD24 with TrpB_pBAD24_bb_fwd and TrpB_pBAD24_bb_rev (primers found in Table S2). These pieces were then assembled into a circular plasmid via a two-piece Gibson assembly (4). The *Tm*9D8* sequence can be found in Table S3.

**Deep-well plate protein expression.** First, to prepare overnight culture plates, *E. coli* colonies from T7 Express (New England Biolabs, Catalog # C2566H) harboring TrpB variants in pET22b(+) vectors are picked into separate wells of a 96-well deep-well plate containing 300–500 µL of Luria Broth containing 100 µg/mL carbenicillin (hereafter referred to as $LB_{carb}$), covered with a microporous film, and grown overnight at 37 °C, 220 rpm, and 80% humidity for 16–20 h. For expression plates, new deep-well plates are prepared with 630 µL of Terrific Broth containing 100 µg/mL carbenicillin (hereafter referred to as $TB_{carb}$) and 20 µL of the overnight cultures is dispensed to each well. Expression plates are then incubated at 37 °C, 220 rpm, and 80% humidity for 6 h before addition of 50 µL of 14 mM IPTG in $TB_{carb}$ (final concentration 1 mM). These expression plates were then incubated at 30 °C, 250 rpm, and ambient humidity for 22 h, spun down for 5–10 min at 4500 g (until pellets form and supernatant is clarified). Supernatant was decanted, and expression plates were frozen at -20 °C for later use.

**Preparation of Trp auxotroph electrocompetent *E. coli* cells and electroporation.** On day 1, the strain of interest was streaked onto an LB agar plate containing the requisite antibiotic and grown overnight at 37 °C. On day 2, a single colony was picked into LB containing the requisite antibiotic and grown overnight at 37 °C and 220 rpm. On day 3, the overnight culture was diluted between 50- and 500-fold into Super Optimal Broth (SOB) medium (5) and grown at 18 °C and 220 rpm until OD ~0.4–0.6. Cultures were then plunged into ice-cold water for a minimum of 10 minutes until they reached 4 °C and then spun down at 5000 g for 5 min at 4 °C. The supernatant was decanted, and cells were resuspended in cold, sterile water to 1/5–1/10 the original volume. Cultures were spun down and resuspended a second time with the same procedure. One final spin was performed, and cells were resuspended in 1/100 the original volume (100X concentrated). Cells were used fresh on the day they were prepared.

Electroporation was performed by combining 50 µL of cells with 1–2 µL of plasmid in a chilled electroporation cuvette (1 mm or 2 mm gap; USA Scientific, 9104-1050 or 9104-5050) and applying current (BioRad MicroPulser$^{TM}$, Catalog # 165-2100, Ec1 or Ec3 respectively). Cells were then rescued via resuspension in Super Optimal broth with Catabolite repression (SOB medium with 20 mM glucose: SOC) and incubated for 15–60 min before plating onto LB agar plates or transferring to overnight cultures.

**Trp-dropout media.** Assay media was composed of 1X M9 salts (Sigma Aldrich, Catalog # M6030), 0.74 g/L dropout supplement -Trp (Takara Bio Inc., Catalog # 630413), 2 mM $MgSO_4$, 100 µM $CaCl_2$, and 0.4% glycerol. To prepare this media, M9 salts, dropout supplement -Trp, and glycerol were sterile filtered and stored at 4 °C. Stock solutions of $MgSO_4$ and $CaCl_2$ (1 M each) were sterilized by autoclaving and added to the media directly before beginning an assay. Antibiotics to select for the Trp auxotroph strain (35 µg/mL kanamycin) and the TrpB-containing plasmid (100 µg/mL carbenicillin) were also added to the media. Using pBAD24-*Tm*9D8*, arabinose concentrations from 0.001% to 0.1% and indole concentrations from 10 µM to 1000 µM were tested before choosing final concentrations of 0.05% arabinose (stock concentration 20% in M9) and 200 µM indole (stock concentration 500 mM in DMSO). This final mix is referred to as Trp-dropout (Trp-DO) media.

**DNA library construction.** Libraries were constructed via simultaneous site-saturation mutagenesis using either NNK degenerate primers or the "22 codon trick" (6). Unless otherwise stated, the following default PCR mix was used for all reactions, which used the Phusion® High-Fidelity DNA Polymerase according to manufacturer recommendations (New England Biolabs, Catalog # M0530L).

| Reagent | Volume (µL) |
|---|---|
| 5x HF Buffer | 10 |
| DMSO (100%) | 1.5 |
| dNTPs (10 mM) | 1 |
| Template | variable (x) |
| Phusion | 0.5 |
| Forward primer (10 µM) | 2.5 |
| Reverse primer (10 µM) | 2.5 |
| PCR water | 32 - x |
| Total | 50 |

The single- and double-site saturation DNA libraries were built with the 22-codon trick (6) using primers in Table S4. Extension time was varied based on fragment length.

| Temperature (°C) | Time (s) | Cycles |
|---|---|---|
| 98 | 00:30 | 1 |
| 98 | 00:10 | 1 |
| 55<br>Ramp speed: 1 °C / s | 00:15 | |
| 72 | variable | |
| 98 | 00:10 | 29 |
| 63 | 00:15 | |
| 72 | variable | |
| 72 | 05:00 | 1 |

| | | |
|---|---|---|
| 10 | infinite | 1 |

Each fragment was DpnI (New England Biolabs, Catalog # R0176L) digested and run on a 1% agarose gel containing SYBR™ Gold Nucleic Acid Gel Stain (ThermoFisher Scientific, Catalog # S11494). The relevant bands were excised, and the DNA fragments were purified using a Zymoclean Gel DNA Recovery Kit (Zymo Research, Catalog # D4002). Products were assembled into circular plasmid using NEBuilder® HiFi DNA Assembly (New England Biolabs, Catalog # E2621X) following manufacturer instructions. The resultant product was cleaned and concentrated with a DNA Clean & Concentrator®-5 kit (Zymo Research, Catalog # D4004).

When scaling up to larger DNA libraries composed of more possible sequences, we wanted to construct a relatively uniform input library and reduce the bias for the parent sequence; therefore, we adopted a two-step PCR approach used for both the triple- and quadruple site saturation libraries.

For the triple-site libraries, to produce the first fragment, an inner PCR (termed "gap" PCR) was performed where none of the variable region was included, using the gap primer (F gap) for its respective library (Table S5) along with AmpR_internal_rev (Table S4). The template plasmid used for all libraries was pBAD24-*Tm*9D8* except libraries F and G, which used a 301X plasmid library as the template sequence (prepared following the same method as described for single- and double-site saturation libraries using SSM primers Tm9D8*_301X_fwd and Tm9D8*_301_rev (Table S4). The following thermal cycler protocol was used for all reactions:

| **Temperature (°C)** | **Time (s)** | **Cycles** |
|---|---|---|
| 98 | 00:30 | 1 |
| 98 | 00:10 | 5 |
| 55 → 59 (+1 °C / cycle)<br>*ramp speed*: 1 °C / s | 00:15 | |
| 72 | 00:45 | |
| 98 | 00:10 | 25 |
| 59 | 00:15 | |
| 72 | 00:45 | |
| 72 | 10:00 | 1 |
| 10 | infinite | 1 |

The resulting PCR product was DpnI digested according to manufacturer directions and run on a 1% agarose gel containing SYBR™ Gold Nucleic Acid Gel Stain (ThermoFisher Scientific, Catalog # S11494). The relevant bands were excised, and the DNA fragments were purified using a Zymoclean Gel DNA Recovery Kit (Zymo Research, Catalog # D4002). This fragment was then used as template in a second PCR using the same reaction mix and thermal cycler settings but exchanging the "F gap" primer for the "F library" primer. This product was also run on a 1% agarose gel containing SYBR™ Gold Nucleic Acid Gel Stain, excised, and purified with a Zymoclean Gel DNA Recovery Kit.

The second fragment was constructed in a single step using the same reaction mix as used for the first fragment. For these reactions, the forward primer was the AmpR_internal_fwd (Table S4) and the reverse primer was a library specific "R primer" based on Table S5. The thermal cycler conditions used are stated below:

| Temperature (°C) | Time (s) | Cycles |
|---|---|---|
| 98 | 00:30 | 1 |
| 98 | 00:10 | 30 |

| | | |
|---|---|---|
| 72 | 00:15 | |
| 72 | 02:15 | |
| 72 | 10:00 | 1 |
| 10 | infinite | 1 |

This product was also run on a 1% agarose gel containing SYBR™ Gold Nucleic Acid Gel Stain, excised, and purified with a Zymoclean Gel DNA Recovery Kit.

Due to the design of the quadruple-site saturation library having two sets of two variable sites, the method for building it was slightly different than that used for the triple-site saturation libraries. The same two-step approached was used, where the inner region between the variable regions was first amplified without the regions to be diversified. This PCR used the default PCR mix, "F Gap" and "R Gap" primers from Table S6, the pBAD24-*Tm*9D8* as template, and the following PCR protocol:

| Temperature (°C) | Time (s) | Cycles |
|---|---|---|
| 98 | 00:30 | 1 |
| 98 | 00:10 | 5 |
| 72 → 68 (-1 °C / cycle)<br>*ramp speed*: 1 °C / s | 00:15 | |
| 72 | 00:30 | |
| 98 | 00:10 | 25 |
| 68 | 00:15 | |
| 72 | 00:30 | |
| 72 | 10:00 | 1 |
| 10 | infinite | 1 |

The resulting PCR product was DpnI digested and run on a 1% agarose gel containing SYBR™ Gold Nucleic Acid Gel Stain. The relevant bands were excised, and the DNA fragments were purified using a Zymoclean Gel DNA Recovery Kit. This fragment was then used as template in a second PCR using the same reaction mix and thermal cycler settings but using "F Library" and "R Library" from Table S6. The following thermal cycler program was used for amplification:

| Temperature (°C) | Time (s) | Cycles |
|---|---|---|
| 98 | 00:30 | 1 |
| 98 | 00:10 | 5 |
| 67 → 63 (-1 °C / cycle)<br>*ramp speed*: 1 °C / s | 00:15 | |
| 72 | 00:30 | |
| 98 | 00:10 | 25 |
| 63 | 00:15 | |
| 72 | 00:30 | |
| 72 | 10:00 | 1 |
| 10 | infinite | 1 |

This product was also run on a 1% agarose gel containing SYBR™ Gold Nucleic Acid Gel Stain, excised, and purified with a Zymoclean Gel DNA Recovery Kit.

The backbone was prepared in two pieces using the AmpR cassette break strategy described previously. The first backbone piece was prepared with the default PCR reaction mix using pBAD24-*Tm*9D8* as template and primers AmpR_internal rev (Table S4) and "F" from Table S6. The following thermal cycler program was used for amplification:

| **Temperature (°C)** | **Time (s)** | **Cycles** |
|---|---|---|
| 98 | 00:30 | 1 |
| 98 | 00:10 | 5 |
| 72 → 68 (-1 °C / cycle)<br>*ramp speed*: 1 °C / s | 00:15 | |
| 72 | 00:45 | |
| 98 | 00:10 | 25 |
| 69 | 00:15 | |
| 72 | 00:45 | |
| 72 | 10:00 | 1 |
| 10 | infinite | 1 |

The second backbone piece was also prepared with the default PCR reaction mix using pBAD24-*Tm*9D8* as template. The primers used for amplification were AmpR_internal_fwd (Table S4) and "R" from Table S6. The following thermal cycler program was used for amplification:

| **Temperature (°C)** | **Time (s)** | **Cycles** |
|---|---|---|
| 98 | 00:30 | 1 |
| 98 | 00:10 | 5 |
| 72 → 68 (-1 °C / cycle)<br>*ramp speed*: 1 °C / s | 00:15 | |
| 72 | 02:00 | |
| 98 | 00:10 | 25 |
| 69 | 00:15 | |
| 72 | 02:00 | |
| 72 | 10:00 | 1 |
| 10 | infinite | 1 |

Both backbone products were run on a 1% agarose gel containing SYBR™ Gold Nucleic Acid Gel Stain, excised, and purified with a Zymoclean Gel DNA Recovery Kit.

Once necessary fragments were prepared, DNA concentrations were measured with a GE Healthcare NanoVue™ Plus Spectrophotometer and they were assembled into circular plasmid using NEBuilder® HiFi DNA Assembly following manufacturer instructions. The resultant product was cleaned and concentrated with a DNA Clean & Concentrator®-5 kit. To achieve high transformation efficiency for the triple- and quadruple-site libraries, assembled plasmid libraries were first transformed into high-efficiency electrocompetent cells. We used NEB® 10-beta electrocompetent *E. coli* (New England Biolabs Inc., Catalog # C3020K) and the manufacturer recommended protocol. DNA concentration and electroporation settings were optimized to achieve high transformation efficiency. Following application of current, cells were rescued for only 15 minutes in the provided rescue media before being transferred to an overnight culture of $LB_{carb}$ and grown overnight at 37 °C and 220 rpm. Simultaneously, a dilution was plated on $LB_{carb}$ agar to be used to estimate the transformation efficiency and ensure sampling depth. The liquid cultures were miniprepped using the QIAprep Spin Miniprep Kit (Qiagen, Catalog # 27104) to prepare plasmid DNA libraries in high concentration and purity for transformation into Trp-auxotroph strains.

**Sequencing library preparation.** All libraries were prepared for sequencing with the same overall strategy using inner primers from Table S10. Mapping of these primers to the libraries can be found in Table S11. First, an initial two-cycle PCR amplification attached inner handles for the Illumina barcodes using the default PCR mix and the following thermocycler program:

| Step | Temperature (°C) | Ramp rate | Time | Cycles |
|---|---|---|---|---|
| Initial denaturation | 98 | max | 3 min | 1 |
| Denaturation | 98 | max | 30 sec | 2 |
| Anneal start temp | 64 | max | 1 sec | |
| Anneal slow ramp | 58 | 0.2 C/s | 90 sec | |
| Extension | 72 | max | 90 sec | |
| Final extension | 72 | max | 5 min | 1 |
| Hold | 4 | - | - | - |

These samples were then digested with ExoCIP (New England Biolabs, Catalog # E1050L) using a 20 min incubation at 37 °C followed by a 15 min inactivation step at 80 °C. The resulting product was used as template for a second PCR using IDT® for Illumina® DNA/RNA UD Indexes Set A, Tagmentation (Illumina, Catalog # 20027213).

| Reagent | Volume (µL) |
|---|---|
| PCR water | 1.25 |
| 5X KAPA HiFi | 5 |
| 10 mM dNTP | 0.75 |
| UDP Primer Mix | 5 |
| DNA eluate | 12.5 |
| KAPA HiFi Polymerase | 0.5 |
| Total | 25 |

| Step | Temp | Time | Cycles |
|---|---|---|---|
| Initial Denaturation | 98 | 00:30 | 1 |
| Denaturation | 98 | 00:10 | 10 |
| Anneal/Extend | 65 | 1:15 | |
| Final extension | 65 | 5:00 | 1 |
| Hold | 4 | Forever | |

Samples were then DpnI digested and purified via magnetic bead cleanup using Agencourt AMPure XP (Beckman Coulter, Catalog # A63880) according to manufacturer recommendations. Sample concentrations were measured using Quant-iT™ PicoGreen™ (ThermoFisher Scientific, Invitrogen, Catalog # P7581) and pooled equimolarly for submission to high-throughput sequencing with an Illumina HiSeq2500.

**Site-directed mutagenesis to construct variants for in-depth biochemical characterization.** Using a template plasmid of *Tm*9D8* in pET22b(+), primers were ordered to make exact mutations at positions 183, 184, 227, and 228. Full gene sequences are provided in Table S3.

***Tm*9D8* crystallization.** For the crystallization of tryptophan synthase variant *Tm*9D8*, the protein was purified as described previously (Methods, Enzyme purification, crystallography, and measurement of kinetic parameters). Due to the similarity between this enzyme and previously characterized tyrosine synthase variants, the same precipitant (1.2 M $NaH_2PO_4$/0.8 M $K_2HPO_4$, 0.1 M *N*-cyclohexyl-3-aminopropanesulfonic acid (CAPS), 0.2 M $Li_2SO_4$) was used to crystallize this variant. In a 24-well CrysChem M Plate (Hampton Research), 2 mg/mL protein were screened using 1–6 µL protein drops and 2–5 µL precipitant drops. While small salt crystals were observed in drops containing higher initial precipitant concentrations, drops with a higher protein concentration remained clear after six days. At this point, these drops (5–6 µL 2 mg/mL *Tm*9D8* + 2 µL precipitant) were streak seeded using a cat whisker, the generous gift of Crick Boville, and a seed stock derived from crystals of the related tyrosine synthase variant *Tm*TyrS1 (nine mutations). Within two days, small hexagonal prism crystals formed in these wells.

To prepare samples for x-ray diffraction experiments, a cryoprotectant solution was prepared by mixing 80 µL of equilibrated reservoir solution with 20 µL of ethylene glycol. This solution was then added to the crystal drop, sequentially adding and removing equivalent volumes until no schlieren was observed. Following cryoprotection, all crystals were mounted in nylon loops, cooled in liquid nitrogen, and stored prior to data collection.

***Tm*9D8* crystal structure determination**. Diffraction data were collected at the Stanford Synchrotron Radiation Laboratory (SSRL) beamline 12-2. Data reduction and integration were carried out using XDS (7) and scaled using Aimless in the CCP4 suite of programs (8). Molecular replacement (MR) was performed using the structure of holo *Tm*TyrS1 (PDB ID: 8EGY) as a search model in Phaser (9). Model building and modification in the electron density was performed using Coot and structure refinement was performed using Phenix (10, 11). Other ligands, including free PLP, as well as water molecules and ethylene glycol were added during later stages of refinement. Occasionally, spurious electron density peaks were present in the active site, dimer interface, and COMM domain that could not be unambiguously modeled by alternative protein conformations, solvent, or other additives applied during the procedure, so these were left uninterpreted. The quality of the final models was evaluated with MolProbity and PROCHECK (12, 13). Data collection and refinement statistics are presented in Table S17.

#### III. Figures

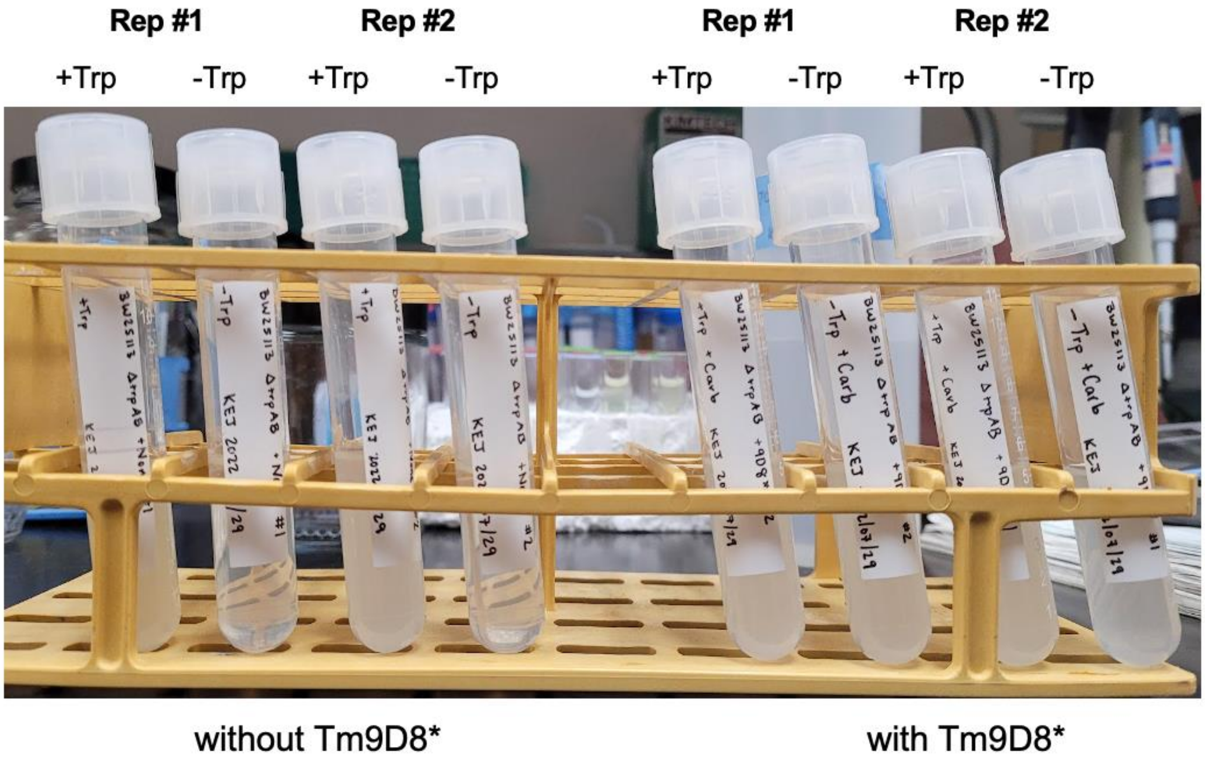

**Figure S1. TrpB- and Trp-dependent growth for the *E. coli* Trp auxotroph.** The left set of tubes are cultures where the Trp auxotroph is given no TrpB variant while the right set does harbor an efficient TrpB (*Tm*9D8*). Within the sets, cultures were grown with and without exogenous Trp added to the Trp-DO media (all cultures contained indole and arabinose). Only cultures where exogenous Trp is added, or the cells are harboring an active TrpB variant able to convert indole to Trp, show detectable growth.

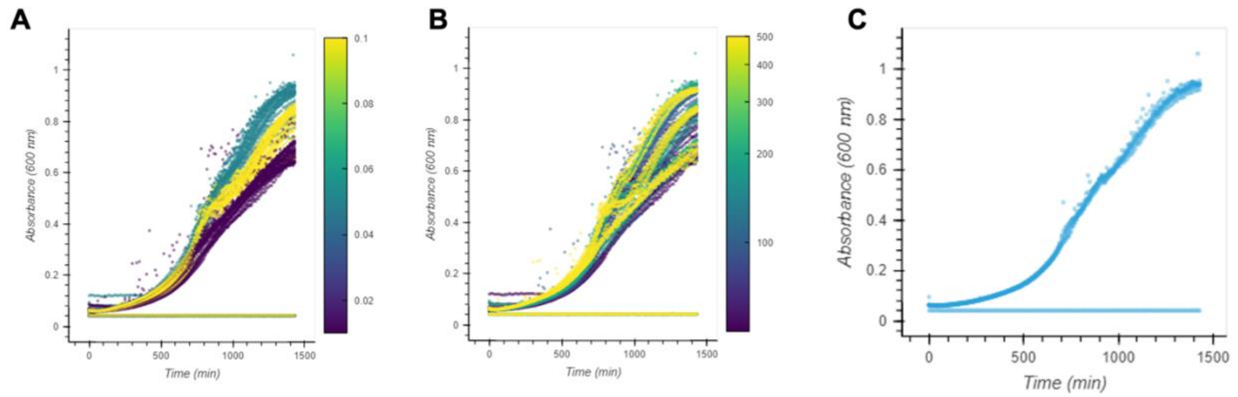

**Figure S2. Choosing growth assay conditions.** *E. coli* Trp auxotroph cells harboring pBAD24-*Tm*9D8* and grown in Trp-dropout media supplemented with arabinose and indole. Sterile wells appear as flat lines in this assay. **A** Absorbance at 600 nm ($OD_{600}$) versus time for all wells colored by arabinose concentration (%). This showed that 0.05% arabinose was optimal. **B** $OD_{600}$ versus time for all wells colored by indole concentration. This showed that 200 µM indole was optimal. **C** Replicates of $OD_{600}$ versus time at 0.05% arabinose and 200 µM indole.

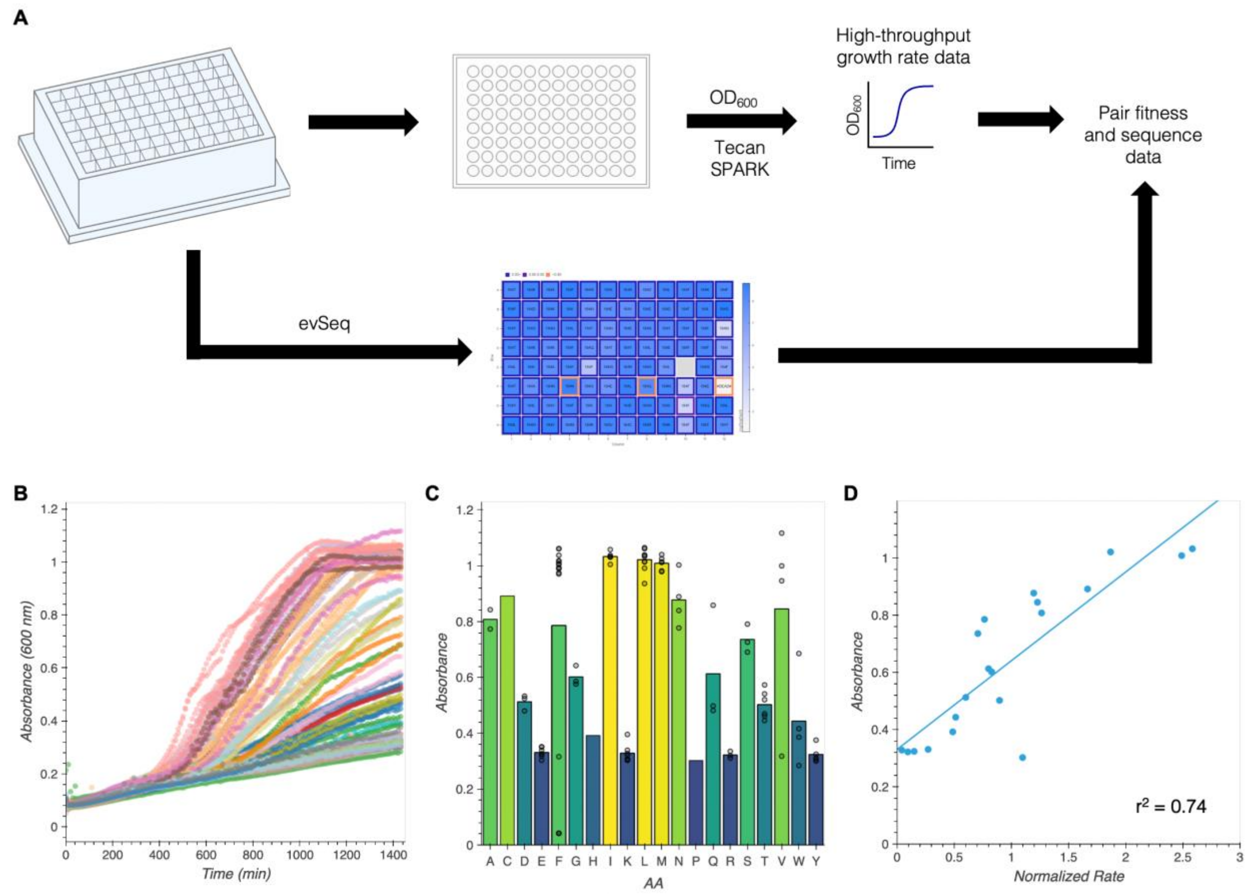

**Figure S3. Preliminary results for independent growth rates. A** Independent growth rates for a 184X library were monitored by measuring $OD_{600}$ over time and pairing it with sequencing data obtained by evSeq. **B** $OD_{600}$ vs time (min) with the scatter plots colored by amino acid identity. We observed clear clustering between amino acids that indicated reproducibility. **C** Average absorbance ($OD_{600}$) of the final timepoint along the collected data for each unique amino acid. Absorbances for each observed replicate are overlaid. **D** Average absorbance ($OD_{600}$) for each unique amino acid against the normalized rate of Trp formation (data obtained from Wittmann et al. (14)). There was a reasonable correlation between the two that indicated the fitness we were measuring was similar.

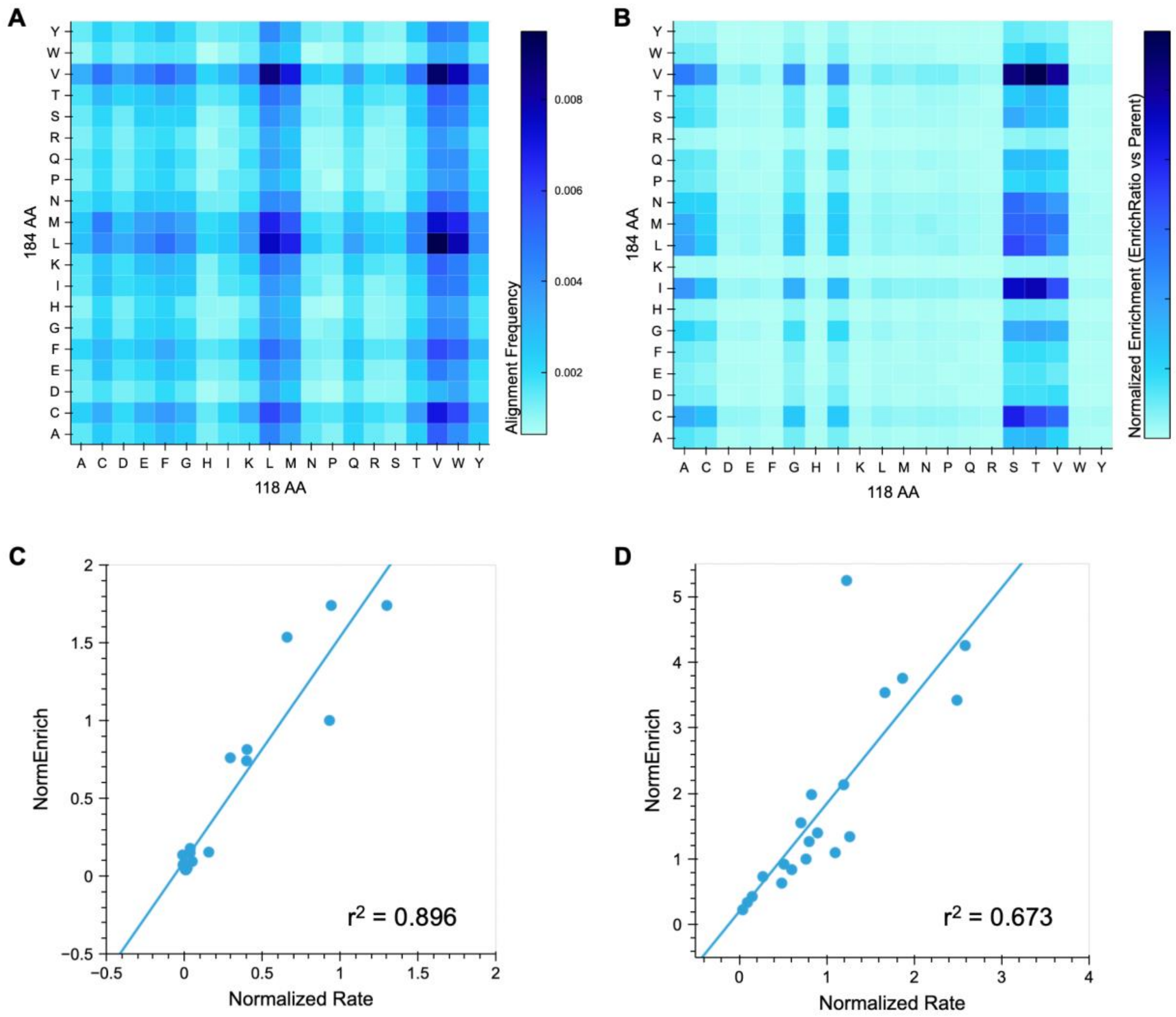

**Figure S4. 118X/184X growth-based enrichment assay. A** Frequency of each 118/184 amino acid pair in the input DNA library. **B** Normalized enrichment was calculated as the output sequencing frequency (at 22 h) divided by the input sequencing frequency and normalized to parent (A118, F184). **C** Correlation of normalized enrichment and normalized rate of Trp formation for all variants in the 118X/184X library with F184 (a proxy for a 118X library). **D** Correlation of normalized enrichment and normalized rate of Trp formation for all variants in the 118X/184X library with A118 (a proxy for a 184X library).

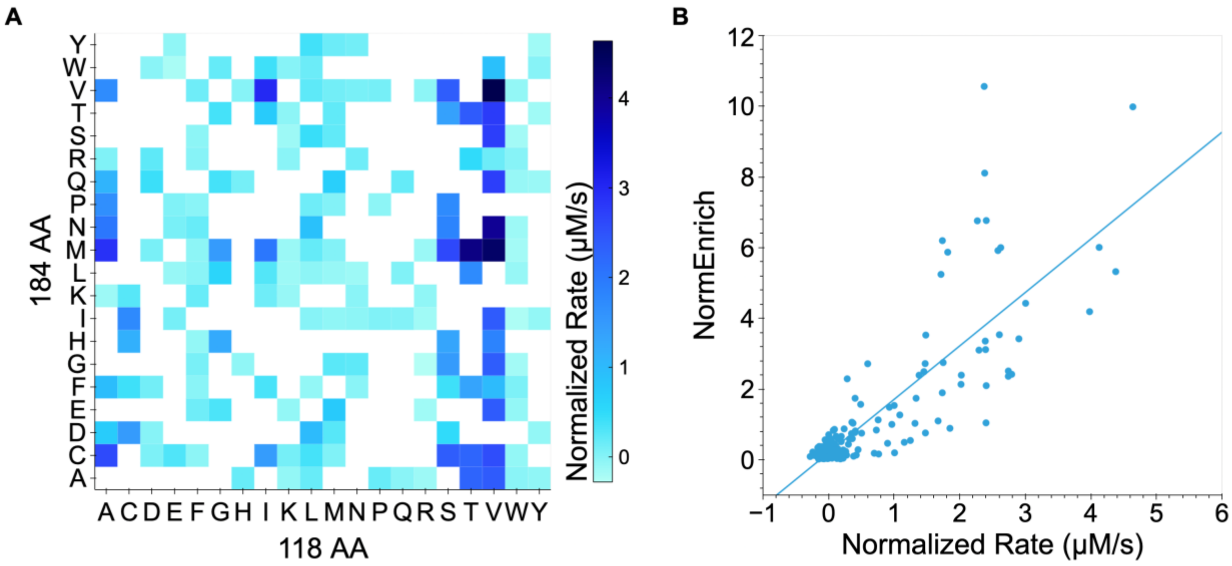

**Figure S5. Comparison of 118X/184X pooled-culture enrichment and *in vitro* rate of Trp formation. A** Normalized rate of Trp formation for a subset of the 118X/184X library. Missing values appear white. **B** Correlation between the normalized rate of Trp formation and normalized enrichment (output frequency divided by input frequency and normalized to parent A118/F184) for the subset of the 118X/184X library at 22 h.

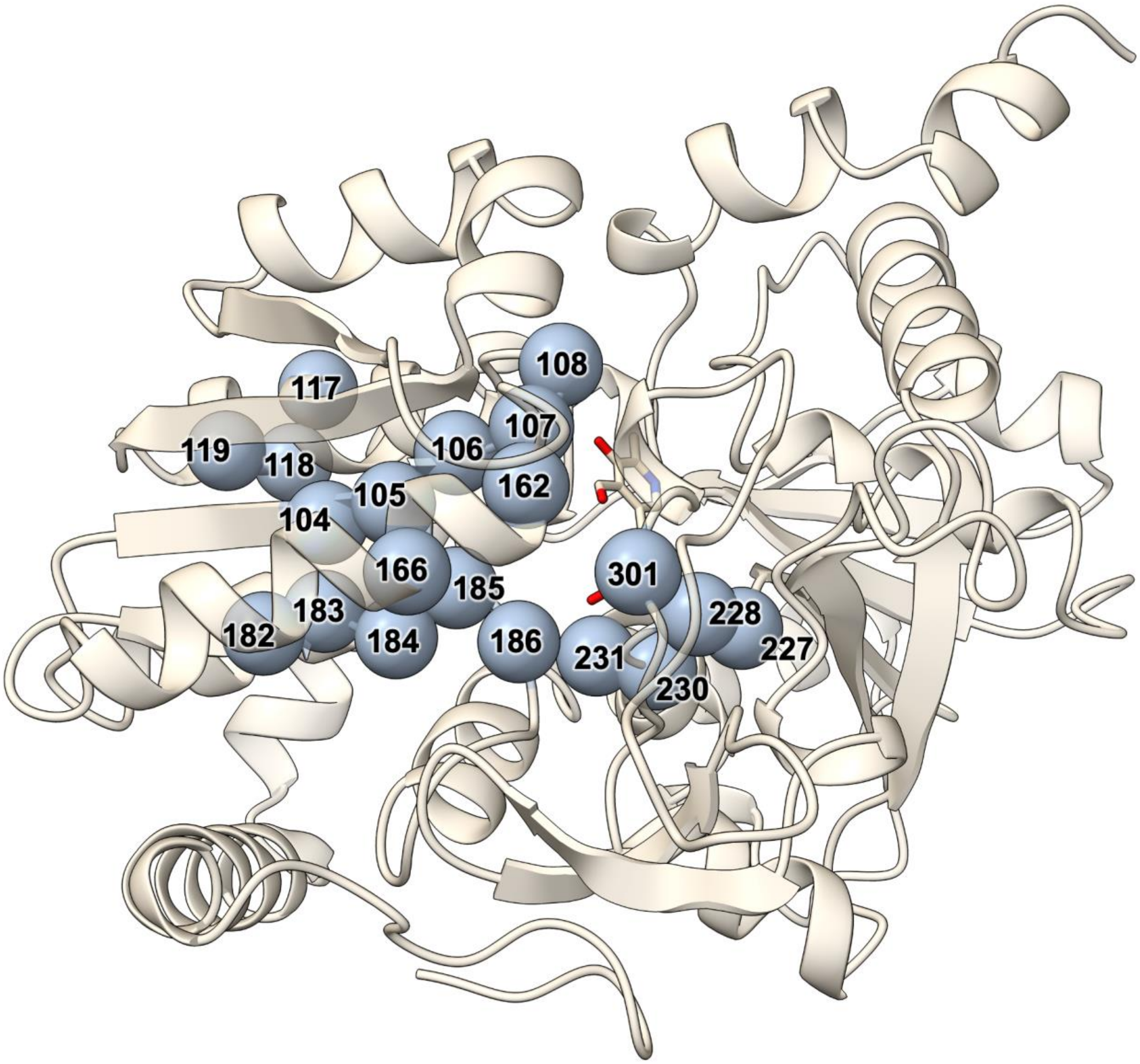

**Figure S6. All positions targeted within triple-site saturation libraries.** A total of twenty different residues were targeted among nine triple-site saturation libraries. Residues were chosen to be near the active site or known to modulate the activity of TrpB.

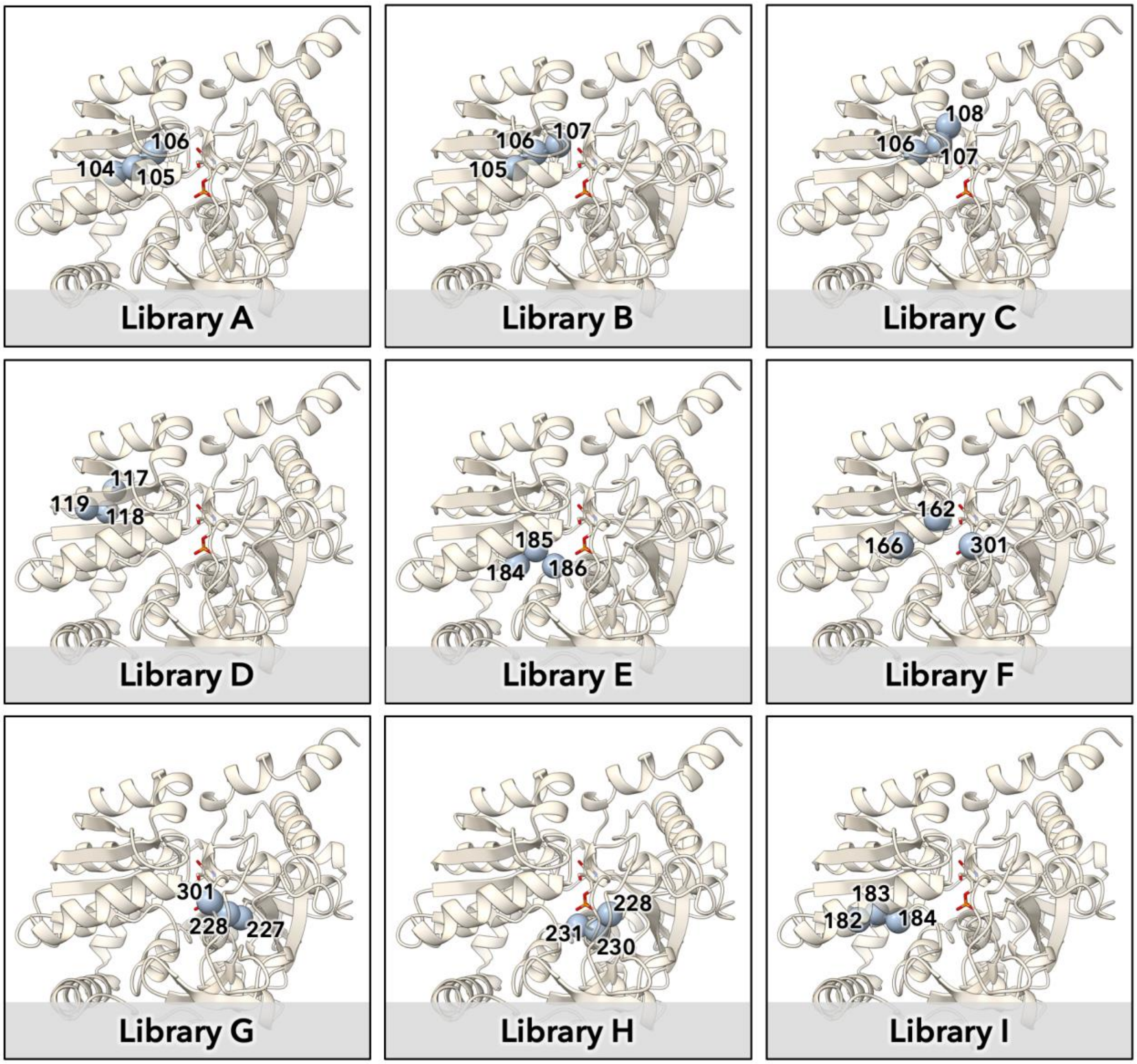

**Figure S7. Sets of residues chosen for the triple-site saturation libraries.** Nine sets of three residues were targeted based on proximity to each other as well as each of construction with available molecular biology methods. Different numbers of replicates and timepoints were obtained for libraries A, B, and C vs libraries D, E, F, G, H, and I.

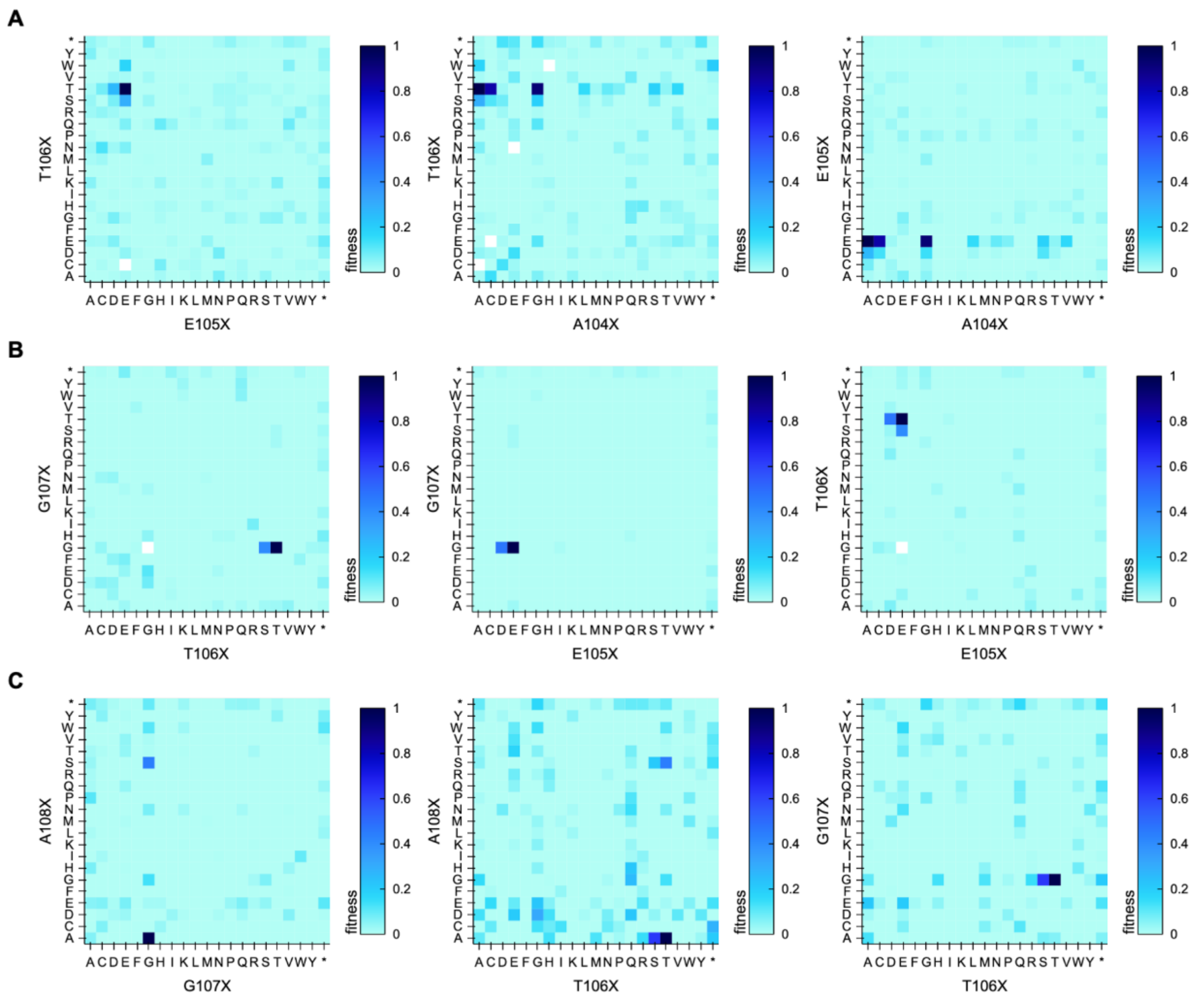

**Figure S8. Fitness for all pairs of residues for Libraries A, B, and C.** Initial investigations into the utility of these sets of positions for larger libraries. For each plot, the unplotted residue was held at the parent amino acid at that position, and for all three libraries very few amino acids are accepted at each position. Missing values appear white. **A** Library A separated into the three possible pairs of positions: 105 & 106, 104 & 106, and 104 & 105. **B** Library B separated into the three possible pairs of positions: 106 & 107, 105 & 107, and 105 & 106. **C** Library C separated into the three possible pairs of positions: 107 & 108, 106 & 108, and 106 & 107.

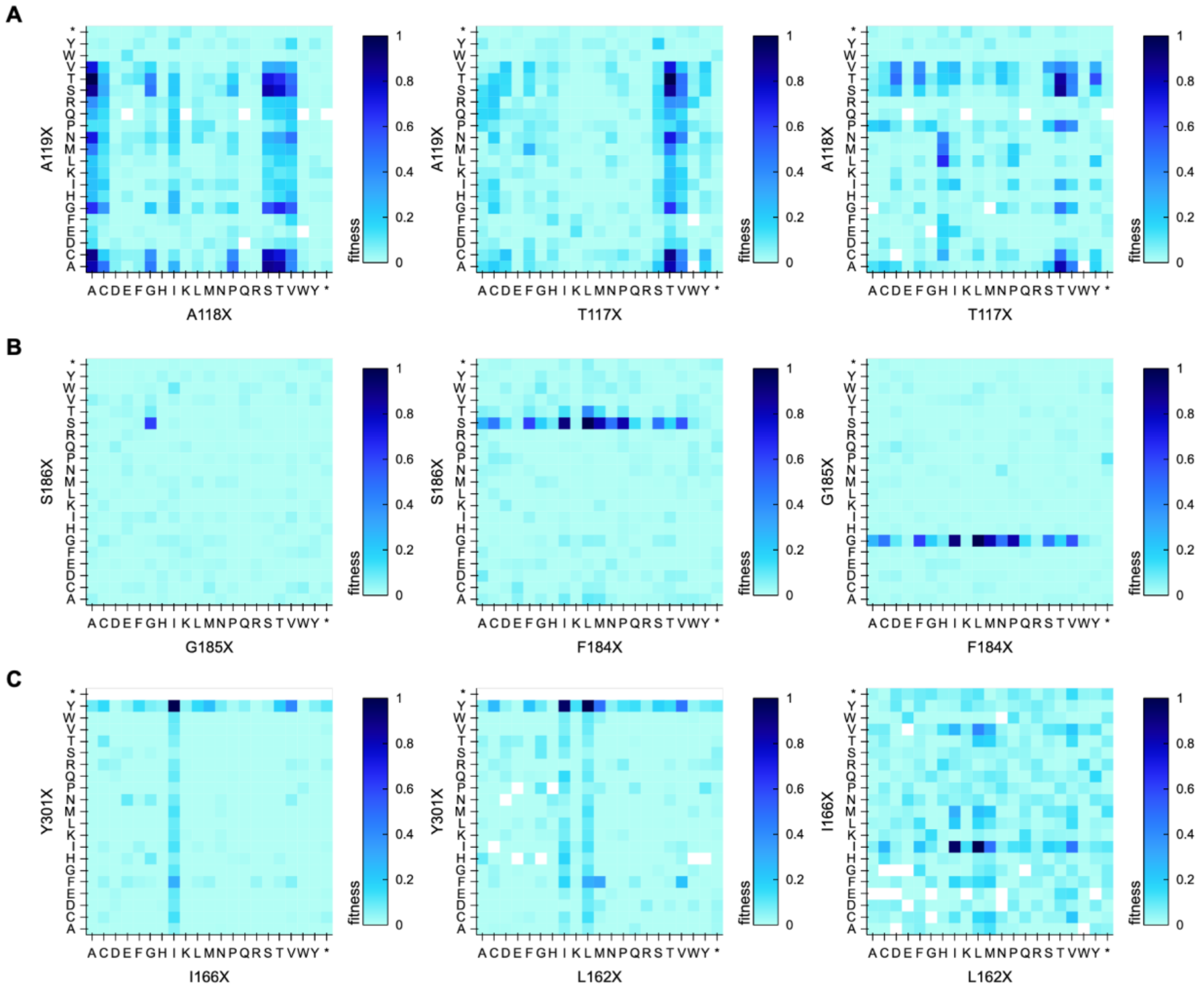

**Figure S9. Fitness for all pairs of residues for Libraries D, E, and F.** Initial investigations into the utility of these sets of positions for larger libraries. For each plot, the unplotted residue was held at the parent amino acid at that position. Missing values appear white. **A** Library D separated into the three possible pairs of positions: 118 & 119, 117 & 119, and 117 & 118. There is a relatively broad range of positions allowed at 118 and 119 while 117 allows many fewer. Interestingly, substitutions at 118 to leucine, methionine, or aspartic acid allow histidine to emerge as an option for 117. **B** Library E separated into the three possible pairs of positions: 185 & 186, 184 & 186, and 184 & 185. Position 184 shows a broad range of allowed amino acids while 185 and 186 are much more limited. **C** Library F separated into the three possible pairs of positions: 166 & 301, 162 & 301, and 162 & 166. This library allowed very few deviations from the parent sequence. Y301 appeared nearly mandatory as did I166 while either Leu or Iso were accepted at 162.

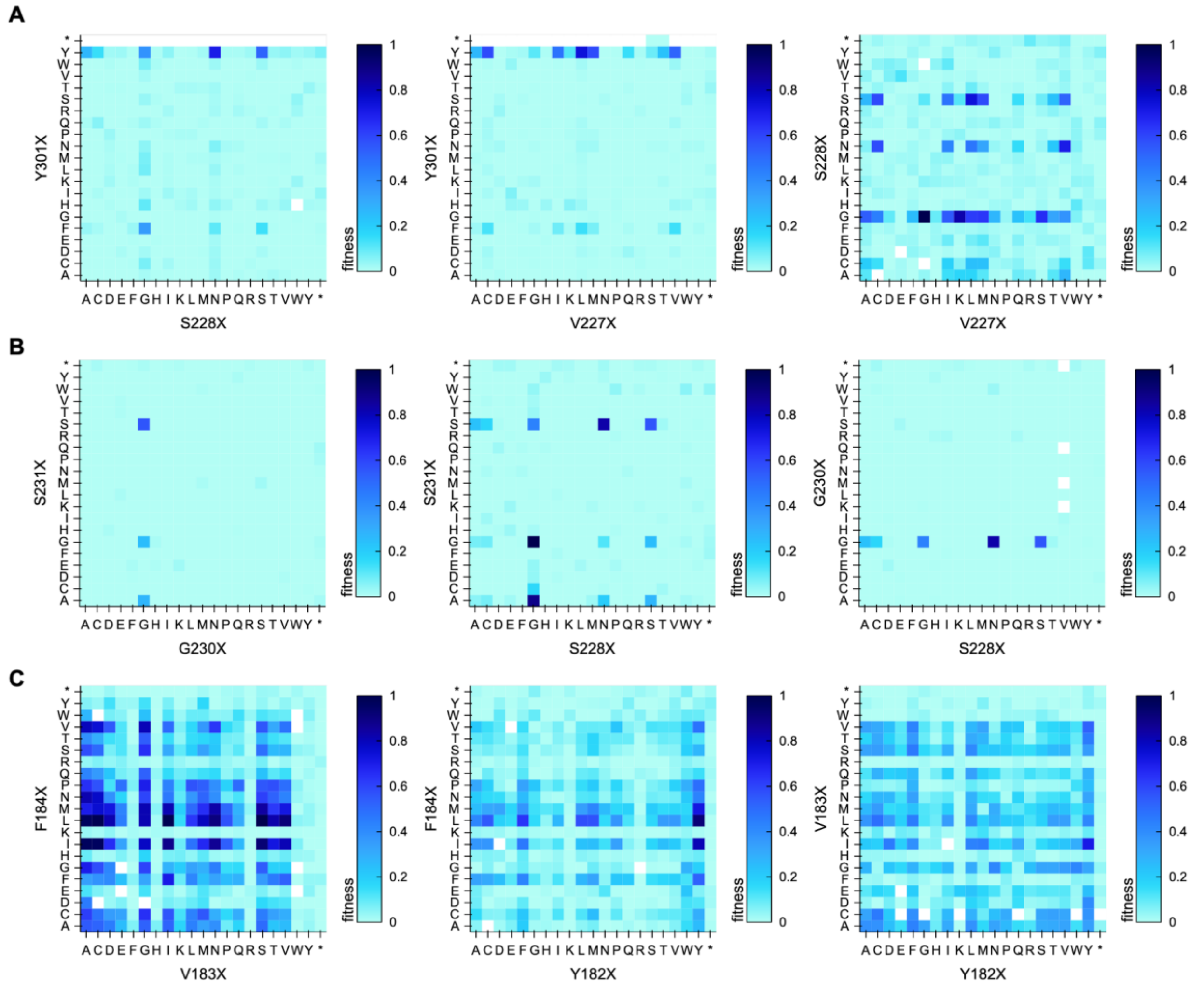

**Figure S10. Fitness for all pairs of residues for libraries G, H, and I.** Initial investigations into the utility of these sets of positions for larger libraries. For each plot, the unplotted residue was held at the parent amino acid at that position. Missing values appear white. **A** Library G separated into all three possible pairs of residues: 228 & 301, 227 & 301, and 227 & 228. Once again Y301 appeared nearly mandatory. Positions 227 and 228 appeared to allow many more residues. One especially exciting observation was that G227 was highly activating when paired with G228, but in the original S228 background it ablated activity. **B** Library H separated into all three possible pairs of residues: 230 & 231, 228 & 231, and 228 & 230. Very few amino acids were accepted at positions 230 and 231 while the range of those accepted at position 228 appeared similar to what was observed in library G. **C** Library I separated into all three possible pairs of residues: 183 & 184, 182 & 184, and 182 & 183. This library had many reasonably active variants, with some of the biggest improvements coming from the 183/184 pairing. Y182 appeared to be reasonably important for activity, but Phe, Leu, and Met were accepted at 182 to varying degrees with different residues at 183 and 184.

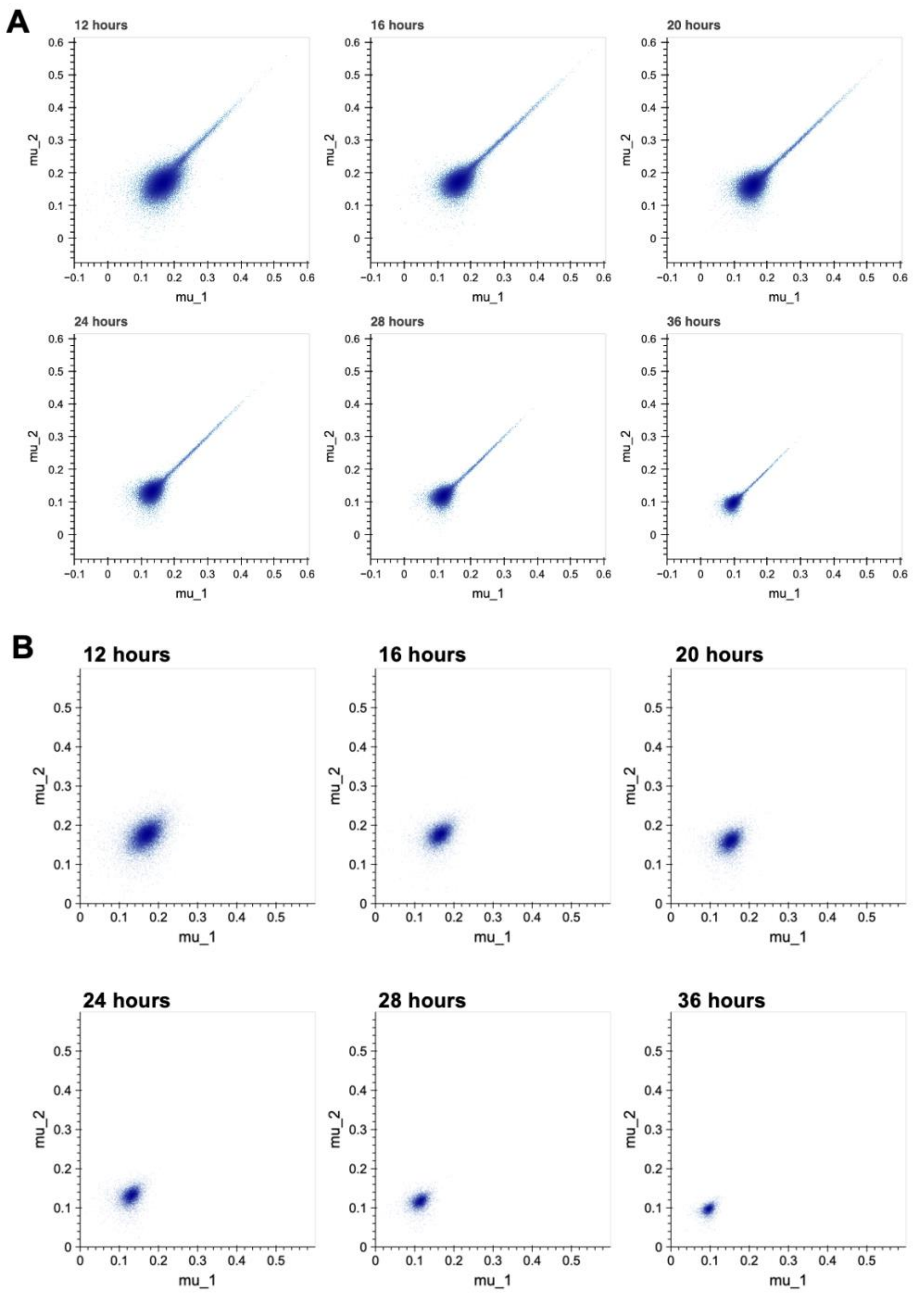

**Figure S11. Correlation of μ scores between replicates for 4-site data and stop-codon sequences.** Mu values were calculated as described by Kowalsky et al. (15) for all variants at all timepoints. **A** Panel of six plots showing mu for replicate 1 plotted against mu for replicate 2 for all variants at all timepoints. **B** Panel of six plots showing mu for replicate 1 plotted against mu for replicate 2 for just the stop codon-containing sequences at all timepoints. We observed that the distribution shrunk over time for both all variants as well as for stop codons, leading us to subtract the average mu for the stop codons within each replicate and timepoint and divide by the maximum mu value within that replicate and timepoint (mu_1-bg/max or mu_2-bg/max).

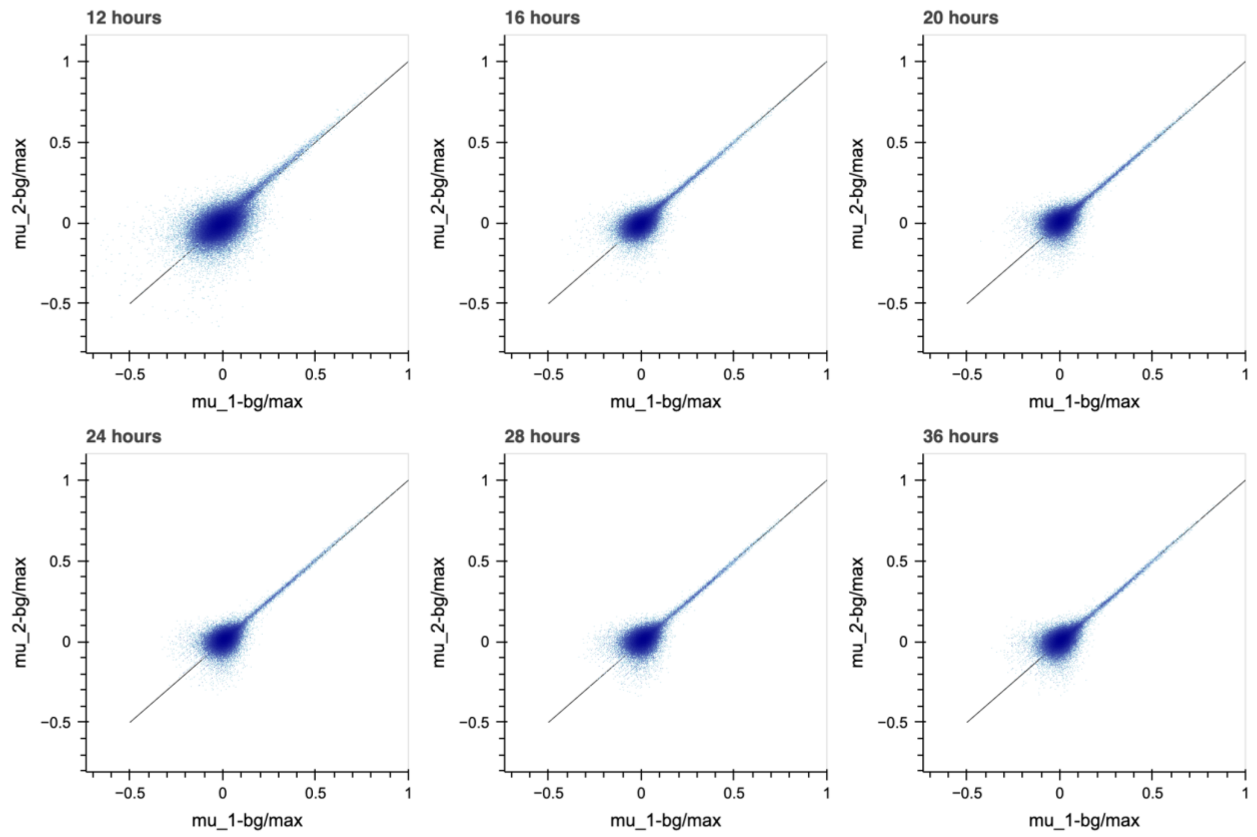

**Figure S12. Correlation of fitness scores between replicates for all timepoints.** The scaled mu values for both replicates plotted against each other by timepoint overlaid on the identity line, y=x. There was a high degree of agreement between the replicates as well as between the timepoints.

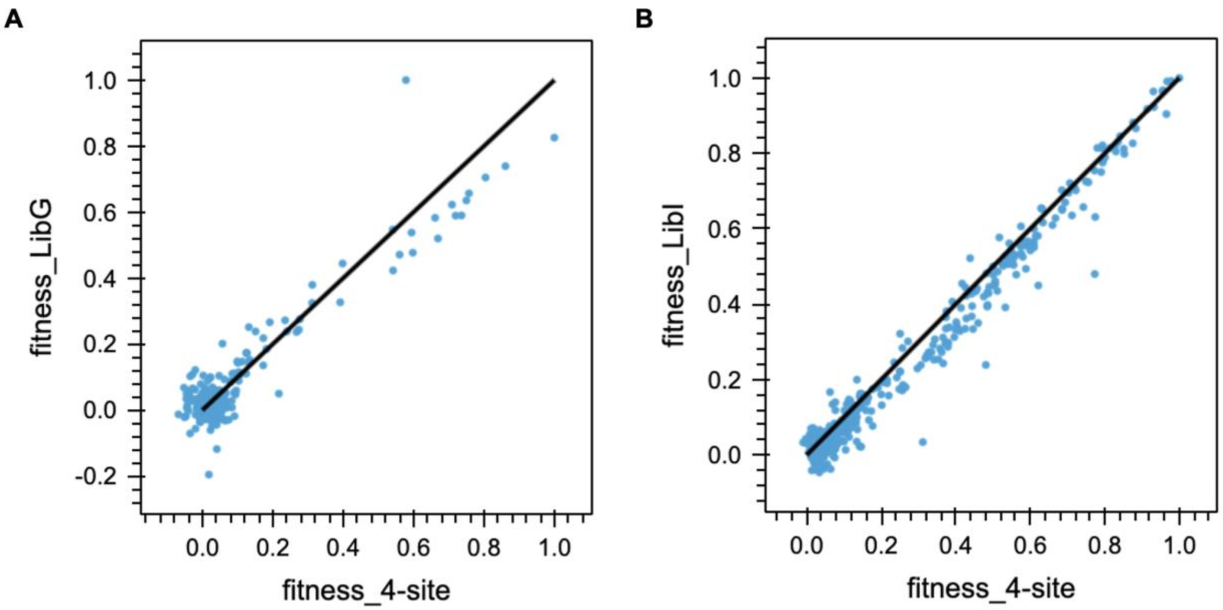

**Figure S13. Comparison of fitness values from subset 3- and 4-site libraries**. Library G (sites 227, 228, and 301) and Library I (sites 182, 183, 184) both contained sites found in the larger 4-site library (183, 184, 227, 228). Fitness data for the shared variable positions was compared by normalizing to the maximum within the subset and plotting the fitness values for the 3-site libraries against the 4-site library. The black lines represent y=x. **A** Fitness data for sequences containing Y301 from Library G plotted against sequences containing V183 and F184 from the 4-site library. **B** Fitness data for sequences containing Y182 from Library I plotted against sequences containing V227 and S228 from the 4-site library.

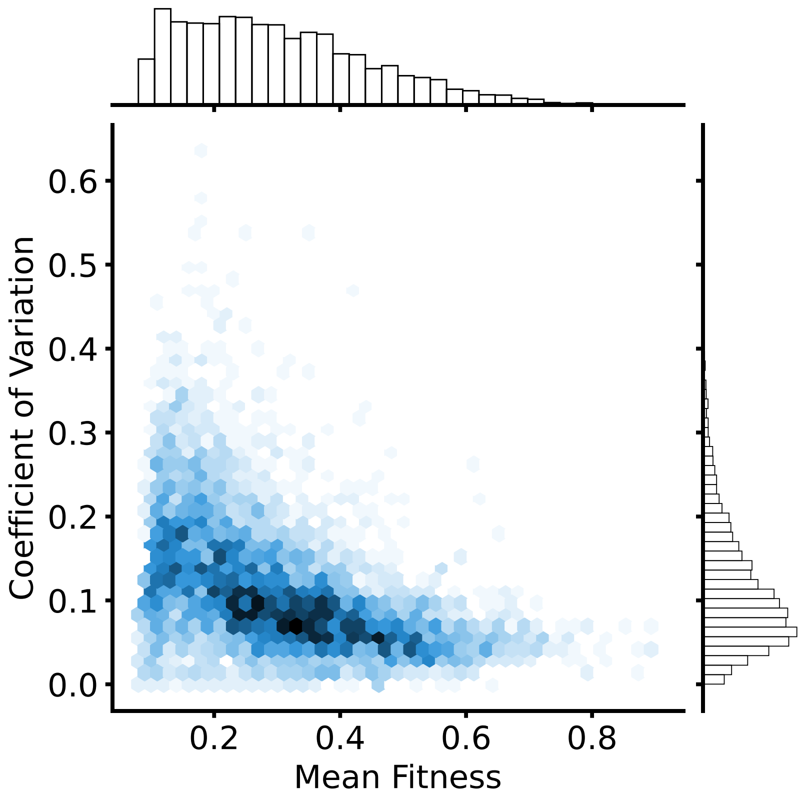

**Figure S14. Deviation from aggregate amino acid fitness for unique DNA sequences.** For a given amino acid sequence, variance of fitness across different DNA sequences is low. Mean fitness is defined as the mean fitness for all codons encoding the same amino acid sequence. The coefficient of variation is the standard deviation of fitness across codons encoding the same amino acid sequence, divided by the mean fitness for those codons. Coefficient of variation was calculated for each unique amino acid sequence of an active variant. Hex color indicates the density of points in that region. For most variants, the coefficient of variation is less than 0.2, suggesting that synonymous mutations do not have significant effects on fitness. The coefficient of variation also tends to decrease as fitness increases.

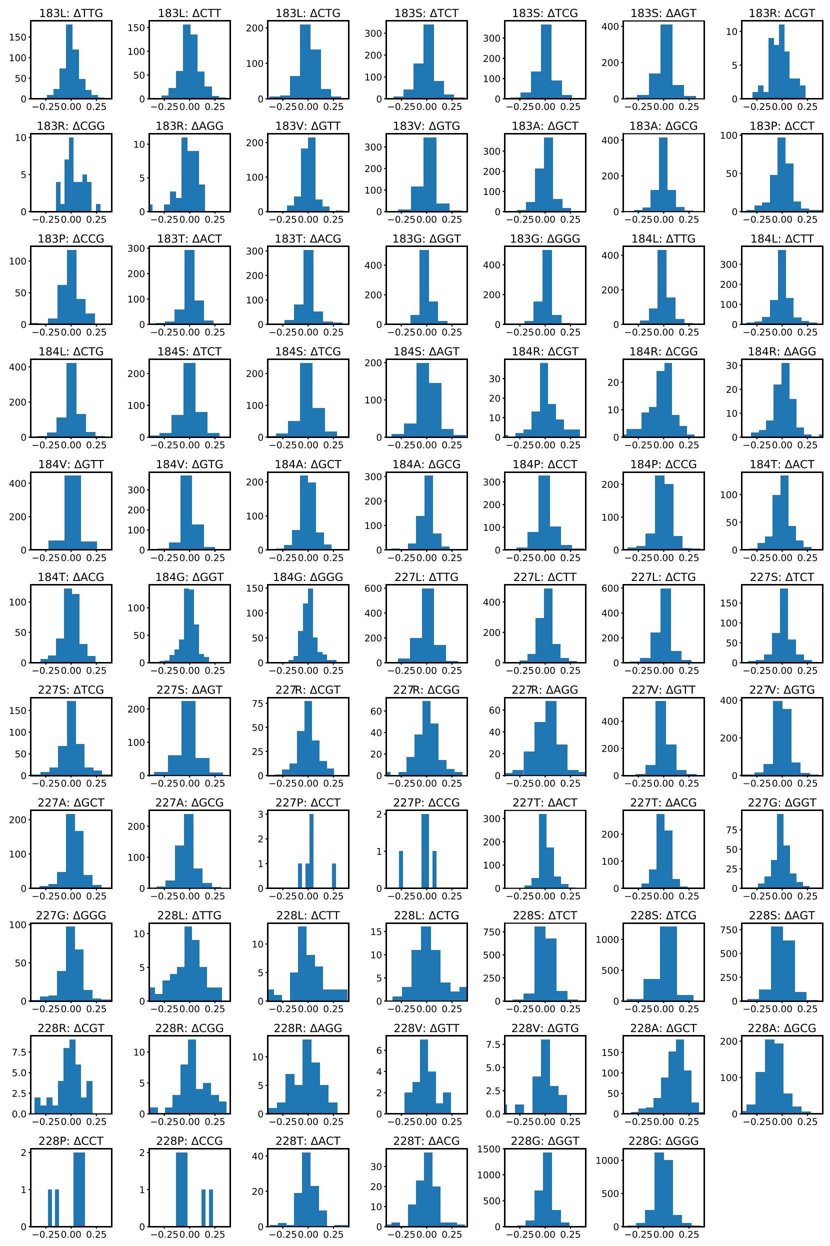

**Figure S15. Distribution of the fractional change in fitness for a specified codon and position.** Specific codons for a given amino acid and position generally do not have clear impacts on fitness. To examine this, we looked at only active variants and amino acid substitutions with synonymous mutations. For each subplot, we sampled all active sequences containing the specified amino acid at that position (e.g. 183L). We then sampled all codon sequences containing a specified codon mapping to that same amino acid (e.g. TTG at position 183). For each of the amino acid sequences, we first calculated the difference in fitness of using the specified codon at that position, compared to the mean fitness of sequences with any synonymous codon at the position. Then we divided this change by the average fitness of the sequence for all codons at that position. The distribution of these effects across all sampled sequences is plotted as a histogram where the Y-axis is the number of counts. Most distributions have a mean of zero, which suggests that specific synonymous mutations do not shift the distribution of fitness in obvious ways. Distributions have low standard deviations, which suggests that most synonymous mutations do not have significant effects on fitness.

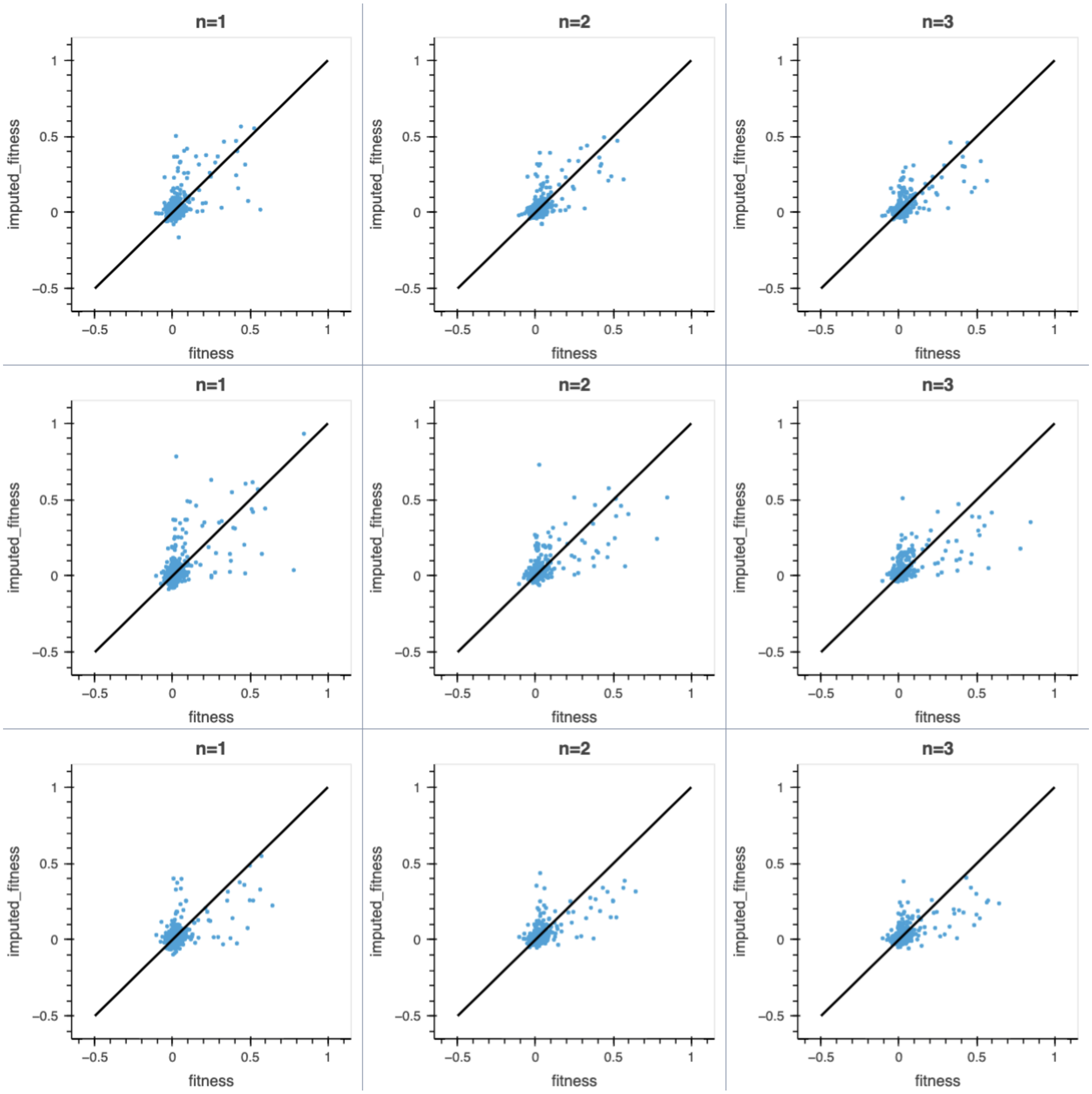

**Figure S16. Results of KNN imputation on 1000 randomly ablated fitness scores.** For three different sets of 1000 randomly ablated fitness scores, the KNN imputer was tested with weights='distance' and n_neighbors = 1, 2, or 3. The imputed fitness values are plotted against the actual fitness values and overlaid with the line y=x. The ablated fitness values were restored and then the KNN imputer with n_neighbors = 2 was chosen to impute the 871 missing fitness values for the TrpB four-site landscape.

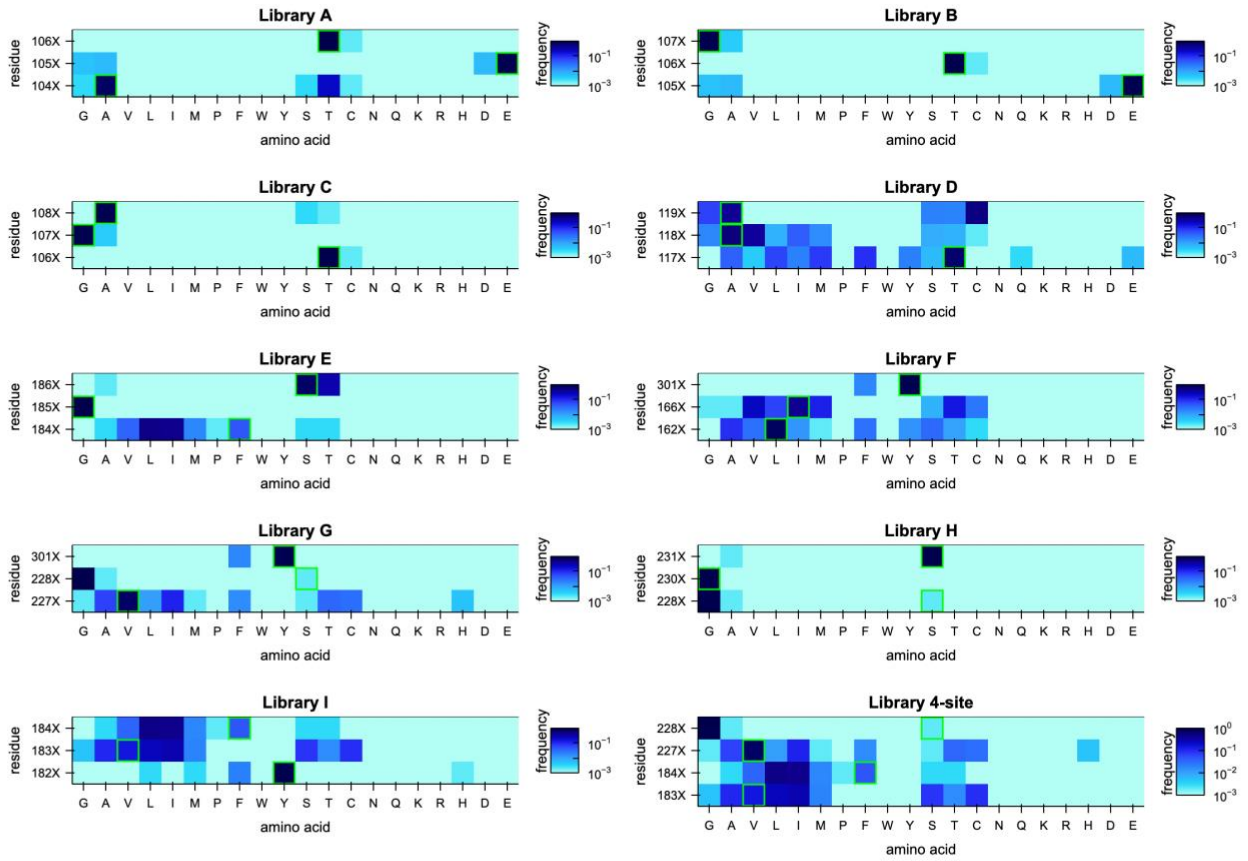

**Figure S17. Frequency of amino acids at each position for all 3- and 4-site landscapes.** EVcouplings was run with the webserver available at https://evcouplings.org/ to generate the multiple-sequence alignment used to determine the amino acid frequencies at each position (16). These results are from the *Tm*9D8* sequence as an input and using a bitscore of 0.3. The boxed amino acid is the residue present in *Tm*9D8*.

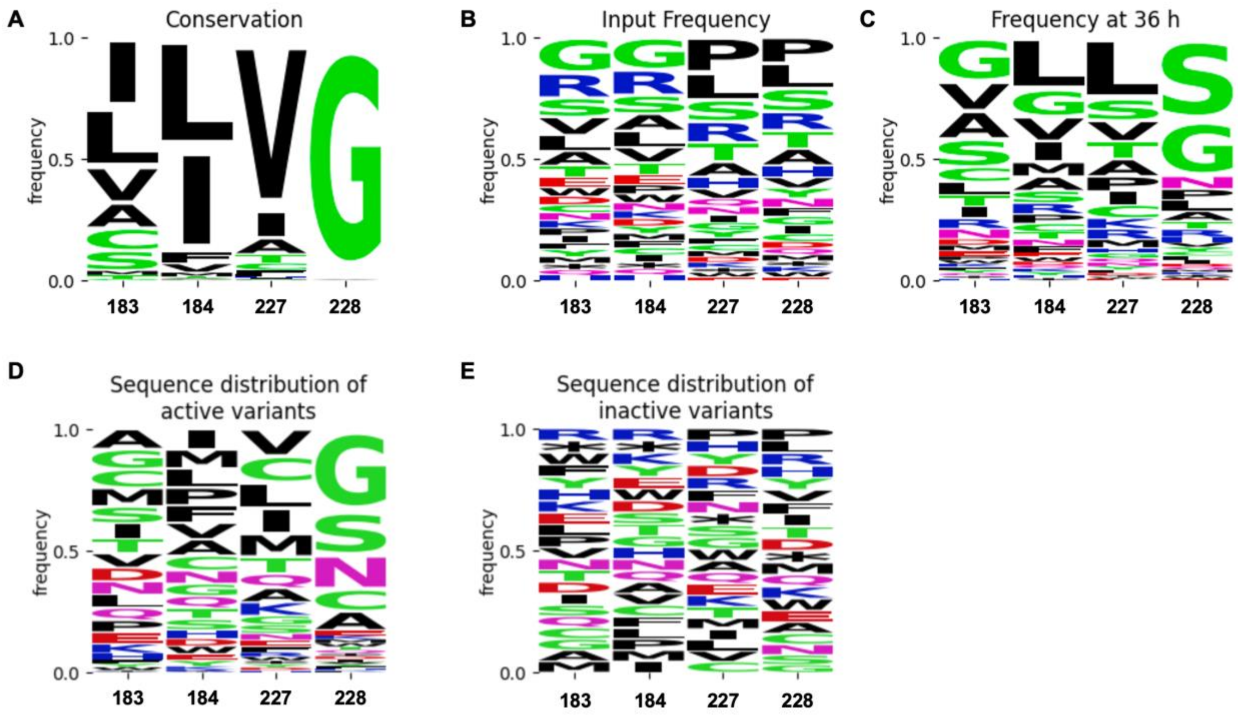

**Figure S18. Logo plots examining experimental and MSA results for the 4-site library**. For all plots, positions are decoupled for the analysis and frequency is calculated for each position independently. **A** Logo plot of frequency based on the MSA generated by EVcouplings using *Tm*9D8* as an input and a bitscore of 0.3. **B** Logo plot of the input frequency of each amino acid at each position for the input 4-site library. **C** Logo plot of the frequency of each amino acid at each position at the 36 h selection timepoint of the 4-site library. **D** Logo plot of the frequency of each amino acid at each position among variants classified as active. **E** Logo plot of the frequency of each amino acid at each position among variants classified as inactive.

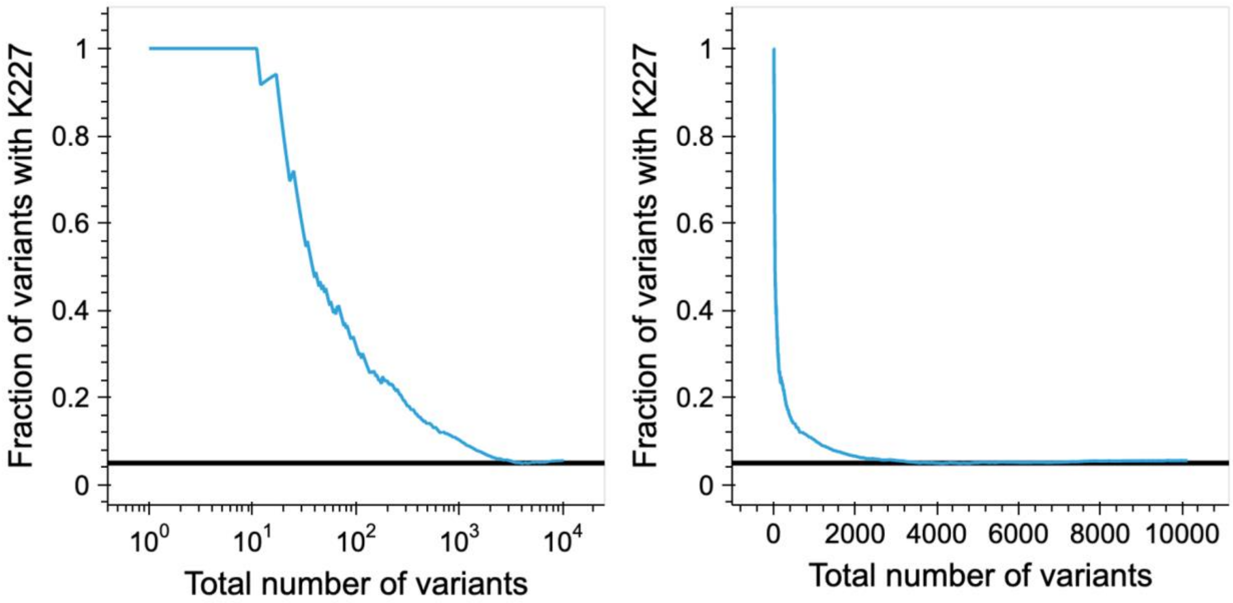

**Figure S19. Prevalence of K227 in the top sequences.** The fraction of variants containing K227 with that ranking or better versus the ranking of the variant. All ten top sequences contained K227 and even among the top ~2000 K227 remains overrepresented compared to the other nineteen possible amino acids. The black horizontal line is the expected fraction of sequences containing K227 (1/20) if sampling were random.

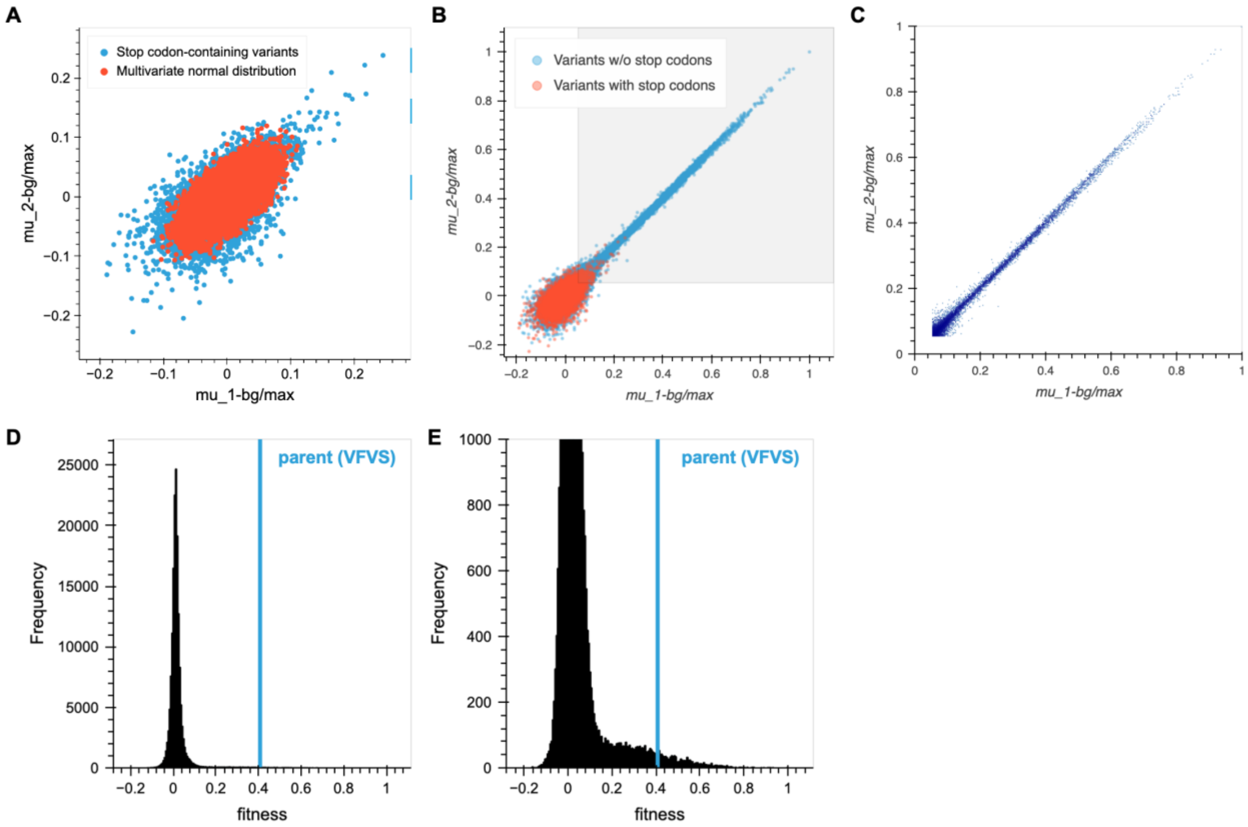

**Figure S20. Selecting an activity threshold for the 4-site landscape. A** Correlation between fitness for the two replicates for stop-codon containing sequences (blue). This was overlaid with points sampled from a multivariate normal distribution based on the stop-codon distribution (red). This shows that the distribution is roughly normal. **B** Correlation between fitness for the two replicates for all variants with stop-codon containing variants colored red and variants without stop codons colored blue. Since it appeared that the fitness distribution of stop codons was normal for each replicate, the activity threshold was imposed such that the fitness of a variant was at least 1.96 standard deviations above the mean fitness of all stop-codon-containing sequences for each replicate. The area of variants labeled "active" is colored in light gray, defined by mu_1-bg/max > 0.05397 and mu_2-bg/max > 0.05369. This threshold resulted in classification of 1.05% of the sampled stop-codon-containing sequences as active. **C** Data-shaded plot showing the correlation between fitness for the two replicates for variants without stop codons that were classified as active. **D** Histogram of the fitness values by averaging the two replicates. **E** Histogram of the fitness values calculated by averaging the two replicates with the maximum of the y-axis set to 1000. For both histograms the fitness of the parent variant, VFVS, is displayed with a blue, vertical line.

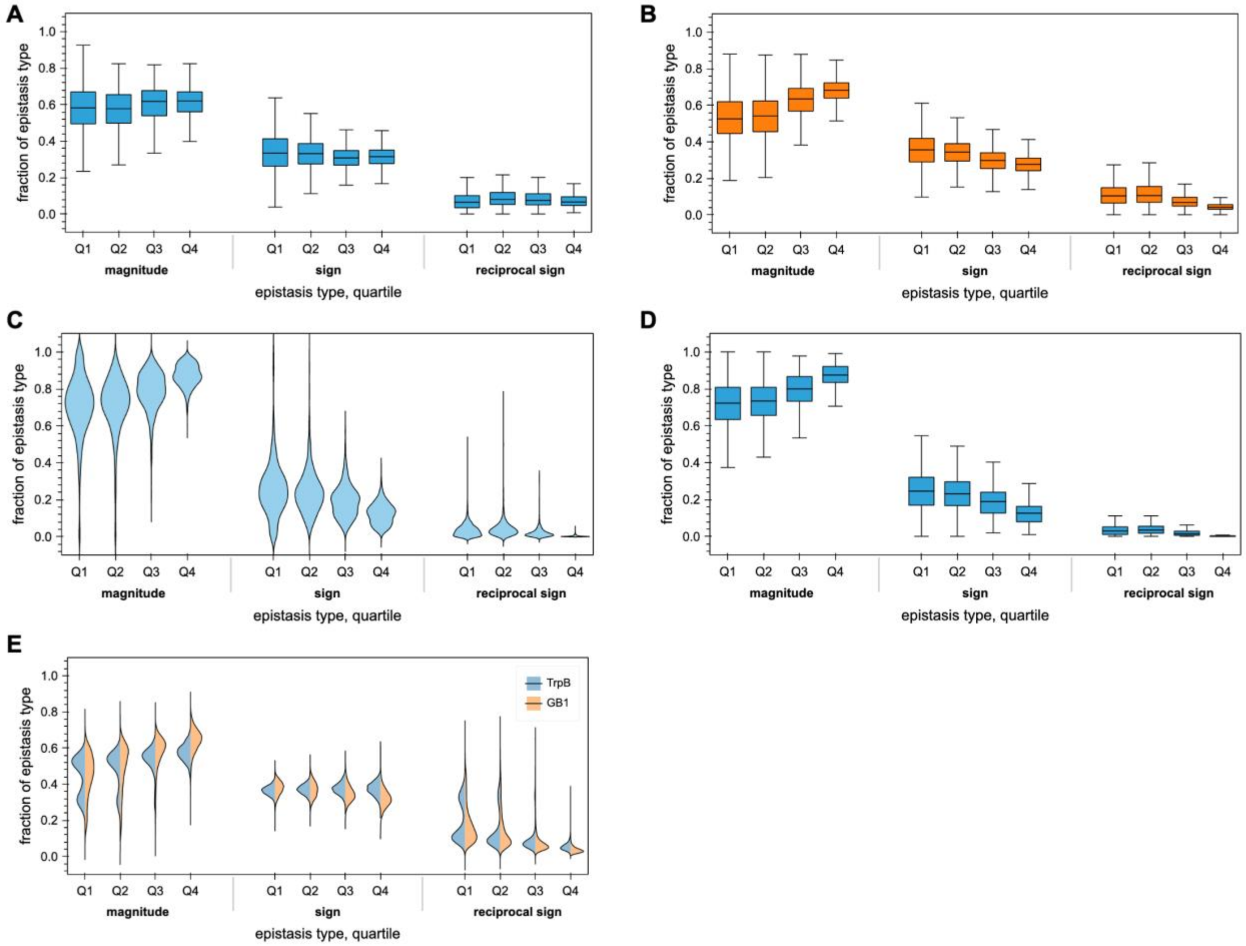

**Figure S21. Distribution of pairwise epistasis across quartiles. A** Distributions of the fractions of each epistasis type for TrpB as a box-and-whisker plot across fitness quartiles. **B** Distributions of the fractions of each epistasis type for GB1 as a box-and-whisker plot across fitness quartiles. **C** Distributions of the fractions of each epistasis type for the null model as a violin plot across fitness quartiles. **D** Distributions of the fractions of each epistasis type for the null model as a box-and-whisker plot across fitness quartiles. **E** Violin plot of how the distributions of epistasis change when enforcing final fitness > fitness to examine beneficial epistasis. Both the TrpB and GB1 landscapes appear to have an increase in magnitude epistasis with increasing fitness quartile and a reduction in reciprocal sign. The patterns diverge slightly for sign epistasis, where the fraction appears more constant for TrpB while it decreases for GB1.

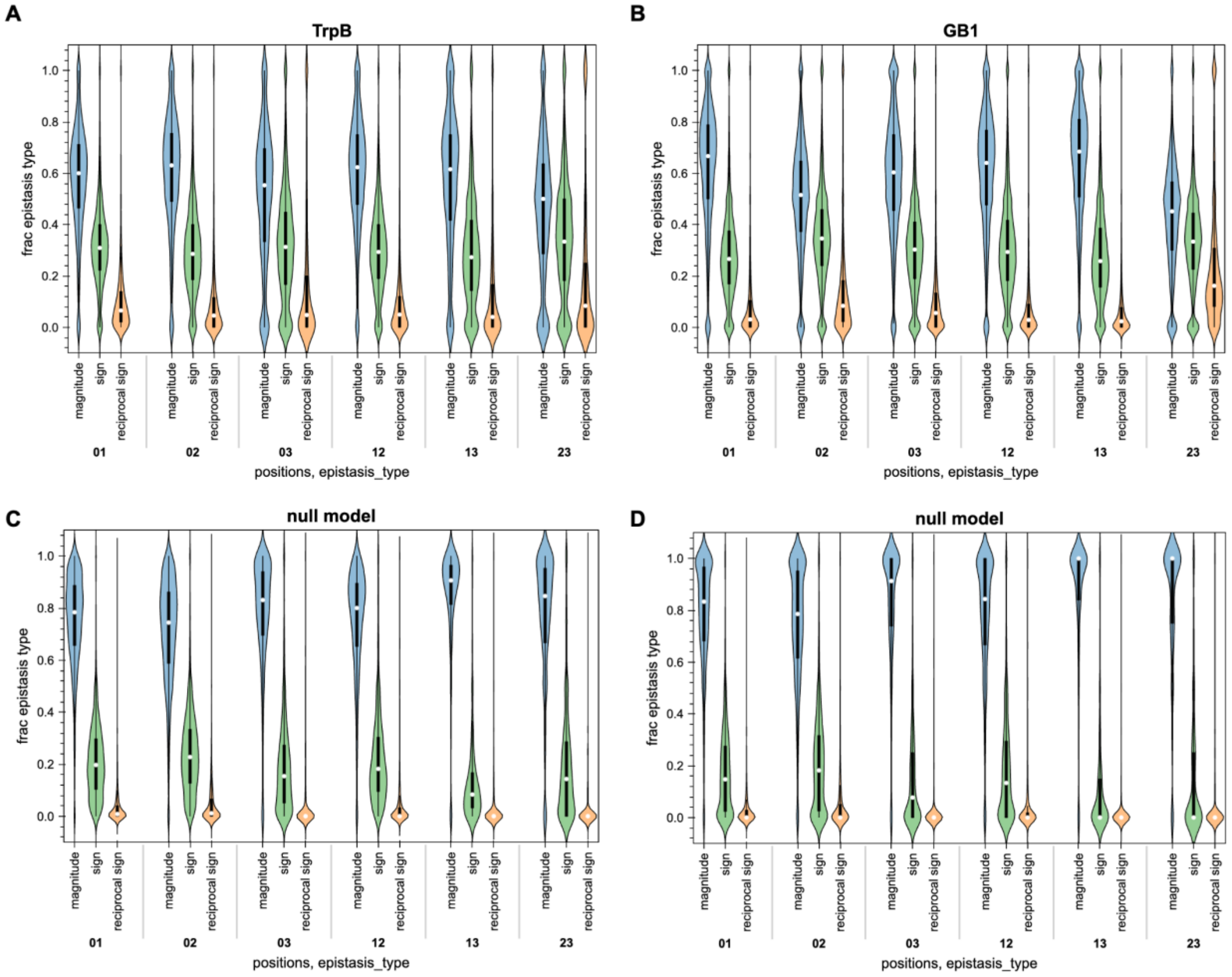

**Figure S22. Additional distributions of epistasis by position pair.** For all sequences above the respective activity thresholds, distributions of the fraction of each epistasis type grouped by pair of positions. **A** TrpB enforcing final fitness > initial fitness **B** GB1 enforcing final fitness > initial fitness **C** Null model **D** Null model enforcing final fitness > initial fitness. All variants within a set were required to be above the activity threshold to determine the epistasis type. In the null model, much of the epistasis is lost when it is enforced that final fitness > initial fitness, meaning there is less beneficial epistasis in this model. Differences appeared to be relatively minor for TrpB and GB1 with a slight increase in the amount of magnitude epistasis across all position pairs for both TrpB and GB1. Positions are listed in sequence order. TrpB: 0→183, 1→184, 2→227, 3→228. GB1: 0→39, 1→40, 2→41, 3→54.

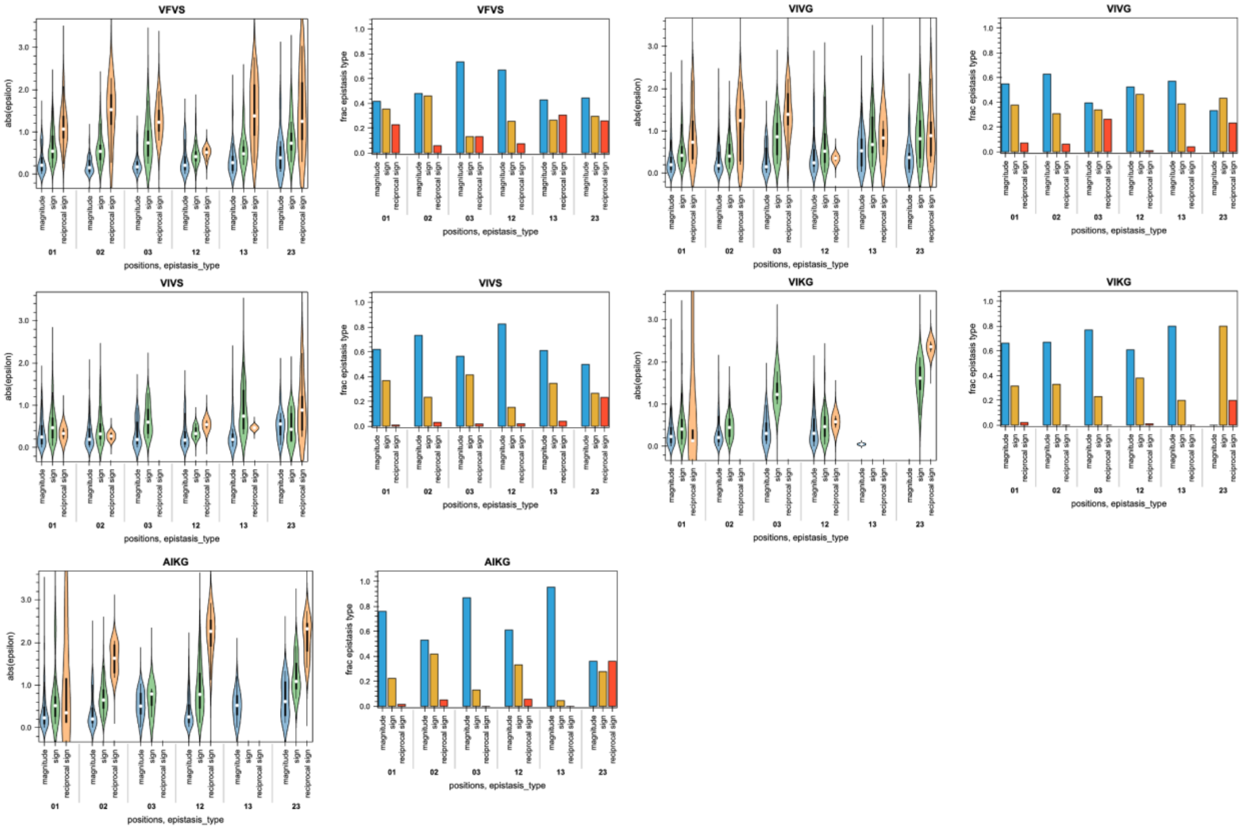

**Figure S23. Investigating the distribution of epsilon and epistasis type by variant.** For the five variants involved in the path from parent (VFVS) to the best variant (AIKG), we plotted the distribution of epsilon separated by epistasis type and position pair as well as the fractions of epistasis types for each position pair. Although we had seen that the position pairs showed similar fractions of each epistasis type when examining all variants, when examined one background sequence at a time there is a lot of variation. VFVS and VIVG appear to have reciprocal sign epistasis for all position pairs, while VIKG has almost none. There is also much more variation in the fractions of epistasis between each of the pairs of positions for each background. For example, VIVS exhibits very little reciprocal sign epistasis except between residue 227 (2) and 228 (3). For all sets, all variants were required to have fitness above the activity threshold, which is why some epsilon distributions are empty (no sets existed where all variants were above the threshold). Positions for TrpB and GB1 are listed in sequence order. TrpB: 0→183, 1→184, 2→227, 3→228. GB1: 0→39, 1→40, 2→41, 3→54.

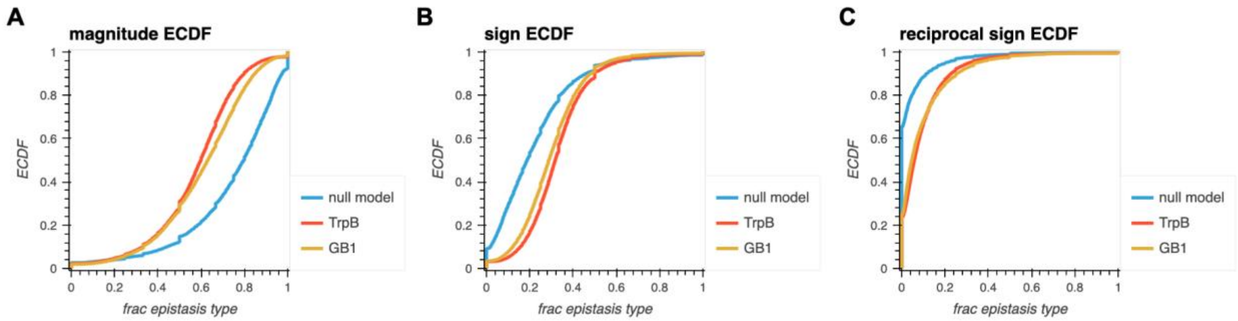

**Figure S24. Distributions of epistasis types as ECDFs overlaid with the null model.** For all sets of variants above the activity threshold we calculated a fraction of epistasis type for a given starting sequence. We then plotted the overall distributions of these fractions of epistasis type as ECDFs for TrpB, GB1, and the null model. **A** Magnitude epistasis **B** sign epistasis **C** reciprocal sign epistasis. There is a clear separation between the null model and the TrpB and GB1 landscapes, which are enriched in epistasis.

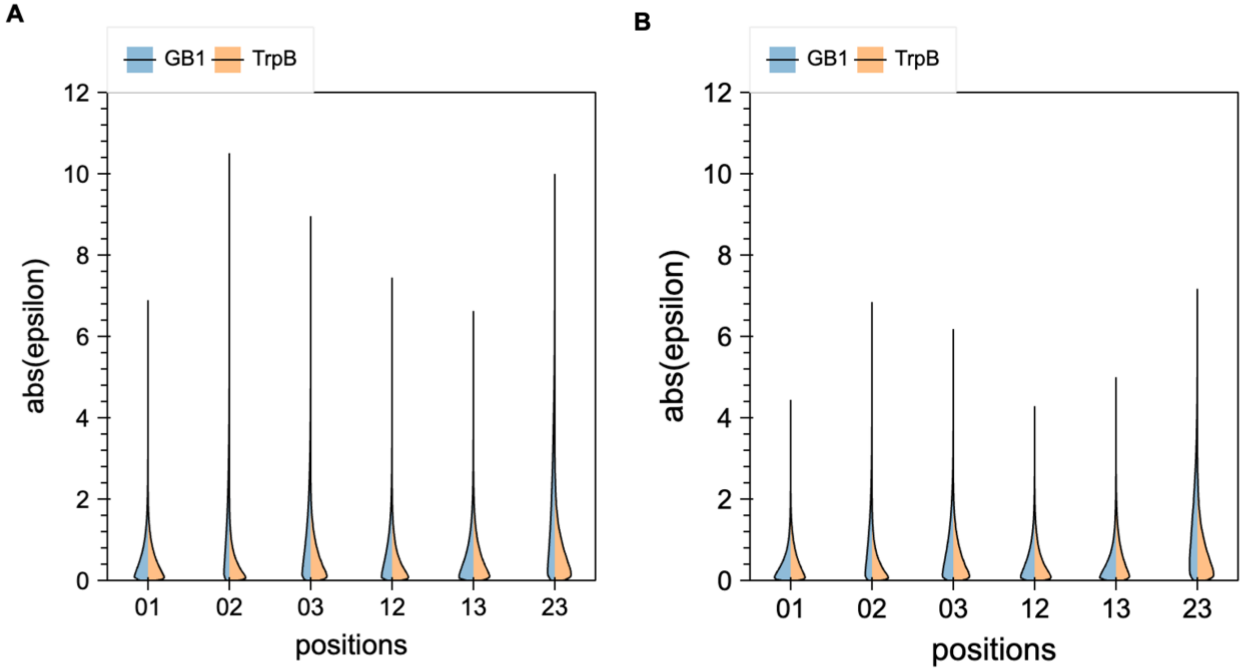

**Figure S25. Epsilon distribution by pair of positions for TrpB and GB1.** Epsilon was calculated as described in Olson et al. (17) for each set of variants: $\varepsilon = \ln\left(\frac{fit_{11}}{fit_{00}}\right) - \ln\left(\frac{fit_{01}}{fit_{00}}\right) - \ln\left(\frac{fit_{10}}{fit_{00}}\right)$ where 00 is the starting variant, 11 is the final variant with two substitutions, and 01 and 10 are the two single substitutions. **A** Epsilon distributions if all variants are required to be above the activity threshold. **B** Epsilon distributions if all variants are required to be in the top 9783 variants. Positions for TrpB and GB1 are listed in sequence order. TrpB: 0→183, 1→184, 2→227, 3→228. GB1: 0→39, 1→40, 2→41, 3→54.

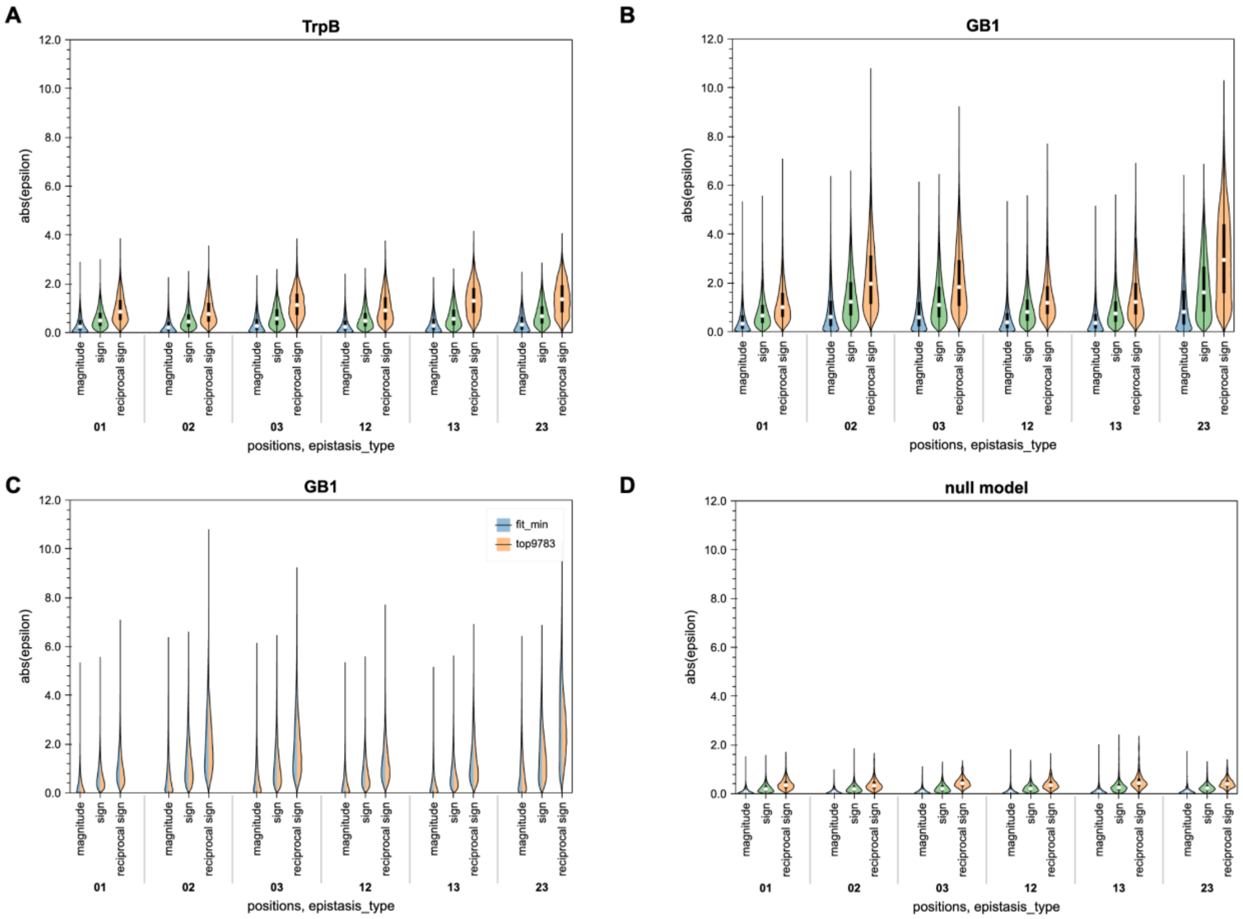

**Figure S26. Epsilon distributions by position pair and epistasis type. A** Epsilon distributions for TrpB for sets of variants where the fitness of every variant is above the activity threshold. **B** Epsilon distributions for GB1 for sets of variants where the fitness of every variant is above the activity threshold. **C** Epsilon distributions for GB1 where each variant in the set is either above the activity threshold (blue) or in the top 9783 variants (orange). For TrpB, this does not change the distribution, so it is displayed only for GB1. **D** Epsilon distributions for a null model for sets of variants where the fitness of every variant is above the activity threshold. Positions for TrpB and GB1 are listed in sequence order. TrpB: 0→183, 1→184, 2→227, 3→228. GB1: 0→39, 1→40, 2→41, 3→54.

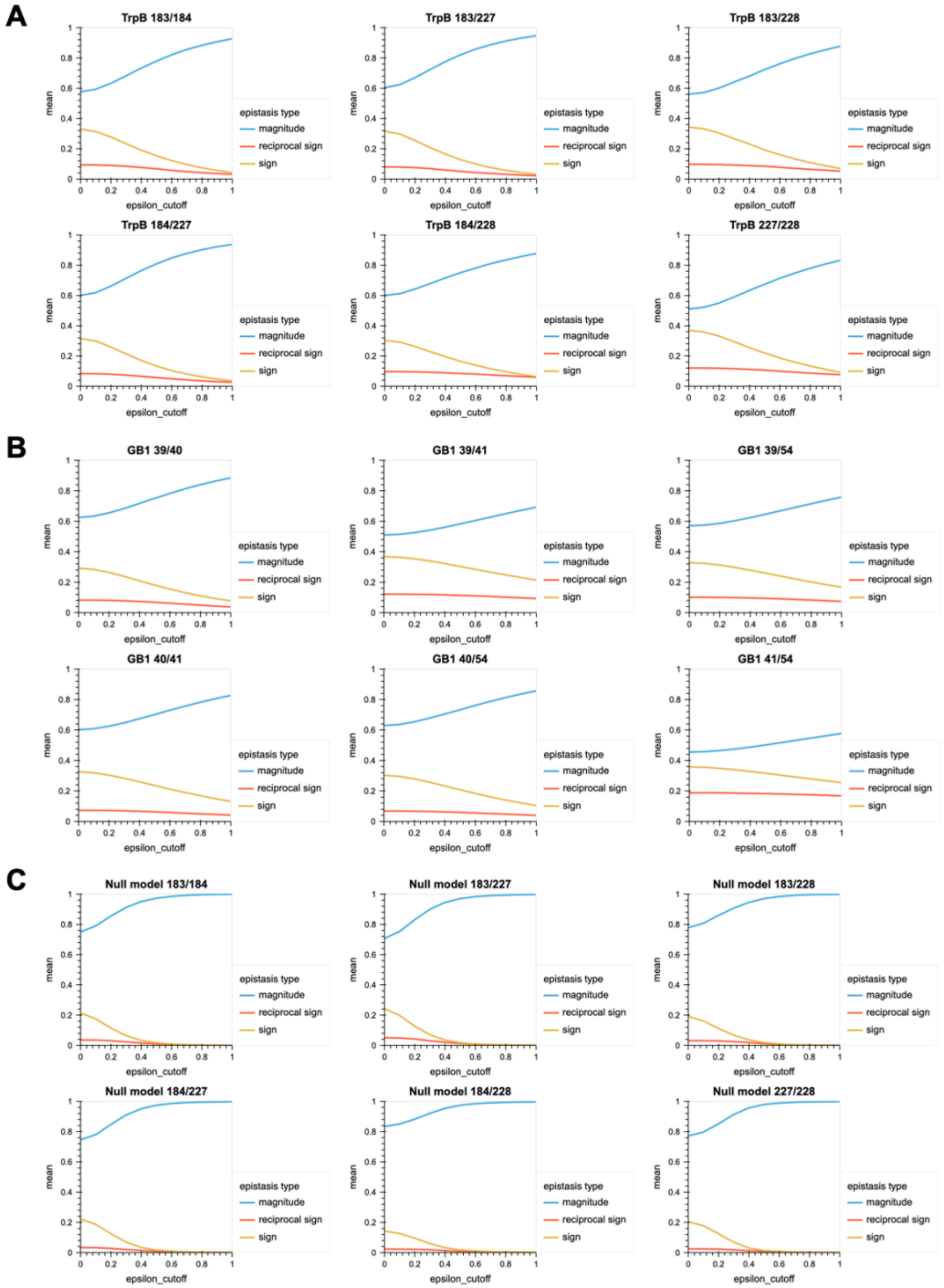

**Figure S27. Fraction of epistasis type as a function of an epsilon threshold.** The magnitude of epistasis in a landscape is an important attribute for characterizing the strength of the effects. For this analysis, we plot the mean fraction of each epistasis type for each pair of positions across TrpB, GB1, and the null model as a function of an epsilon threshold, filtering out smaller epistatic effects as epsilon increases. **A** For TrpB, the fraction of magnitude epistasis ranges from ~0.5–0.6 initially and reaches around ~0.8–0.9 by epsilon=1. **B** For GB1, the fraction of magnitude epistasis ranges from 0.45–0.6 initially and reaches around 0.6–0.85 by epsilon=1. **C** For the null model, the fraction of magnitude epistasis ranges from 0.7–0.85 initially and reaches 1 for all positions by the time epsilon≈0.8. This indicates that the strength of the epistatic effects ranks as follows: GB1>TrpB>>null model.

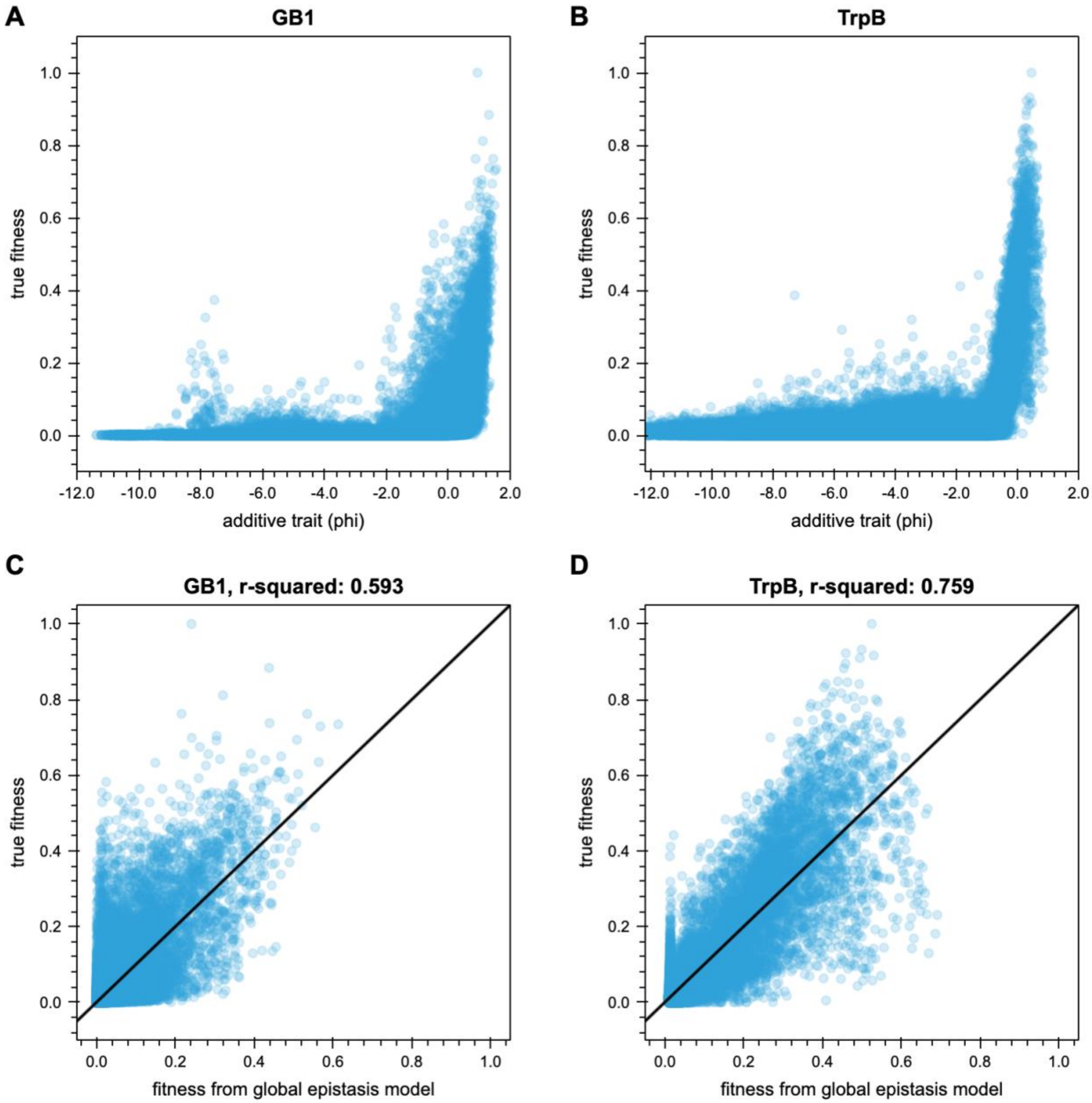

**Figure S28. Global epistasis of the TrpB and GB1 landscapes.** Using the software provided by Otwinowski et al. (18), we analyzed the global epistasis within the TrpB and GB1 landscapes using a tolerance of $10^{-12}$ and $10^{-14}$ respectively. The solid black line represents the linear fit. **A** The true fitness values of the GB1 landscape plotted against the additive trait, $\Phi$, an inferred additive trait depending on genetic sequence ($\phi = \beta_0 + \sum_i^L \beta_{i,a_i}$, where $L$ is the number of sites, $\beta_0$ is the parent phenotype, and $\beta_{i,a_i}$ is the effects of substitutions for each position $i$ and amino acid at that position $a_i$). **B** The true fitness values of the TrpB landscape plotted against the additive trait, $\Phi$. **C** The true fitness values of the GB1 landscape plotted against the fitness values arising from a best-fit model of global epistasis ($y = g(\phi) + \varepsilon$, where $g(\phi)$ is a nonlinear function mapping the additive trait to the data and $\varepsilon$ is noise). **D** The true fitness values of the TrpB landscape plotted against the fitness values from the global epistasis model. We observe that the global epistasis model captures a large amount of variation in fitness, but it is far from perfect. Many of the highest fitness variants are predicted to be of intermediate fitness under the global epistasis model, and for TrpB, many intermediate fitness variants are predicted to be better than they actually are.

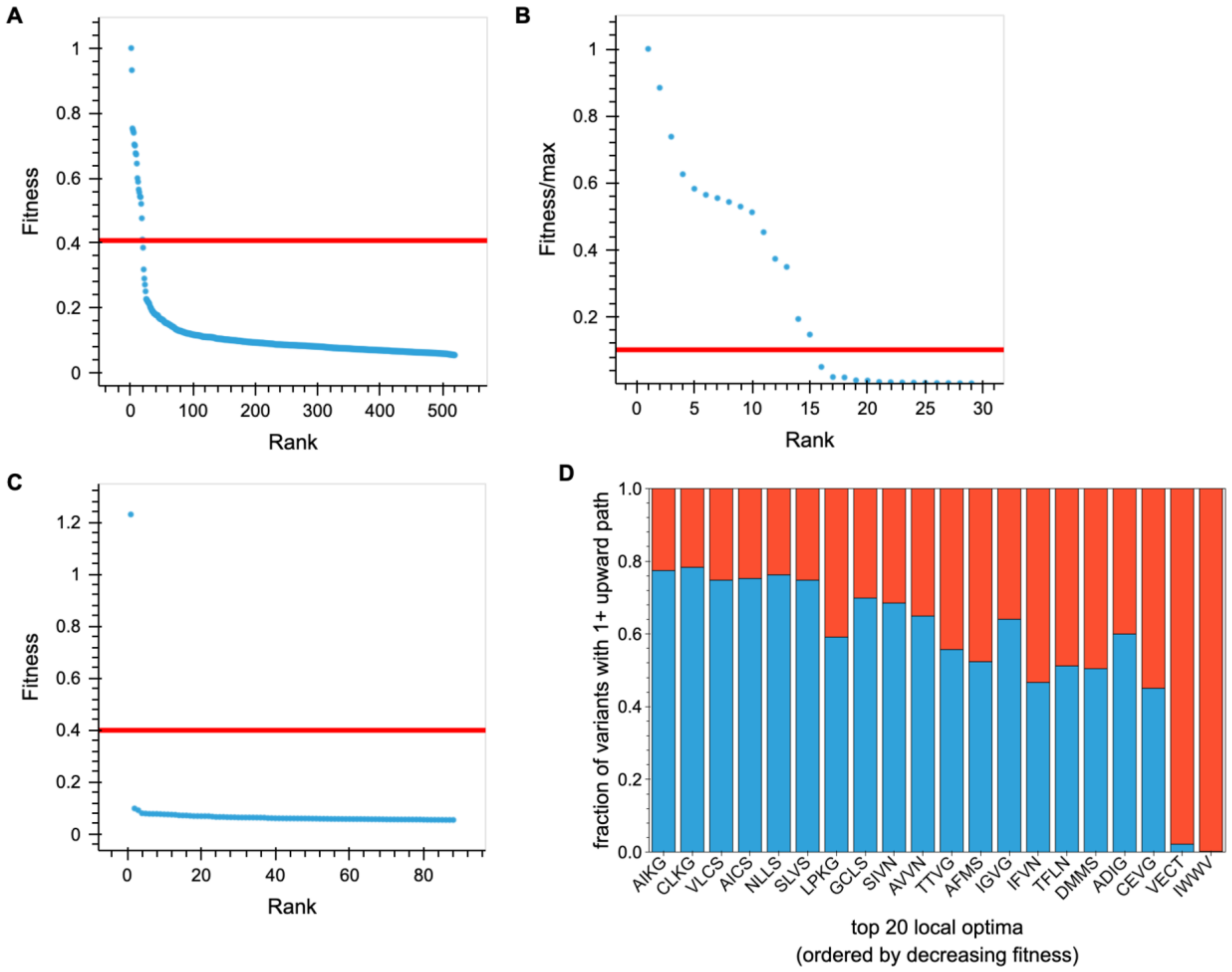

**Figure S29. Fitness and accessibility local optima. A** Fitness of each local optima in the TrpB landscape versus rank based on fitness. The red line denotes the fitness of the parent variant, VFVS, and the top variant, with fitness=1, is AIKG. **B** Fitness of each local optima in the GB1 landscape versus rank based on fitness. The red line denotes the fitness of the parent variant, VDGV, and the top variant, with fitness=1, is AHCA. **C** Fitness of each local optima in the null landscape versus rank based on fitness. The red line denotes the fitness of the parent variant, VFVS, and the top variant, with fitness=1.23, is ILLN. **D** Fraction of active variants in the TrpB landscape with at least one upward path to each local optima. Allowing no decreases in fitness, this is the fraction of starting variants that have at least one upward path to each of the top 20 local optima (blue). The fraction of starting variants that do not have even one upward path to the local optima is in red. There is a general trend that as the fitness of the local optima decreases it becomes less accessible, likely because some of the paths require variants with higher fitness than the optima, which makes them inaccessible. However, many of these local optima are still similarly accessible as AIKG, meaning they could trap evolution campaigns.

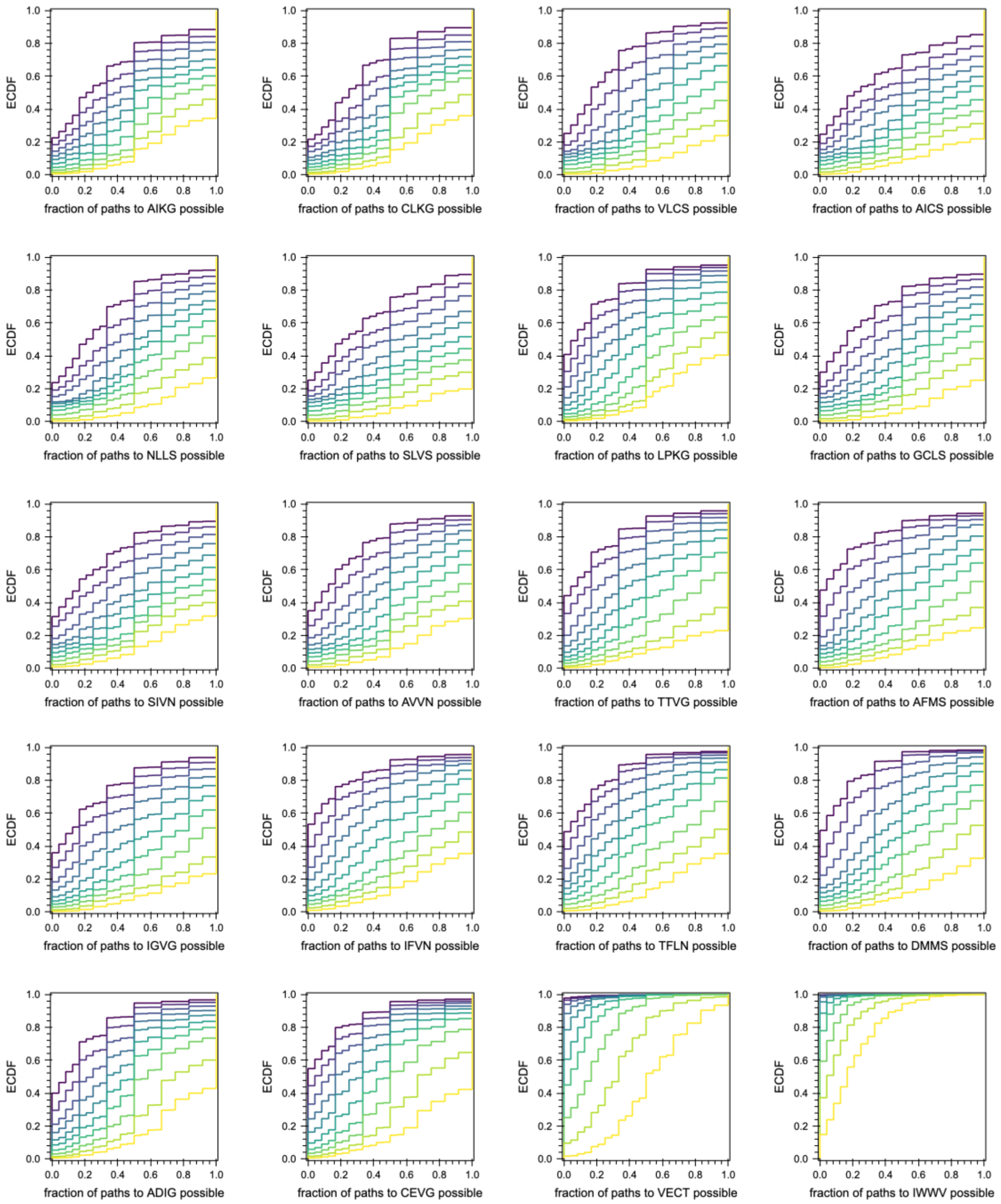

**Figure S30. Analysis of paths to each local optimum.** Empirical cumulative density functions (ECDFs) of active variants with at least one upward path to each of the top twenty local optima given the allowed fitness decreases. Most ECDFs are similar to that of AIKG with varying widths of distributions, indicating similar levels of accessibility. This means that evolution campaigns could be easily caught in these local optima. The final two optima presented, VECT and IWWV, are much less accessible.

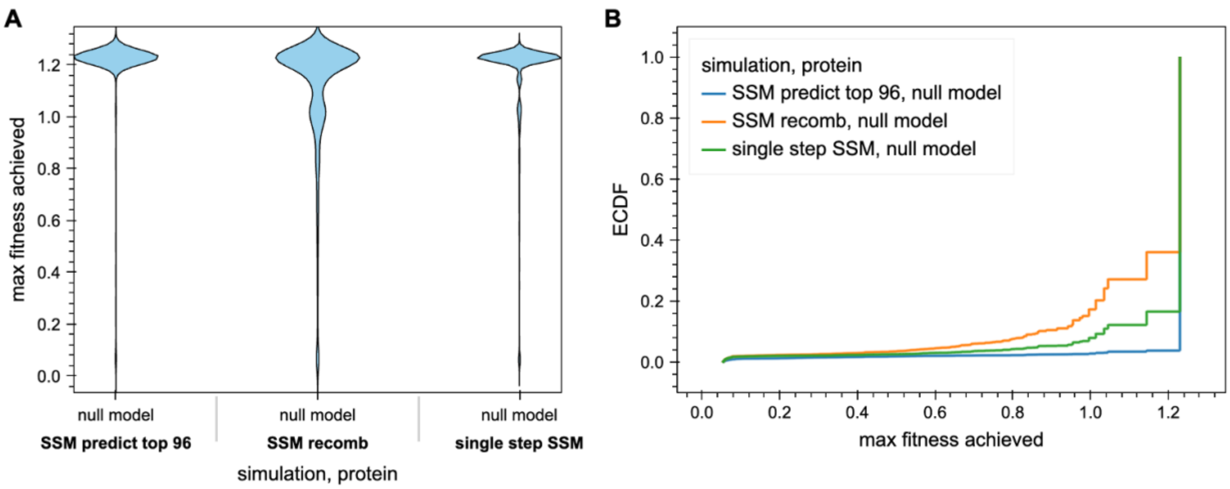

**Figure S31. Performance of DE methods for the null model.** Results of the directed evolution simulations run starting from any variant above the fitness minimum derived from the original TrpB landscape. **A** Violin plots of the distributions of the maximum fitness achieved show very similar performance for all methods with many trajectories reaching the fitness maximum (1.23). **B** ECDFs of the distributions of the maximum fitness achieved show that the same order of performance holds for the null model as for TrpB and GB1 (Method 3 > Method 2 > Method 1), although all three methods do significantly better. Results are summarized in Table S15.

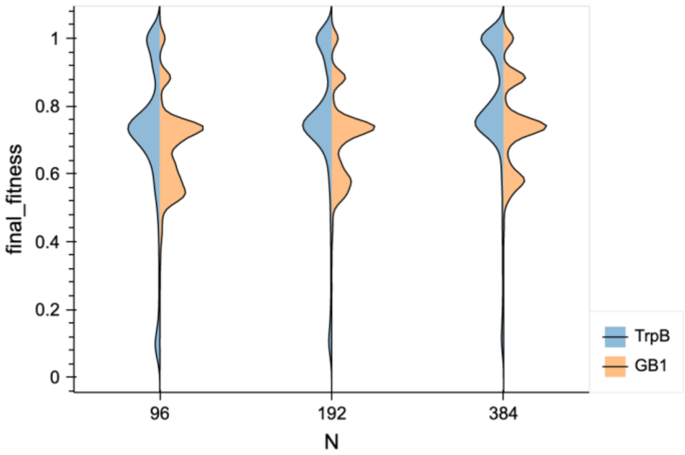

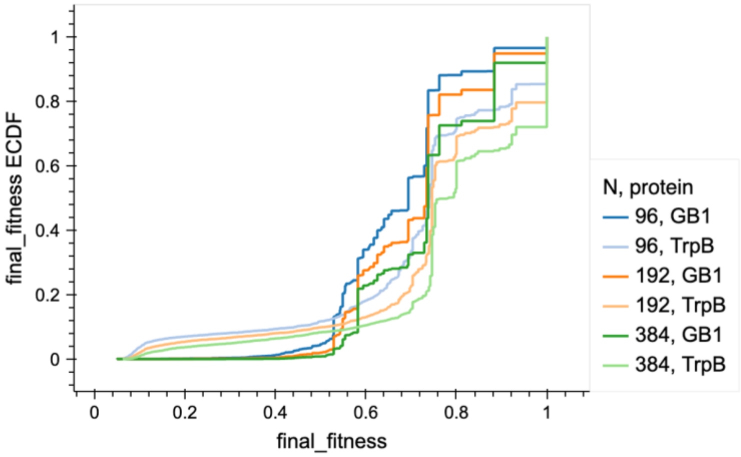

**Figure S32. Varying the number of exact variants tested in SSM predict top *N*.** Since many machine learning-assisted directed evolution approaches require direct synthesis of *N* variants for a prediction round, we tested if increasing *N* improved the max fitness achieved via choosing top variants with SSM and recombination based on additivity. Testing more variants did shift the distributions slightly upward, but the return was quite small for a 2X or 4X increase in required synthesis costs.

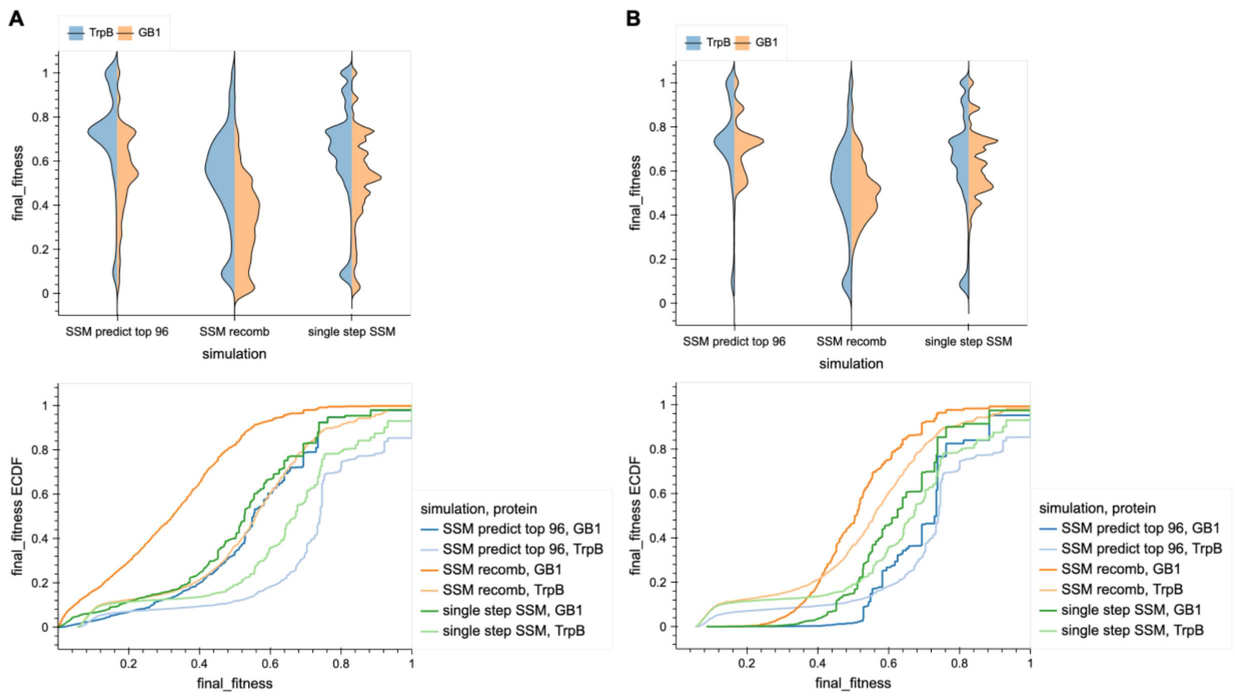

**Figure S33. Comparing simulation results with varying activity cutoffs for GB1 data.** Presented in the main text, we show the simulation results starting from each of the top 9783 non-imputed variants. We observed significant differences in the performance based on the starting cutoffs imposed for GB1 that made comparison between GB1 and TrpB difficult. **A** A comparison of the directed evolution simulation results starting from all variants above the respective activity thresholds for TrpB and GB1. In these results, the maximum fitness achieved is generally much higher for TrpB. However, the minimum fitness allowed for TrpB is ~1/20 the max while for GB1 it is ~1/1000 the max. This means a given starting point for TrpB is fewer fold-improvements from the maximum to begin with. **B** Alternatively, the simulations can be run for GB1 using the same fraction of the maximum fitness as a cutoff. In this case, only 5587 variants from the GB1 landscape are within this cutoff. This makes the simulation results between TrpB and GB1 much more similar, with GB1 having fewer campaigns trapped at very low fitness (~0–0.3), but more trapped and medium fitness (~0.3–0.8).

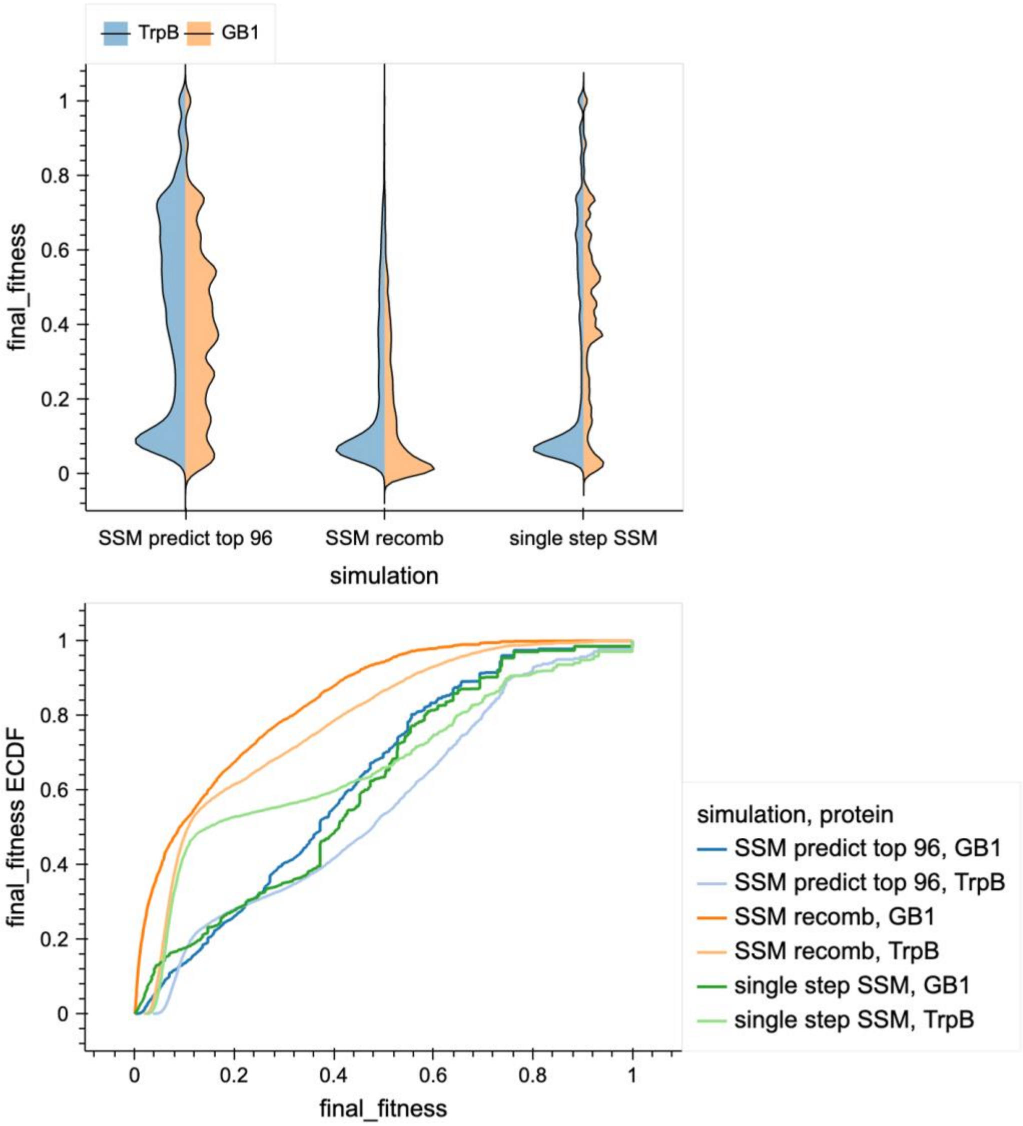

**Figure S34. Directed evolution simulations from any variant with fitness >0 in the landscape.** As a final comparison, we tested how well simulations did starting from any random variant in the landscape of either TrpB or GB1. As expected, the performance was significantly worse, especially for site-saturation mutagenesis + recombine best (middle violin). Single-step site saturation and site-saturation predict top 96 performed similarly for GB1, but single-step SSM was by far the best for TrpB, likely because it allowed escape from the distribution of inactive variants. This suggests that a single-step walk may be a more robust approach, especially for early evolution campaigns when activity is near-noise levels even though SSM predict top 96 did somewhat better when starting from the detectably active variants.

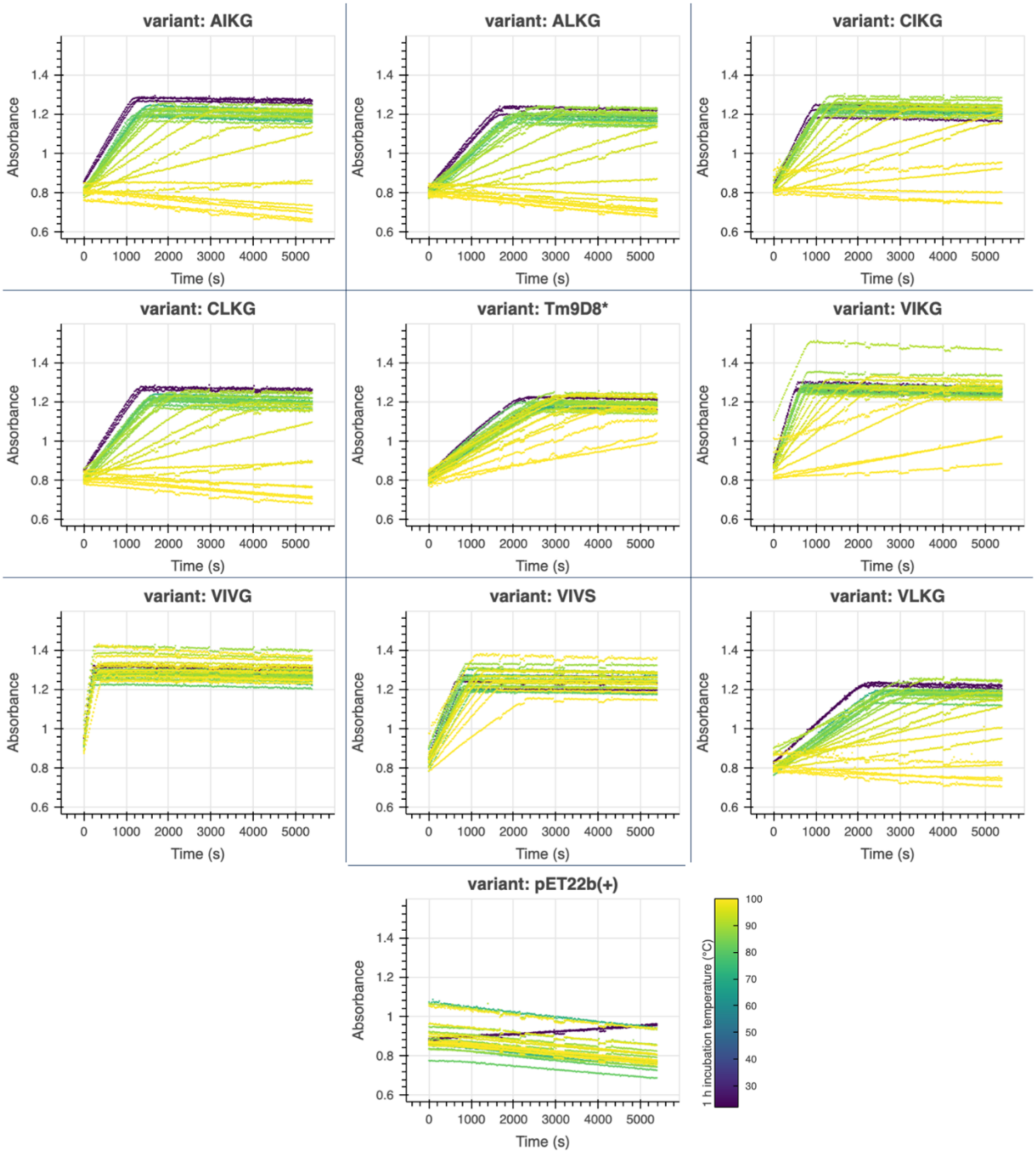

**Figure S35. $T_{50}$ absorbance vs time (s) plots colored by temperature for each variant.** Across three different collection plates, lysate harboring each variant was incubated for 1 h at a temperature between room temperature and 99 °C. Lysate was spun down and then added to a reaction mix containing indole and serine, and absorbance at 290 nm was collected over time. It is clear that some variants lose activity with temperature while others barely change (VIVG). The initial rates were captured with linear fits.

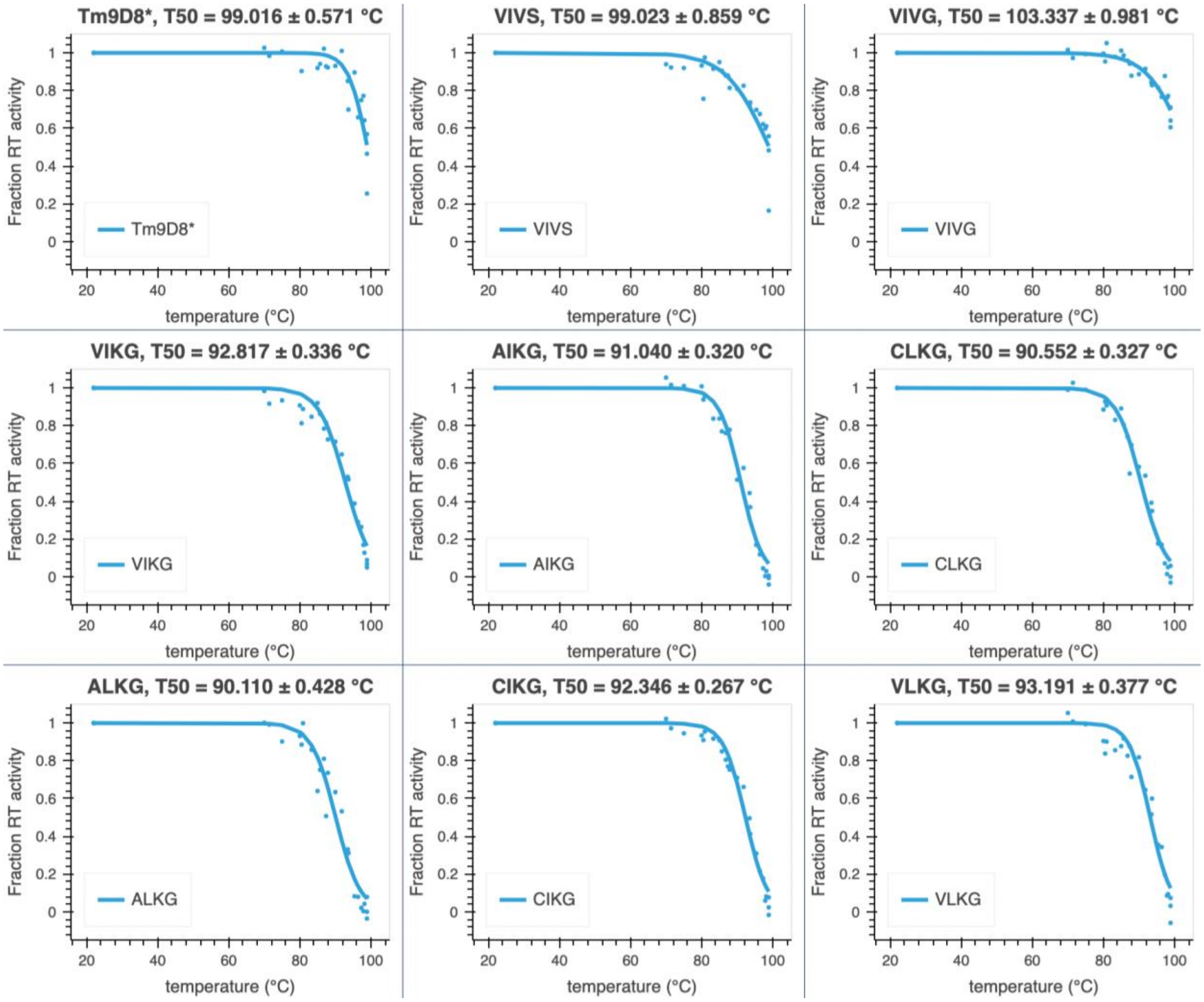

**Figure S36. Sigmoid fits for fraction of room temperature activity vs. 1 h incubation temperature.** Initial rates were divided by the initial rate of the room temperature-incubated sample and plotted versus temperature. These data were fit with a sigmoid that was used to estimate the $T_{50}$ for each variant. No measurements were taken over an incubation temp of 99 °C.

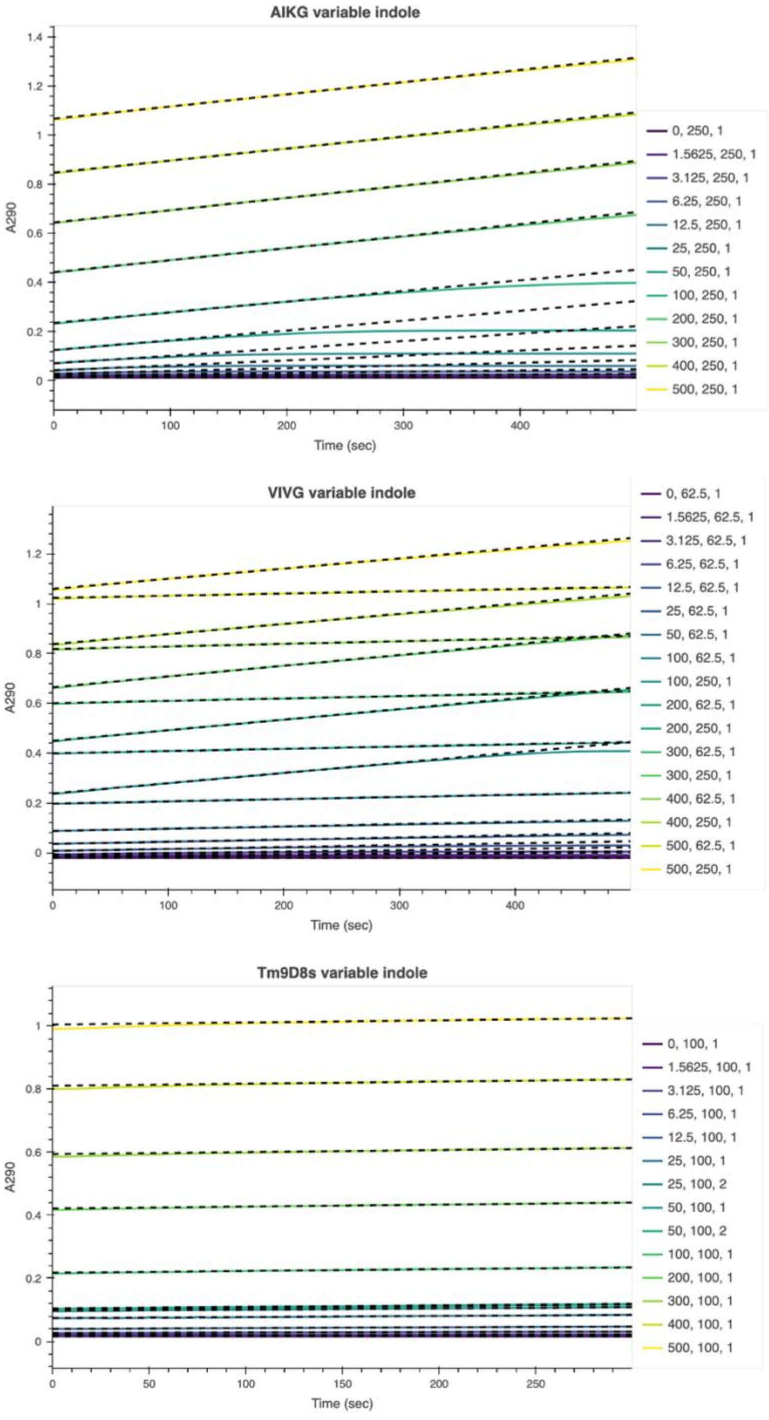

**Figure S37. Fits for variable indole Michaelis-Menten rate estimation data.** Initial rates were estimated from linear or exponential rates. More details provided in the associated code where data processing notebooks can be found.

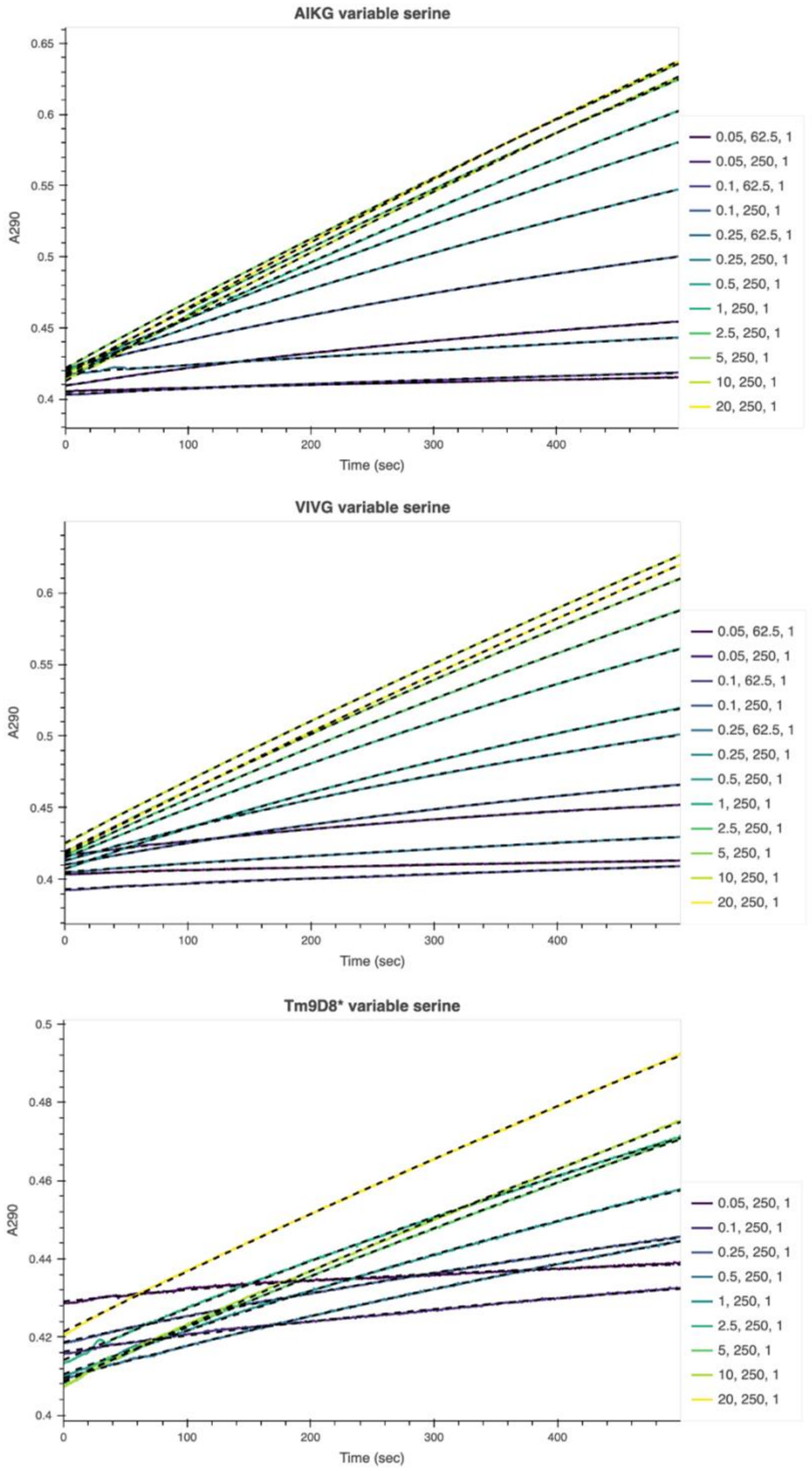

**Figure S38. Fits for variable serine Michaelis-Menten rate estimation data.** Initial rates were estimated from linear or exponential rates. More details provided in the associated code where data processing notebooks can be found.

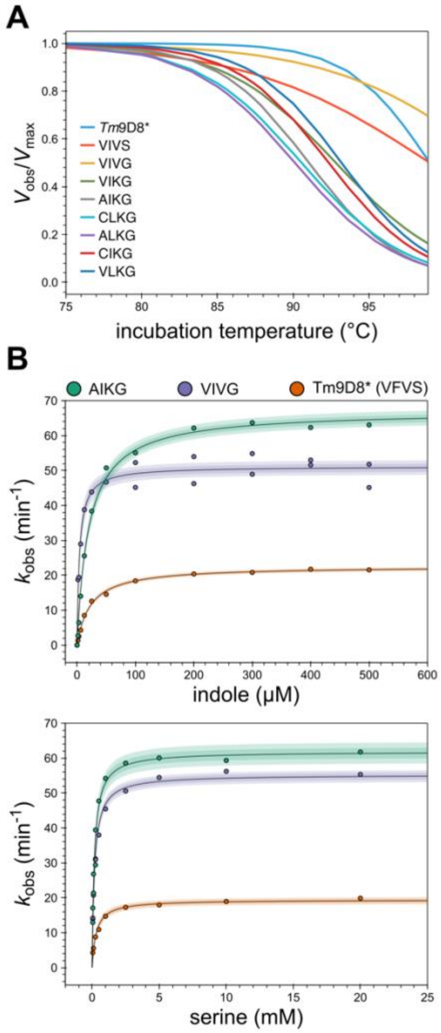

**Figure S39. Biochemical investigation of the top variants.** Stability and activity measurements for select variants. **A** $T_{50}$ curves for all variants in the single possible path from *Tm*9D8* to AIKG as well as the top five variants: AIKG, CLKG, ALKG, CIKG, and VLKG (in order of fitness). **B** Michaelis-Menten curves for *Tm*9D8*, VIVG, and AIKG with different indole concentrations at 20 mM Ser (left) and different Ser concentrations at 200 μM indole (right).

#### IV. Tables

##### Table S1. Primers for knockout strain building and verification

| Name | Sequence (5' → 3') |
|---|---|
| NEB5α_TrpAB::CamR_fwd | TGCCGCCAGCGGAACTGGCGGCTGTGGGATTAACTGCGCGTCGCCGCTTTGTGTAGGCTGGAGCTGCTTC |
| NEB5α_TrpAB::CamR_rev | TTGGCCTCGGTTTTCCAGACGCTGCGCGCATATTAAGGAAAGGAACAATGATGGGAATTAGCCATGGTCC |
| NEB5α_TrpAB_external_fwd | TGCCGCCAGCGGAACTGGC |
| NEB5α_TrpAB_external_rev | TCAAAGACGCACGTCTTTTGGCCTCGG |
| BW25113_TrpAB::KanR_fwd | TGCCGCCAGCGGAACTGGCGGCTGTGGGATTAACTGCGCGTCGCCGCTTTTGTAGGCTGGAGCTGCTTCG |
| BW25113_TrpAB::KanR_rev | TTGGCCTCGGTTTTCCAGACGCTGCGCGCATATTAAGGAAAGGAACAATGATTCCGGGGATCCGTCGACC |
| BW25113_TrpAB_external_fwd | GGTAAGCGAAACGGTAAAAAGATAAATATTAAATGAATTTAGG |
| BW25113_TrpAB_external_rev | GCGCCGGACTTGATTTTAATTCTGC |
| BW25113_TrpAB_internal_fwd | GCCCAGTCATAGCCGAATAGCC |
| BW25113_TrpAB_internal_rev | GGCTATTCGGCTATGACTGGGC |

**Table S2. Primers for *Tm*9D8*-pBAD24 plasmid construction**

| Name | Sequence (5’ → 3’) |
|---|---|
| TrpB_pBAD24_insert_fwd | AGCAGGAGGAATTCGCCAATGAAAGGCTACTTCGGTCCGTACGG |
| TrpB_pBAD24_insert_rev | CCAAGCTTCCCGGGTCATCAGTGGTGGTGGTGGTGGTGC |
| TrpB_pBAD24_bb_fwd | TGACCCGGGAAGCTTGGCTGTTTTGGCGGATGAGAGAAGATTTTCAGC |
| TrpB_pBAD24_bb_rev | TGGCGAATTCCTCCTGCTAGCCCAAAAAAACGGGTATGGAGAAACAG |
| AmpR_internal_fwd | CCAACTTACTTCTGACAACGATCGGAGGACCGAAGGAGCTAACCGCTTTTTTGC |
| AmpR_internal_rev | CGATCGTTGTCAGAAGTAAGTTGGCCGCAGTGTTATCACTCATGGTTATGGCAG |

**Table S3. TrpB sequences**

| Name | Sequence |
|---|---|
| *Tm*9D8*<br><br>in pBAD24: 5276 | ATGAAAGGCTACTTCGGTCCGTACGGTGGCCAGTACGTGCCGGAAATCC<br>TGATGGGAGCTCTGGAAGAACTGGAAGCTGCGTACGAAGGAATCATGAA<br>AGATGAGTCTTTCTGGAAAGAATTCAATGACCTGCTGCGCGATTATGCG<br>GGTCGTCCGACTCCGCTGTACTTCGCACGTCGTCTGTCCGAAAAATACG<br>GTGCTCGCGTATATCTGAAACGTGAAGACCTGCTGCATACTGGTGCGCA<br>TAAAATCAATAACGCTATCGGCCAGGTTCTGCTGGCAAAACTAATGGGC<br>AAAACCCGTATCATTGCTGAAACGGGTGCTGGTCAGCACGGCGTAGCAA<br>CTGCTACCGCAGCAGCGCTGTTCGGTATGGAATGTGTAATCTATATGGG<br>CGAAGAAGACACGATCCGCCAGAAACTAAACGTTGAACGTATGAAACTG<br>CTGGGTGCTAAAGTTGTACCGGTAAAATCCGGTAGCCGTACCCTGAAAG<br>ACGCAATTGACGAAGCTCTGCGTGACTGGATTACCAACCTGCAGACCAC<br>CTATTACGTGTTCGGCTCTGTGGTTGGTCCGCATCCATATCCGATTATC<br>GTACGTAACTTCCAAAAGGTTATCGGCGAAGAGACCAAAAAACAGATTC<br>CAGAAAAAGAAGGCCGTCTGCCGGACTACATCGTTGCGTGCGTGAGCGG<br>TGGTTCTAACGCTGCCGGTATCTTCTATCCGTTTATCGATTCTGGTGTG<br>AAGCTGATCGGCGTAGAAGCCGGTGGCGAAGGTCTGGAAACCGGTAAAC<br>ATGCGGCTTCTCTGCTGAAAGGTAAAATCGGCTACCTGCACGGTTCTAA<br>GACGTTCGTTCTGCAGGATGACTGGGGTCAAGTTCAGGTGAGCCACTCC<br>GTCTCCGCTGGCCTGGACTACTCCGGTGTCGGTCCGGAACACGCCTATT<br>GGCGTGAGACCGGTAAAGTGCTGTACGATGCTGTGACCGATGAAGAAGC<br>TCTGGACGCATTCATCGAACTGTCTCGCCTGGAAGGCATCATCCCAGCC<br>CTGGAGTCTTCTCACGCACTGGCTTATCTGAAGAAGATCAACATCAAGG<br>GTAAAGTTGTGGTGGTTAATCTGTCTGGTCGTGGTGACAAGGATCTGGA<br>ATCTGTACTGAACCACCCGTATGTTCGCGAACGCATCCGCCTCGAGCAC<br>CACCACCACCACCACTGA |
| VIVS<br><br>in pET22b(+): 5277 | ATGAAAGGCTACTTCGGTCCGTACGGTGGCCAGTACGTGCCGGAAATCC<br>TGATGGGAGCTCTGGAAGAACTGGAAGCTGCGTACGAAGGAATCATGAA<br>AGATGAGTCTTTCTGGAAAGAATTCAATGACCTGCTGCGCGATTATGCG<br>GGTCGTCCGACTCCGCTGTACTTCGCACGTCGTCTGTCCGAAAAATACG<br>GTGCTCGCGTATATCTGAAACGTGAAGACCTGCTGCATACTGGTGCGCA<br>TAAAATCAATAACGCTATCGGCCAGGTTCTGCTGGCAAAACTAATGGGC<br>AAAACCCGTATCATTGCTGAAACGGGTGCTGGTCAGCACGGCGTAGCAA<br>CTGCTACCGCAGCAGCGCTGTTCGGTATGGAATGTGTAATCTATATGGG<br>CGAAGAAGACACGATCCGCCAGAAACTAAACGTTGAACGTATGAAACTG<br>CTGGGTGCTAAAGTTGTACCGGTAAAATCCGGTAGCCGTACCCTGAAAG<br>ACGCAATTGACGAAGCTCTGCGTGACTGGATTACCAACCTGCAGACCAC<br>CTATTACGTGATTGGCTCTGTGGTTGGTCCGCATCCATATCCGATTATC<br>GTACGTAACTTCCAAAAGGTTATCGGCGAAGAGACCAAAAAACAGATTC<br>CAGAAAAAGAAGGCCGTCTGCCGGACTACATCGTTGCGTGCGTGAGCGG<br>TGGTTCTAACGCTGCCGGTATCTTCTATCCGTTTATCGATTCTGGTGTG<br>AAGCTGATCGGCGTAGAAGCCGGTGGCGAAGGTCTGGAAACCGGTAAAC<br>ATGCGGCTTCTCTGCTGAAAGGTAAAATCGGCTACCTGCACGGTTCTAA<br>GACGTTCGTTCTGCAGGATGACTGGGGTCAAGTTCAGGTGAGCCACTCC<br>GTCTCCGCTGGCCTGGACTACTCCGGTGTCGGTCCGGAACACGCCTATT<br>GGCGTGAGACCGGTAAAGTGCTGTACGATGCTGTGACCGATGAAGAAGC<br>TCTGGACGCATTCATCGAACTGTCTCGCCTGGAAGGCATCATCCCAGCC<br>CTGGAGTCTTCTCACGCACTGGCTTATCTGAAGAAGATCAACATCAAGG<br>GTAAAGTTGTGGTGGTTAATCTGTCTGGTCGTGGTGACAAGGATCTGGA<br>ATCTGTACTGAACCACCCGTATGTTCGCGAACGCATCCGCCTCGAGCAC<br>CACCACCACCACCACTGA |
| VIVG | ATGAAAGGCTACTTCGGTCCGTACGGTGGCCAGTACGTGCCGGAAATCC<br>TGATGGGAGCTCTGGAAGAACTGGAAGCTGCGTACGAAGGAATCATGAA<br>AGATGAGTCTTTCTGGAAAGAATTCAATGACCTGCTGCGCGATTATGCG |

| | |
|---|---|
| in pET22b(+): 5278 | GGTCGTCCGACTCCGCTGTACTTCGCACGTCGTCTGTCCGAAAAATACG<br>GTGCTCGCGTATATCTGAAACGTGAAGACCTGCTGCATACTGGTGCGCA<br>TAAAATCAATAACGCTATCGGCCAGGTTCTGCTGGCAAAACTAATGGGC<br>AAAACCCGTATCATTGCTGAAACGGGTGCTGGTCAGCACGGCGTAGCAA<br>CTGCTACCGCAGCAGCGCTGTTCGGTATGGAATGTGTAATCTATATGGG<br>CGAAGAAGACACGATCCGCCAGAAACTAAACGTTGAACGTATGAAACTG<br>CTGGGTGCTAAAGTTGTACCGGTAAAATCCGGTAGCCGTACCCTGAAAG<br>ACGCAATTGACGAAGCTCTGCGTGACTGGATTACCAACCTGCAGACCAC<br>CTATTACGTGATTGGCTCTGTGGTTGGTCCGCATCCATATCCGATTATC<br>GTACGTAACTTCCAAAAGGTTATCGGCGAAGAGACCAAAAAACAGATTC<br>CAGAAAAAGAAGGCCGTCTGCCGGACTACATCGTTGCGTGCGTGGGTGG<br>TGGTTCTAACGCTGCCGGTATCTTCTATCCGTTTATCGATTCTGGTGTG<br>AAGCTGATCGGCGTAGAAGCCGGTGGCGAAGGTCTGGAAACCGGTAAAC<br>ATGCGGCTTCTCTGCTGAAAGGTAAAATCGGCTACCTGCACGGTTCTAA<br>GACGTTCGTTCTGCAGGATGACTGGGGTCAAGTTCAGGTGAGCCACTCC<br>GTCTCCGCTGGCCTGGACTACTCCGGTGTCGGTCCGGAACACGCCTATT<br>GGCGTGAGACCGGTAAAGTGCTGTACGATGCTGTGACCGATGAAGAAGC<br>TCTGGACGCATTCATCGAACTGTCTCGCCTGGAAGGCATCATCCCAGCC<br>CTGGAGTCTTCTCACGCACTGGCTTATCTGAAGAAGATCAACATCAAGG<br>GTAAAGTTGTGGTGGTTAATCTGTCTGGTCGTGGTGACAAGGATCTGGA<br>ATCTGTACTGAACCACCCGTATGTTCGCGAACGCATCCGCCTCGAGCAC<br>CACCACCACCACCACTGA |
| VIKG<br><br>in pET22b(+): 5279 | ATGAAAGGCTACTTCGGTCCGTACGGTGGCCAGTACGTGCCGGAAATCC<br>TGATGGGAGCTCTGGAAGAACTGGAAGCTGCGTACGAAGGAATCATGAA<br>AGATGAGTCTTTCTGGAAAGAATTCAATGACCTGCTGCGCGATTATGCG<br>GGTCGTCCGACTCCGCTGTACTTCGCACGTCGTCTGTCCGAAAAATACG<br>GTGCTCGCGTATATCTGAAACGTGAAGACCTGCTGCATACTGGTGCGCA<br>TAAAATCAATAACGCTATCGGCCAGGTTCTGCTGGCAAAACTAATGGGC<br>AAAACCCGTATCATTGCTGAAACGGGTGCTGGTCAGCACGGCGTAGCAA<br>CTGCTACCGCAGCAGCGCTGTTCGGTATGGAATGTGTAATCTATATGGG<br>CGAAGAAGACACGATCCGCCAGAAACTAAACGTTGAACGTATGAAACTG<br>CTGGGTGCTAAAGTTGTACCGGTAAAATCCGGTAGCCGTACCCTGAAAG<br>ACGCAATTGACGAAGCTCTGCGTGACTGGATTACCAACCTGCAGACCAC<br>CTATTACGTGATTGGCTCTGTGGTTGGTCCGCATCCATATCCGATTATC<br>GTACGTAACTTCCAAAAGGTTATCGGCGAAGAGACCAAAAAACAGATTC<br>CAGAAAAAGAAGGCCGTCTGCCGGACTACATCGTTGCGTGCGTGAAGGG<br>TGGTGGTTCTAACGCTGCCGGTATCTTCTATCCGTTTATCGATTCTGGT<br>GTGAAGCTGATCGGCGTAGAAGCCGGTGGCGAAGGTCTGGAAACCGGTA<br>AACATGCGGCTTCTCTGCTGAAAGGTAAAATCGGCTACCTGCACGGTTC<br>TAAGACGTTCGTTCTGCAGGATGACTGGGGTCAAGTTCAGGTGAGCCAC<br>TCCGTCTCCGCTGGCCTGGACTACTCCGGTGTCGGTCCGGAACACGCCT<br>ATTGGCGTGAGACCGGTAAAGTGCTGTACGATGCTGTGACCGATGAAGA<br>AGCTCTGGACGCATTCATCGAACTGTCTCGCCTGGAAGGCATCATCCCA<br>GCCCTGGAGTCTTCTCACGCACTGGCTTATCTGAAGAAGATCAACATCA<br>AGGGTAAAGTTGTGGTGGTTAATCTGTCTGGTCGTGGTGACAAGGATCT<br>GGAATCTGTACTGAACCACCCGTATGTTCGCGAACGCATCCGCCTCGAG<br>CACCACCACCACCACCACTGA |
| AIKG<br><br>in pET22b(+): 5280 | ATGAAAGGCTACTTCGGTCCGTACGGTGGCCAGTACGTGCCGGAAATCC<br>TGATGGGAGCTCTGGAAGAACTGGAAGCTGCGTACGAAGGAATCATGAA<br>AGATGAGTCTTTCTGGAAAGAATTCAATGACCTGCTGCGCGATTATGCG<br>GGTCGTCCGACTCCGCTGTACTTCGCACGTCGTCTGTCCGAAAAATACG<br>GTGCTCGCGTATATCTGAAACGTGAAGACCTGCTGCATACTGGTGCGCA<br>TAAAATCAATAACGCTATCGGCCAGGTTCTGCTGGCAAAACTAATGGGC<br>AAAACCCGTATCATTGCTGAAACGGGTGCTGGTCAGCACGGCGTAGCAA<br>CTGCTACCGCAGCAGCGCTGTTCGGTATGGAATGTGTAATCTATATGGG<br>CGAAGAAGACACGATCCGCCAGAAACTAAACGTTGAACGTATGAAACTG |

| | |
|---|---|
| | CTGGGTGCTAAAGTTGTACCGGTAAAATCCGGTAGCCGTACCCTGAAAG<br>ACGCAATTGACGAAGCTCTGCGTGACTGGATTACCAACCTGCAGACCAC<br>CTATTACGCGATTGGCTCTGTGGTTGGTCCGCATCCATATCCGATTATC<br>GTACGTAACTTCCAAAAGGTTATCGGCGAAGAGACCAAAAAACAGATTC<br>CAGAAAAAGAAGGCCGTCTGCCGGACTACATCGTTGCGTGCAAGGGTGG<br>TGGTTCTAACGCTGCCGGTATCTTCTATCCGTTTATCGATTCTGGTGTG<br>AAGCTGATCGGCGTAGAAGCCGGTGGCGAAGGTCTGGAAACCGGTAAAC<br>ATGCGGCTTCTCTGCTGAAAGGTAAAATCGGCTACCTGCACGGTTCTAA<br>GACGTTCGTTCTGCAGGATGACTGGGGTCAAGTTCAGGTGAGCCACTCC<br>GTCTCCGCTGGCCTGGACTACTCCGGTGTCGGTCCGGAACACGCCTATT<br>GGCGTGAGACCGGTAAAGTGCTGTACGATGCTGTGACCGATGAAGAAGC<br>TCTGGACGCATTCATCGAACTGTCTCGCCTGGAAGGCATCATCCCAGCC<br>CTGGAGTCTTCTCACGCACTGGCTTATCTGAAGAAGATCAACATCAAGG<br>GTAAAGTTGTGGTGGTTAATCTGTCTGGTCGTGGTGACAAGGATCTGGA<br>ATCTGTACTGAACCACCCGTATGTTCGCGAACGCATCCGCCTCGAGCAC<br>CACCACCACCACCACTGA |
| CLKG<br><br>in pET22b(+): 5281 | ATGAAAGGCTACTTCGGTCCGTACGGTGGCCAGTACGTGCCGGAAATCC<br>TGATGGGAGCTCTGGAAGAACTGGAAGCTGCGTACGAAGGAATCATGAA<br>AGATGAGTCTTTCTGGAAAGAATTCAATGACCTGCTGCGCGATTATGCG<br>GGTCGTCCGACTCCGCTGTACTTCGCACGTCGTCTGTCCGAAAAATACG<br>GTGCTCGCGTATATCTGAAACGTGAAGACCTGCTGCATACTGGTGCGCA<br>TAAAATCAATAACGCTATCGGCCAGGTTCTGCTGGCAAAACTAATGGGC<br>AAAACCCGTATCATTGCTGAAACGGGTGCTGGTCAGCACGGCGTAGCAA<br>CTGCTACCGCAGCAGCGCTGTTCGGTATGGAATGTGTAATCTATATGGG<br>CGAAGAAGACACGATCCGCCAGAAACTAAACGTTGAACGTATGAAACTG<br>CTGGGTGCTAAAGTTGTACCGGTAAAATCCGGTAGCCGTACCCTGAAAG<br>ACGCAATTGACGAAGCTCTGCGTGACTGGATTACCAACCTGCAGACCAC<br>CTATTACTGTCTGGGCTCTGTGGTTGGTCCGCATCCATATCCGATTATC<br>GTACGTAACTTCCAAAAGGTTATCGGCGAAGAGACCAAAAAACAGATTC<br>CAGAAAAAGAAGGCCGTCTGCCGGACTACATCGTTGCGTGCAAGGGTGG<br>TGGTTCTAACGCTGCCGGTATCTTCTATCCGTTTATCGATTCTGGTGTG<br>AAGCTGATCGGCGTAGAAGCCGGTGGCGAAGGTCTGGAAACCGGTAAAC<br>ATGCGGCTTCTCTGCTGAAAGGTAAAATCGGCTACCTGCACGGTTCTAA<br>GACGTTCGTTCTGCAGGATGACTGGGGTCAAGTTCAGGTGAGCCACTCC<br>GTCTCCGCTGGCCTGGACTACTCCGGTGTCGGTCCGGAACACGCCTATT<br>GGCGTGAGACCGGTAAAGTGCTGTACGATGCTGTGACCGATGAAGAAGC<br>TCTGGACGCATTCATCGAACTGTCTCGCCTGGAAGGCATCATCCCAGCC<br>CTGGAGTCTTCTCACGCACTGGCTTATCTGAAGAAGATCAACATCAAGG<br>GTAAAGTTGTGGTGGTTAATCTGTCTGGTCGTGGTGACAAGGATCTGGA<br>ATCTGTACTGAACCACCCGTATGTTCGCGAACGCATCCGCCTCGAGCAC<br>CACCACCACCACCACTGA |
| ALKG<br><br>in pET22b(+): 5282 | ATGAAAGGCTACTTCGGTCCGTACGGTGGCCAGTACGTGCCGGAAATCC<br>TGATGGGAGCTCTGGAAGAACTGGAAGCTGCGTACGAAGGAATCATGAA<br>AGATGAGTCTTTCTGGAAAGAATTCAATGACCTGCTGCGCGATTATGCG<br>GGTCGTCCGACTCCGCTGTACTTCGCACGTCGTCTGTCCGAAAAATACG<br>GTGCTCGCGTATATCTGAAACGTGAAGACCTGCTGCATACTGGTGCGCA<br>TAAAATCAATAACGCTATCGGCCAGGTTCTGCTGGCAAAACTAATGGGC<br>AAAACCCGTATCATTGCTGAAACGGGTGCTGGTCAGCACGGCGTAGCAA<br>CTGCTACCGCAGCAGCGCTGTTCGGTATGGAATGTGTAATCTATATGGG<br>CGAAGAAGACACGATCCGCCAGAAACTAAACGTTGAACGTATGAAACTG<br>CTGGGTGCTAAAGTTGTACCGGTAAAATCCGGTAGCCGTACCCTGAAAG<br>ACGCAATTGACGAAGCTCTGCGTGACTGGATTACCAACCTGCAGACCAC<br>CTATTACTGTATTGGCTCTGTGGTTGGTCCGCATCCATATCCGATTATC<br>GTACGTAACTTCCAAAAGGTTATCGGCGAAGAGACCAAAAAACAGATTC<br>CAGAAAAAGAAGGCCGTCTGCCGGACTACATCGTTGCGTGCAAGGGTGG<br>TGGTTCTAACGCTGCCGGTATCTTCTATCCGTTTATCGATTCTGGTGTG |

| | |
|---|---|
| | AAGCTGATCGGCGTAGAAGCCGGTGGCGAAGGTCTGGAAACCGGTAAAC<br>ATGCGGCTTCTCTGCTGAAAGGTAAAATCGGCTACCTGCACGGTTCTAA<br>GACGTTCGTTCTGCAGGATGACTGGGGTCAAGTTCAGGTGAGCCACTCC<br>GTCTCCGCTGGCCTGGACTACTCCGGTGTCGGTCCGGAACACGCCTATT<br>GGCGTGAGACCGGTAAAGTGCTGTACGATGCTGTGACCGATGAAGAAGC<br>TCTGGACGCATTCATCGAACTGTCTCGCCTGGAAGGCATCATCCCAGCC<br>CTGGAGTCTTCTCACGCACTGGCTTATCTGAAGAAGATCAACATCAAGG<br>GTAAAGTTGTGGTGGTTAATCTGTCTGGTCGTGGTGACAAGGATCTGGA<br>ATCTGTACTGAACCACCCGTATGTTCGCGAACGCATCCGCCTCGAGCAC<br>CACCACCACCACCACTGA |
| CIKG<br><br>in pET22b(+): 5283 | ATGAAAGGCTACTTCGGTCCGTACGGTGGCCAGTACGTGCCGGAAATCC<br>TGATGGGAGCTCTGGAAGAACTGGAAGCTGCGTACGAAGGAATCATGAA<br>AGATGAGTCTTTCTGGAAAGAATTCAATGACCTGCTGCGCGATTATGCG<br>GGTCGTCCGACTCCGCTGTACTTCGCACGTCGTCTGTCCGAAAAATACG<br>GTGCTCGCGTATATCTGAAACGTGAAGACCTGCTGCATACTGGTGCGCA<br>TAAAATCAATAACGCTATCGGCCAGGTTCTGCTGGCAAAACTAATGGGC<br>AAAACCCGTATCATTGCTGAAACGGGTGCTGGTCAGCACGGCGTAGCAA<br>CTGCTACCGCAGCAGCGCTGTTCGGTATGGAATGTGTAATCTATATGGG<br>CGAAGAAGACACGATCCGCCAGAAACTAAACGTTGAACGTATGAAACTG<br>CTGGGTGCTAAAGTTGTACCGGTAAAATCCGGTAGCCGTACCCTGAAAG<br>ACGCAATTGACGAAGCTCTGCGTGACTGGATTACCAACCTGCAGACCAC<br>CTATTACGCGCTGGGCTCTGTGGTTGGTCCGCATCCATATCCGATTATC<br>GTACGTAACTTCCAAAAGGTTATCGGCGAAGAGACCAAAAAACAGATTC<br>CAGAAAAAGAAGGCCGTCTGCCGGACTACATCGTTGCGTGCAAGGGTGG<br>TGGTTCTAACGCTGCCGGTATCTTCTATCCGTTTATCGATTCTGGTGTG<br>AAGCTGATCGGCGTAGAAGCCGGTGGCGAAGGTCTGGAAACCGGTAAAC<br>ATGCGGCTTCTCTGCTGAAAGGTAAAATCGGCTACCTGCACGGTTCTAA<br>GACGTTCGTTCTGCAGGATGACTGGGGTCAAGTTCAGGTGAGCCACTCC<br>GTCTCCGCTGGCCTGGACTACTCCGGTGTCGGTCCGGAACACGCCTATT<br>GGCGTGAGACCGGTAAAGTGCTGTACGATGCTGTGACCGATGAAGAAGC<br>TCTGGACGCATTCATCGAACTGTCTCGCCTGGAAGGCATCATCCCAGCC<br>CTGGAGTCTTCTCACGCACTGGCTTATCTGAAGAAGATCAACATCAAGG<br>GTAAAGTTGTGGTGGTTAATCTGTCTGGTCGTGGTGACAAGGATCTGGA<br>ATCTGTACTGAACCACCCGTATGTTCGCGAACGCATCCGCCTCGAGCAC<br>CACCACCACCACCACTGA |
| VLKG<br><br>in pET22b(+): 5284 | ATGAAAGGCTACTTCGGTCCGTACGGTGGCCAGTACGTGCCGGAAATCC<br>TGATGGGAGCTCTGGAAGAACTGGAAGCTGCGTACGAAGGAATCATGAA<br>AGATGAGTCTTTCTGGAAAGAATTCAATGACCTGCTGCGCGATTATGCG<br>GGTCGTCCGACTCCGCTGTACTTCGCACGTCGTCTGTCCGAAAAATACG<br>GTGCTCGCGTATATCTGAAACGTGAAGACCTGCTGCATACTGGTGCGCA<br>TAAAATCAATAACGCTATCGGCCAGGTTCTGCTGGCAAAACTAATGGGC<br>AAAACCCGTATCATTGCTGAAACGGGTGCTGGTCAGCACGGCGTAGCAA<br>CTGCTACCGCAGCAGCGCTGTTCGGTATGGAATGTGTAATCTATATGGG<br>CGAAGAAGACACGATCCGCCAGAAACTAAACGTTGAACGTATGAAACTG<br>CTGGGTGCTAAAGTTGTACCGGTAAAATCCGGTAGCCGTACCCTGAAAG<br>ACGCAATTGACGAAGCTCTGCGTGACTGGATTACCAACCTGCAGACCAC<br>CTATTACGTGCTGGGCTCTGTGGTTGGTCCGCATCCATATCCGATTATC<br>GTACGTAACTTCCAAAAGGTTATCGGCGAAGAGACCAAAAAACAGATTC<br>CAGAAAAAGAAGGCCGTCTGCCGGACTACATCGTTGCGTGCAAGGGTGG<br>TGGTTCTAACGCTGCCGGTATCTTCTATCCGTTTATCGATTCTGGTGTG<br>AAGCTGATCGGCGTAGAAGCCGGTGGCGAAGGTCTGGAAACCGGTAAAC<br>ATGCGGCTTCTCTGCTGAAAGGTAAAATCGGCTACCTGCACGGTTCTAA<br>GACGTTCGTTCTGCAGGATGACTGGGGTCAAGTTCAGGTGAGCCACTCC<br>GTCTCCGCTGGCCTGGACTACTCCGGTGTCGGTCCGGAACACGCCTATT<br>GGCGTGAGACCGGTAAAGTGCTGTACGATGCTGTGACCGATGAAGAAGC<br>TCTGGACGCATTCATCGAACTGTCTCGCCTGGAAGGCATCATCCCAGCC |

| | |
|---|---|
| | CTGGAGTCTTCTCACGCACTGGCTTATCTGAAGAAGATCAACATCAAGG<br>GTAAAGTTGTGGTGGTTAATCTGTCTGGTCGTGGTGACAAGGATCTGGA<br>ATCTGTACTGAACCACCCGTATGTTCGCGAACGCATCCGCCTCGAGCAC<br>CACCACCACCACCACTGA |

**Table S4. Primers for single- and double-site saturation mutagenesis for preliminary assays.**

| Name | Sequence (5' → 3') |
| --- | --- |
| Tm9D8*_184X_fwd | CTGCAGACCACCTATTACGTGXXXGGCTCTGTGGTTGGTCC |
| Tm9D8*_118X_fwd | GGCGTAGCAACTGCTACCXXXGCAGCGCTGTTCGGTATGGAATGTGTAATCTATATGG |
| Tm9D8*_184_rev | CACGTAATAGGTGGTCTGCAGGTTGGTAATCCAGTCACGCAGAGCT |
| Tm9D8*_118_rev | GGTAGCAGTTGCTACGCCGTGCTGACCAGC |
| Tm9D8*_301X_fwd | TCCGCTGGCCTGGACXXXTCCGGTGTCGGTCCGGA |
| Tm9D8*_301_rev | GTCCAGGCCAGCGGAGACGGAGTGGCTCACCTGAACT |
| AmpR_internal_fwd | CCAACTTACTTCTGACAACGATCGGAGGACCGAAGGAGCTAACCGCTTTTTTGC |
| AmpR_internal_rev | CGATCGTTGTCAGAAGTAAGTTGGCCGCAGTGTTATCACTCATGGTTATGGCAG |

XXX = mix of three primers with NDT, VHG, or TGG at that position mixed in a ratio of 12:9:1. This is based on methods presented by Kille et al. (6).

**Table S5. Primers for construction of triple-site saturation libraries**

| Library | Primer | Primer name | Sequence (5' → 3') |
|---|---|---|---|
| **A**<br><br>104<br>105<br>106 | F Gap | 9D8s_106_gap_F | GGTGCTGGTCAGCACG |
| | F Library | 9D8s_104-105-106_F | AATGGGCAAAACCCGTATCATTNNKNNKNNKGGTGCTGGTCAGCACG |
| | R | 9D8s_104_R | AATGATACGGGTTTTGCCCATTAGTTTTGCCAGCAGAACCTGGC |
| **B**<br><br>105<br>106<br>107 | F Gap | 9D8s_107_gap_F | GCTGGTCAGCACGGC |
| | F Library | 9D8s_105-106-107_F | AATGGGCAAAACCCGTATCATTGCTNNKNNKNNKGCTGGTCAGCACGGC |
| | R | 9D8s_104_R | AATGATACGGGTTTTGCCCATTAGTTTTGCCAGCAGAACCTGGC |
| **C**<br><br>106<br>107<br>108 | F Gap | 9D8s_108_gap_F | GGTCAGCACGGCGTAG |
| | F Library | 9D8s_106-107-108_F | AATGGGCAAAACCCGTATCATTGCTGAANNKNNKNNKGGTCAGCACGGCGTAG |
| | R | 9D8s_104_R | AATGATACGGGTTTTGCCCATTAGTTTTGCCAGCAGAACCTGGC |
| **D**<br><br>117<br>118<br>119 | F Gap | 9D8s_119_gap_F | GCGCTGTTCGGTATGGAAT |
| | F Library | 9D8s_117-118-119_F | ACGGCGTAGCAACTGCTNNKNNKNNKGCGCTGTTCGGTATGGAATGTGTAATC |
| | R | 9D8s_117_R | AGCAGTTGCTACGCCGTGCTGACCAGCACCCGTTTCAG |
| **E**<br><br>184<br>185<br>186 | F Gap | 9D8s_186_gap_F | GTGGTTGGTCCGCATCC |
| | F Library | 9D8s_184-185-186_F | CTGCAGACCACCTATTACGTGNNKNNKNNKGTGGTTGGTCCGCATCCATATCC |
| | R | 9D8s_184_R | CACGTAATAGGTGGTCTGCAGGTTGGTAATCCAGTCACGCAGAGCT |
| **F***<br><br>162<br>166<br>301 | F Gap | 9D8s_166_gap_F | GACGAAGCTCTGCGTGAC |
| | F Library | 9D8s_162-166_F | GTAAAATCCGGTAGCCGTACCNNKAAAGACGCANNKGACGAAGCTCTG |
| | R | 9D8s_162_R | GGTACGGCTACCGGATTTTACCGGTACAACTTTAGCACCCAGCAG |
| **G***<br><br>227<br>228<br>301 | F Gap | 9D8s_228_gap_F | GGTGGTTCTAACGCTGCC |
| | F Library | 9D8s_227-228_F | GGACTACATCGTTGCGTGCNNKNNKGGTGGTTCTAACGCTGCCGGTA |
| | R | 9D8s_227_R | GCACGCAACGATGTAGTCCGGCAGACGGCCTTCTTTTTCTGG |

| | | | |
|---|---|---|---|
| **H**<br><br>228<br>230<br>231 | F Gap | 9D8s_231_gap_F | AACGCTGCCGGTATCTTCTAT |
| | F Library | 9D8s_228-230-231_F | GGACTACATCGTTGCGTGCGTGNNKGGTN NKNNKAACGCTGCCGGTATCTTCTATCCG |
| | R | 9D8s_227_R | GCACGCAACGATGTAGTCCGGCAGACGGC CTTCTTTTTCTGG |
| **I**<br><br>182<br>183<br>184 | F Gap | 9D8s_184_gap_F | GGCTCTGTGGTTGGTCC |
| | F Library | 9D8s_182-183-184_F | CCAACCTGCAGACCACCTATNNKNNKNNK GGCTCTGTGGTTGGTCCGC |
| | R | 9D8s_182_R | ATAGGTGGTCTGCAGGTTGGTAATCCAGT CACGCAGAGCTTCGT |

*These libraries used a template of a 301X plasmid library created with primers in Table S4.

**Table S6. Primers for construction of the quadruple-site saturation library**

| Library | Primer | Primer name | Sequence (5’ → 3’) |
|---|---|---|---|
| 4-site<br><br>183<br>184<br>227<br>228 | F | 9D8s_foursite_183-184-227-228_bb_f | GGTGGTTCTAACGCTGCCGGTATCTTCTATCCGTTTATCG |
| | F Gap | 9D8s_foursite_183-184-227-228_inner_f | GGCTCTGTGGTTGGTCCGCATCCATATCCG |
| | F Library | 9D8s_foursite_183-184-227-228_outer_NNK_f | CCAACCTGCAGACCACCTATTACNNKNNKGGCTCTGTGGTTGGTCCGC |
| | R | 9D8s_foursite_183-184-227-228_bb_r | GTAATAGGTGGTCTGCAGGTTGGTAATCCAGTCACGC |
| | R Gap | 9D8s_foursite_183-184-227-228_inner_r | GCACGCAACGATGTAGTCCGGCAGACGGCCTTC |
| | R Library | 9D8s_foursite_183-184-227-228_outer_NNK_r | GGCAGCGTTAGAACCACCMNNMNNGCACGCAACGATGTAGTCCG |

**Table S7. $OD_{600}$ over time by library: libraries A, B, and C**

| Hours | A1 | A2 | B1 | B2 | C1 | C2 |
|---|---|---|---|---|---|---|
| **0** | 0.1* | 0.1 | 0.1* | 0.1 | 0.1* | 0.1 |
| **18** | 0.72* | 0.75 | 0.75* | 0.84 | 0.74* | 0.76 |
| **20** | 0.78 | 0.83 | 0.83 | 0.98 | 0.78 | 0.84 |
| **24** | 0.94 | 1.01 | 1.09 | 1.50 | 0.86 | 0.92 |
| **44** | 2.55* | 2.7 | 3.3* | 3.85 | 1.95* | 4.15 |

*These samples were sequenced. Based on preliminary results where most variants were inactive, the remaining timepoints and replicates were not sequenced.

**Table S8. $OD_{600}$ over time by library: libraries D, E, F, G, H, and I**

| Time (h) | D1 | D2 | E1 | E2 | F1 | F2 | G1 | G2 | H1 | H2 | I1 | I2 |
|---|---|---|---|---|---|---|---|---|---|---|---|---|
| **0** | 0.05 | 0.05 | 0.05 | 0.05 | 0.05 | 0.05 | 0.05 | 0.05 | 0.05 | 0.05 | 0.05 | 0.05 |
| **12** | 0.19[A] | 0.18[A] | 0.20[A] | 0.20[A] | 0.17[A] | 0.17[A] | 0.14[A] | 0.14[A] | 0.15[A] | 0.14[A] | 0.36[A] | 0.39[A] |
| **16** | 0.29 | 0.28 | 0.27[B] | 0.26[B] | 0.20[C] | 0.20[C] | 0.18 | 0.18 | 0.19 | 0.18 | 0.83 | 0.87 |
| **20** | 0.51 | 0.49 | 0.47 | 0.44 | 0.23 | 0.24 | 0.23 | 0.23 | 0.26 | 0.26 | 1.24 | 1.36 |
| **24** | 0.85 | 0.97 | 0.91 | 0.94 | 0.27[C] | 0.27[C] | 0.44 | 0.44 | 0.67 | 0.58 | 1.7 | 2.1 |
| **36** | 1.42 | 1.81 | 1.41 | 1.54 | 0.79 | 0.79 | 2.0 | 1.95 | 2.9 | 1.85 | 1.95 | 2.25 |

[A] All 12 h timepoints were dropped due to (1) low correlation between replicates compared to later timepoints and (2) weaker separation between the stop-codon-containing variants and remaining variants.

[B] The 16 h timepoint for Lib E also showed lower correlation between replicates compared later timepoints.

[C] These timepoints were omitted due to the near absence of correlation between replicates ($r^2 \approx 0$), potentially due to errors arising from sampling, barcoding, amplification, or sequencing.

**Table S9. $OD_{600}$ over time: quadruple-site library**

| Time (h) | 4-site rep #1 | 4-site rep #2 |
|---|---|---|
| **0** | 0.025 | 0.025 |
| **12** | 0.19 | 0.19 |
| **16** | 0.51 | 0.52 |
| **20** | 1.26 | 1.34 |
| **24** | 1.50 | 1.625 |
| **28** | 1.675 | 1.75 |
| **36** | 1.75 | 1.875 |

**Table S10. Sequencing preparation primers for triple- and quadruple-site libraries**

| Primer name | Barcode | Seed (bp) | F/R | Sequence 5’ → 3’ |
|---|---|---|---|---|
| pr191_seq_TGT_266-285_F | TGT | 266-285 | F | TCGTCGGCAGCGTCAGATGTGTATAAGAGACAGNNNNNNNNNNTGTGCCAGGTTCTGCTGGCAAAA |
| pr192_seq_GTG_266-285_F | GTG | 266-285 | F | TCGTCGGCAGCGTCAGATGTGTATAAGAGACAGNNNNNNNNNNGTGGCCAGGTTCTGCTGGCAAAA |
| pr193_seq_ACA_266-285_F | ACA | 266-285 | F | TCGTCGGCAGCGTCAGATGTGTATAAGAGACAGNNNNNNNNNNACAGCCAGGTTCTGCTGGCAAAA |
| pr195_seq_TGT_415-394_R | TGT | 415-394 | R | GTCTCGTGGGCTCGGAGATGTGTATAAGAGACAGNNNNNNNNNNTGTTCTGGCGGATCGTGTCTTCTTC |
| pr196_seq_GTG_415-394_R | GTG | 415-394 | R | GTCTCGTGGGCTCGGAGATGTGTATAAGAGACAGNNNNNNNNNNGTGTCTGGCGGATCGTGTCTTCTTC |
| pr197_seq_ACA_415-394_R | ACA | 415-394 | R | GTCTCGTGGGCTCGGAGATGTGTATAAGAGACAGNNNNNNNNNNACATCTGGCGGATCGTGTCTTCTTC |
| pr206_seq_TGT_448-473_F | TGT | 448-473 | F | TCGTCGGCAGCGTCAGATGTGTATAAGAGACAGNNNNNNNNNNTGTGCTAAAGTTGTACCGGTAAAATCCGG |
| pr207_seq_GTG_448-473_F | GTG | 448-473 | F | TCGTCGGCAGCGTCAGATGTGTATAAGAGACAGNNNNNNNNNNGTGGCTAAAGTTGTACCGGTAAAATCCGG |
| pr209_seq_TGT_589-565_R | TGT | 589-565 | R | GTCTCGTGGGCTCGGAGATGTGTATAAGAGACAGNNNNNNNNNNTGTCGATAATCGGATATGGATGCGGACC |
| pr210_seq_GTG_589-565_R | GTG | 589-565 | R | GTCTCGTGGGCTCGGAGATGTGTATAAGAGACAGNNNNNNNNNNGTGCGATAATCGGATATGGATGCGGACC |
| pr215_seq_TGT_659-678_F | TGT | 659-678 | F | TCGTCGGCAGCGTCAGATGTGTATAAGAGACAGNNNNNNNNNNTGTCGGACTACATCGTTGCGTGC |
| pr216_seq_GTG_659-678_F | GTG | 659-678 | F | TCGTCGGCAGCGTCAGATGTGTATAAGAGACAGNNNNNNNNNNGTGCGGACTACATCGTTGCGTGC |
| pr212_seq_TGT_929-911_R | TGT | 929-911 | R | GTCTCGTGGGCTCGGAGATGTGTATAAGAGACAGNNNNNNNNNNTGTTAGGCGTGTTCCGGACCG |
| pr213_seq_GTG_929-911_R | GTG | 929-911 | R | GTCTCGTGGGCTCGGAGATGTGTATAAGAGACAGNNNNNNNNNNGTGTAGGCGTGTTCCGGACCG |
| pr018_seq_TGT_518-540_F | TGT | 518-540 | F | TCGTCGGCAGCGTCAGATGTGTATAAGAGACAGNNNNNNNNNNTGTGGATTACCAACCTGCAGACCACC |
| pr019_seq_GTG_518-540_F | GTG | 518-540 | F | TCGTCGGCAGCGTCAGATGTGTATAAGAGACAGNNNNNNNNNNGTGGGATTACCAACCTGCAGACCACC |

| | | | | |
|---|---|---|---|---|
| pr020_seq_ACA_518-540_F | ACA | 518-540 | F | TCGTCGGCAGCGTCAGATGTGTATAAGAGACAGNNNNNNNNNNACAGGATTACCAACCTGCAGACCACC |
| pr021_seq_TGT_709-688_R | TGT | 709-688 | R | GTCTCGTGGGCTCGGAGATGTGTATAAGAGACAGNNNNNNNNNNTGTAGATACCGGCAGCGTTAGAACC |
| pr022_seq_GTG_709-688_R | GTG | 709-688 | R | GTCTCGTGGGCTCGGAGATGTGTATAAGAGACAGNNNNNNNNNNGTGAGATACCGGCAGCGTTAGAACC |
| pr023_seq_ACA_709-688_R | ACA | 709-688 | R | GTCTCGTGGGCTCGGAGATGTGTATAAGAGACAGNNNNNNNNNNACAAGATACCGGCAGCGTTAGAACC |

**Table S11. Mapping sequencing primers to libraries**

| Primer name | Libraries |
|---|---|
| pr191_seq_TGT_266-285_F | A/T0<br>B/T0<br>C/T0<br>D/T0, D1/T1–T5 |
| pr192_seq_GTG_266-285_F | A1/T1<br>B1/T1<br>C1/T1<br>D2/T1–T5 |
| pr193_seq_ACA_266-285_F | A1/T4<br>B1/T4<br>C1/T4 |
| pr195_seq_TGT_415-394_R | A/T0<br>B/T0<br>C/T0<br>D1/T1–T5 |
| pr196_seq_GTG_415-394_R | A1/T1<br>B1/T1<br>C1/T1<br>D2/T1–T5 |
| pr197_seq_ACA_415-394_R | A1/T4<br>B1/T4<br>C1/T4 |
| pr206_seq_TGT_448-473_F | E/T0, E1/T1–T5<br>F/T0, F1/T1–T5<br>I/T0, I1/T1–T5 |
| pr207_seq_GTG_448-473_F | E2/T1–T5<br>F2/T1–T5<br>I2/T1–T5 |
| pr209_seq_TGT_589-565_R | E/T0<br>E1/T1–T5<br>I/T0, I1/T1–T5 |
| pr210_seq_GTG_589-565_R | E2/T1–T5<br>I2/T1–T5 |
| pr215_seq_TGT_659-678_F | G/T0, G1/T1–T5<br>H/T0, H1/T1–T5 |
| pr216_seq_GTG_659-678_F | G2/T1–T5<br>H2/T1–T5 |
| pr212_seq_TGT_929-911_R | F/T0, F1/T1–T5<br>G/T0, G1/T1–T5<br>H/T0, H1/T1–T5 |
| pr213_seq_GTG_929-911_R | F2/T1–T5<br>G2/T1–T5 |

| | H2/T1–T5 |
|---|---|
| pr018_seq_TGT_518-540_F | 4-site replicate #1/T0–T6 |
| pr019_seq_GTG_518-540_F | 4-site replicate #2/T0–T6 |
| pr021_seq_TGT_709-688_R | 4-site replicate #1/T0–T6 |
| pr022_seq_GTG_709-688_R | 4-site replicate #2/T0–T6 |

**Table S12. Mean fractions of epistasis types across fitness quartiles.**

| | | mean fraction of epistasis type | | |
|---|---|---|---|---|
| quartile | epistasis type | TrpB | GB1 | null model |
| Q1 | magnitude | 0.56 | 0.53 | 0.71 |
| | reciprocal sign | 0.09 | 0.11 | 0.04 |
| | sign | 0.35 | 0.35 | 0.25 |
| Q2 | magnitude | 0.55 | 0.53 | 0.72 |
| | reciprocal sign | 0.11 | 0.12 | 0.04 |
| | sign | 0.34 | 0.34 | 0.24 |
| Q3 | magnitude | 0.60 | 0.62 | 0.79 |
| | reciprocal sign | 0.09 | 0.08 | 0.02 |
| | sign | 0.31 | 0.30 | 0.19 |
| Q4 | magnitude | 0.61 | 0.68 | 0.87 |
| | reciprocal sign | 0.08 | 0.05 | 0.00 |
| | sign | 0.31 | 0.28 | 0.12 |

**Table S13. Mean fractions of epistasis type across positions pairs.**

| | | mean fraction of epistasis type | | |
|---|---|---|---|---|
| positions | epistasis type | TrpB | GB1 | null model |
| 01 | magnitude | 0.58 | 0.69 | 0.73 |
| | reciprocal sign | 0.09 | 0.05 | 0.04 |
| | sign | 0.33 | 0.27 | 0.23 |
| 02 | magnitude | 0.60 | 0.52 | 0.68 |
| | reciprocal sign | 0.08 | 0.13 | 0.06 |
| | sign | 0.32 | 0.35 | 0.26 |
| 03 | magnitude | 0.56 | 0.61 | 0.75 |
| | reciprocal sign | 0.10 | 0.08 | 0.03 |
| | sign | 0.34 | 0.31 | 0.22 |
| 12 | magnitude | 0.60 | 0.65 | 0.72 |
| | reciprocal sign | 0.08 | 0.06 | 0.04 |
| | sign | 0.32 | 0.29 | 0.24 |
| 13 | magnitude | 0.60 | 0.71 | 0.80 |
| | reciprocal sign | 0.10 | 0.04 | 0.02 |
| | sign | 0.30 | 0.25 | 0.17 |
| 23 | magnitude | 0.51 | 0.42 | 0.74 |
| | reciprocal sign | 0.12 | 0.25 | 0.03 |
| | sign | 0.37 | 0.33 | 0.23 |
| overall | magnitude | 0.58 | 0.61 | 0.74 |
| | reciprocal sign | 0.09 | 0.10 | 0.04 |
| | sign | 0.33 | 0.30 | 0.22 |

**Table S14. P-values for the mean fraction of each epistasis type.**

| | **p-value of mean vs. null model** | |
|---|---|---|
| **epistasis type** | TrpB | GB1 |
| magnitude | $p < 1e-6$ | $p < 1e-6$ |
| sign | $p < 1e-6$ | $p < 1e-6$ |
| reciprocal sign | $p < 1e-6$ | $p < 1e-6$ |

**Table S15. Summary statistics for the directed evolution simulations.**

| metric | DE method | metric value for each landscape | | |
|---|---|---|---|---|
| | | TrpB | GB1 | null model* |
| **Mean maximum fitness achieved** | Method 1 | 0.53 | 0.47 | 1.11 |
| | Method 2 | 0.63 | 0.61 | 1.17 |
| | Method 3 | 0.71 | 0.67 | 1.20 |
| **Median maximum fitness achieved** | Method 1 | 0.56 | 0.46 | 1.23 |
| | Method 2 | 0.67 | 0.59 | 1.23 |
| | Method 3 | 0.74 | 0.69 | 1.23 |
| **Fraction of simulations reaching global maximum** | Method 1 | 0.02 | 0.01 | 0.64 |
| | Method 2 | 0.07 | 0.02 | 0.83 |
| | Method 3 | 0.15 | 0.03 | 0.96 |

*Because the null model was calculated additively from the single-mutants of the TrpB landscape, the maximum was 1.23 instead of 1.

**Table S16. $T_{50}$[a] values for selected variants.**

| ***variant*** | **$T_{50}$ (°C)** |
|---|---|
| *Tm*9D8* | 99.0 ± 0.6 |
| VIVS | 99.0 ± 0.8 |
| VIVG | >100 |
| VIKG | 92.8 ± 0.3 |
| AIKG | 91.0 ± 0.3 |
| CLKG | 90.6 ± 0.3 |
| ALKG | 90.1 ± 0.4 |
| CIKG | 92.3 ± 0.3 |
| VLKG | 93.2 ± 0.4 |

[a]$T_{50}$ is defined as the temperature at which a 1 h incubation causes a fitness reduction of 50% as compared to a room temperature incubation.

**Table S17. Data collection and refinement statistics for the structure of *Tm*9D8***

| **Structure** | ***Tm*9D8*** |
|---|---|
| **Unit cell** | |
| Space group | *I*4 |
| a, b, c (Å) | 165.7, 165.7, 83.06 |
| α, β, γ (°) | 90.0, 90.0, 90.0 |
| **Data collection** | |
| Wavelength (Å) | 0.97946 |
| Resolution (Å) | 45.99 – 2.15 |
| Total/unique no. of reflections | 825756/61153 |
| $R_{merge}$[a,b] | 0.17 (2.19) |
| $R_{p.i.m.}$[a,c] | 0.05 (0.64) |
| $CC_{1/2}$[a,d] | 0.99 (0.60) |
| I/σ(I)[a] | 13.2 (1.7) |
| Redundancy[a] | 13.5 (12.4) |
| Completeness[a] (%) | 99.9 (99.4) |
| **Refinement** | |
| No. of reflections used in refinement/test set | 61110/6021 |
| $R_{work}$[a,e] | 0.214 (0.303) |
| $R_{free}$[a,e] | 0.237 (0.326) |
| No. of nonhydrogen atoms | |
| protein | 5814 |
| ligand | 49 |
| solvent | 96 |
| root-mean-square deviation from ideal geometry | |
| bonds (Å) | 0.002 |
| angles (°) | 0.49 |
| Ramachandran plot[f] (%) | |
| favored | 97.24 |
| allowed | 2.37 |
| disallowed | 0.39 |
| PDB accession code | N/A |

[a]Values in parentheses refer to data in the highest shell.

[b]$R_{merge} = \sum_{hkl}\sum_i|I_{i,hkl} - \langle I\rangle_{hkl}|/\sum_{hkl}\sum_i I_{i,hkl}$, where $\langle I\rangle_{hkl}$ is the average intensity calculated for reflection *hkl* from replicate measurements.

[c]$R_{p.i.m.} = (\sum_{hkl}(1/(N-1))^{1/2}\sum_i|I_{i,hkl} - \langle I\rangle_{hkl}|)/\sum_{hkl}\sum_i I_{i,hkl}$, where $\langle I\rangle_{hkl}$ is the average intensity calculated for reflection *hkl* from replicate measurements and N is the number of reflections.

[d]Pearson correlation coefficient between random half-datasets.

[e]$R_{work} = \sum||F_o| - |F_c||/\sum|F_o|$ for reflections contained in the working set. $|F_o|$ and $|F_c|$ are the observed and calculated structure factor amplitudes, respectively. $R_{free}$ is calculated using the same expression for reflections contained in the test set held aside during refinement.

[f]Calculated with PROCHECK.
